# A repair-associated bronchial epithelial differentiation trajectory through *KRT14*+ basal and hillock-like cells drives airway inflammation and remodelling in childhood-onset asthma

**DOI:** 10.64898/2026.09.16.752044

**Authors:** Tessa E. Gillett, Aurore C.A. Gay, Jelmer R. Vlasma, Alexandra B. Firsova, Martin B. Banchero, Anna K. Renner, Amanda J. Oliver, Marijn Berg, Sjors Maassen, Leonie Apperloo, Bao-Han Ly, Djoke van Gosliga, Marnix R. Jonker, Waradon Sungnak, Orestes A. Carpaij, Tessa M. Kole, Laura Hesse, Sharon Brouwer, Petra L. van der Velde, Marissa Wisman, Mieke C. Zwager, Akshaya K. Saikumar Jayalatha, Bas G. Doddema, Frederique Alleblas, Putri A. Fajar, A. Alexandros Imprachim, Rosalie C. van Hulst, Marjan Luinge, Monique E. Lodewijk, Janna Bakker, Markus Weckmann, Corry-Anke Brandsma, Wim Timens, Judith Vonk, Sarah A. Teichmann, Christos Samakovlis, Kerstin B. Meyer, Gerard H. Koppelman, Maarten van den Berge, Martijn C. Nawijn

## Abstract

The bronchial epithelium in asthma is vulnerable to damage and has impaired barrier function, but the mechanisms by which it contributes to airway inflammation and remodelling remain unclear. Here, we dissect these epithelial and immunological disease mechanisms by establishing a comprehensive single cell atlas of the bronchial wall from 21 patients with childhood-onset asthma and 25 matched healthy controls. We identify a novel asthma-associated non-canonical epithelial differentiation trajectory in which a repair-associated *KLF4+* basal cell subset differentiates into *KRT13+* hillock-like cells through a proliferative *KRT14*+ intermediate. *In vitro* cultured matched primary bronchial epithelial cells show that this trajectory is retained in absence of exogenous factors. We find that IL-13 induces hillock-like cell differentiation into *CEACAM5*^hi^ goblet cells, driving goblet cell metaplasia. *Repair-associated* basal cells and transitioning *CEACAM5*^hi^ hillock-like cells strongly contribute to airway inflammation and remodelling. In turn, dendritic cells and mast cells promote a state of highly active epithelial differentiation, which shows increased multiciliated cell fate decisions, in concordance with an increase in multiciliated cell death observed in asthma. Proportions of the epithelial cells of the non-canonical differentiation trajectory are associated with clinical outcomes such as disease severity, FeNO, and small airway function.

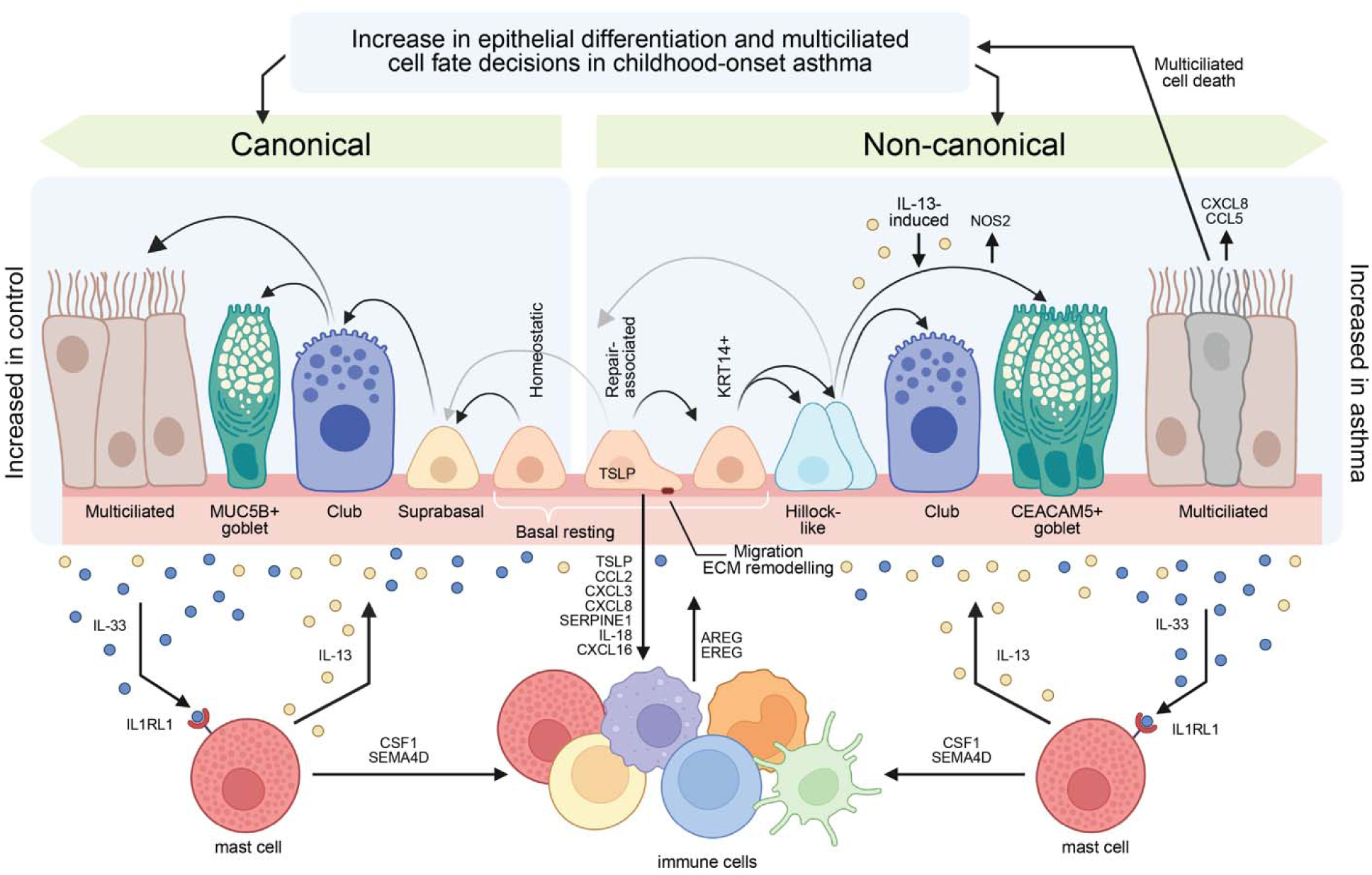

## Introduction

Asthma is a complex and highly prevalent obstructive respiratory disease characterized by chronic inflammation and remodelling of the airways, which mostly starts in childhood^1–3^. Childhood-onset asthma is often marked by type-2 cytokine activity and basement membrane thickening, and accompanied by allergic symptoms and other atopic disease^4^. Genetic and experimental studies^5–7^ have revealed an important role for the airway epithelium, which in asthma is characterized by goblet cell hyperplasia and metaplasia and a decreased barrier integrity^8^. Analysis of bronchial biopsies has shown reduced expression of adherens junction and tight junction proteins in the epithelium of patients with asthma^9,10^, and cultured primary bronchial epithelial cells (PBECs) from these patients have lower barrier function compared to PBECs from healthy controls^9^.

Additionally, the airway epithelium in asthma is vulnerable to damage, which drives both innate and adaptive type-2 immune activation through the release of alarmins such as IL-25, IL-33 and TSLP^1,3,11^. Biologics targeting the type-2 cytokines IL-4, IL-5 and IL-13 or their receptors are now used to treat patients with severe asthma, and novel therapeutics targeting IL-33 and TSLP are in clinical development, but response to these biologics is variable^12^.

Single cell sequencing studies have resulted in an improved understanding of the airway epithelial landscape^7,13^, and characterized unique features of the bronchial epithelium and its interaction with the immune system in patients with asthma both at baseline^13^ and after allergen challenge^7^, and provided spatial context^14^. These studies have revealed chronic IL-13 driven inflammation in asthma accompanied by a loss of cell-cell communication between the structural cells of the airways^7,13^. An increased transcriptional activity of genes involved in goblet metaplasia and extracellular matrix (ECM) remodelling was observed upon allergen challenge, as well as epithelial-immune cell crosstalk involved in a sustained immune response and remodelling^7^. However, these studies were hampered by a small number of biological replicates, limiting the generalizability of the results. Moreover, it remains unclear which cell-intrinsic features of the airway epithelium in patients with asthma cause its increased vulnerability and maintain chronic airway inflammation and remodelling in patients with childhood-onset asthma. We established a comprehensive single cell atlas of the airway wall in 21 patients with childhood-onset asthma and 25 healthy controls, aiming to perform detailed analysis of the molecular and cellular mechanisms underlying chronic airway inflammation and remodelling. We characterize the bronchial epithelium and its interaction with immune cells in relation to clinical features of asthma in a cohort with extensive clinical characterisation, and perform validation of key findings by *in vitro* primary cultures and spatial transcriptomic analysis on bronchial biopsies. We demonstrate that bronchial epithelial cell differentiation can proceed along an alternative trajectory that is intrinsically increased in patients with childhood-onset asthma, and originates from a repair-associated *KLF4*+ basal resting subset that transitions via KRT14+ basal cells to hillock-like cells. In asthma, hillock-like cells preferentially differentiate into goblet cells due to IL-13 exposure. Finally, we identify increased multiciliated cell death as a likely driver of the disproportionate differentiation activity in the asthmatic airway epithelium, and illustrate how these changes can contribute to airway remodelling and a pro-inflammatory feedback loop through epithelial-immune crosstalk.

## Results

### The childhood-onset asthma airway cell atlas

To study the molecular and cellular mechanisms that drive chronic inflammation and remodelling in the lower airways in asthma, we obtained bronchial biopsies from 21 patients with childhood-onset asthma and 25 age- and sex-matched healthy controls from the ARMS cohort^15^ (clinical trial number NCT03141814). We generated single-cell suspensions of 4-6 bronchial biopsies per study participant, which were processed for scRNA-seq analysis to establish an airway cell atlas of childhood-onset asthma. Asthma diagnosis was confirmed through measurement of the reversibility of airway obstruction following inhalation of salbutamol, or a positive provocation test for airway hyperresponsiveness (AHR) to methacholine. Patients with asthma were included if they had clinically stable disease without exacerbation, and no symptoms of respiratory infection for at least 6 weeks prior to bronchoscopy. Childhood onset of disease was confirmed through a documented asthma diagnosis before the age of 20. Participants did not use inhaled corticosteroids for at least 6 weeks prior to bronchoscopy. Clinical characteristics of patients and controls can be found in Suppl. Table 1A.

We performed batch integration on our dataset using scANVI^16^ which had the best combination of conservation of biological signals and correction of technical batch effects in the single-cell integration benchmark (scIB)^17^ (Extended Fig. 1A.) We performed iterative subsetting, quality control and cell type annotation on the integrated dataset according to current best standards^18,19^, yielding a fully integrated cell atlas of 240,756 cells, and 41 distinct cell types (Fig. 1A, 1B, Extended Fig. 1B, 1C), annotated across four hierarchical levels of increasing specificity, following the consensus cell type identities in the Human Lung Cell Atlas (HLCA)^20^ (Suppl. Fig. 1). Overall, our airway cell atlas of childhood-onset asthma, or in short asthma cell atlas, consists of a majority of 67.3% of epithelial, and 26.8% immune, 3.45% endothelial and 2.46% stromal cells (Fig. 1C). The asthma cell atlas is freely accessible in an online interface as a resource for the community at http://griacdata.org/. In patients with asthma, we observe a relative increase in the number of epithelial cells from the airways and submucosal glands (SMG), accompanied by a relative decrease in the number of endothelial, lymphoid, fibroblast and smooth muscle cells (Fig. 1D). The low proportion of smooth muscle cells sampled from patients with asthma may be due to a biopsy sampling bias caused by remodelling or increased stiffness of the airway wall, in conjunction with an increase in both the number of cells and thickness of the airway epithelium^21^. An in-depth overview of the top 50 differentially predicted cell-cell interactions between all cell types in the bronchial airway wall in patients with asthma compared to healthy controls, organized per sender or receive cell type can be found online at https://github.com/Nawijn-Group-Bioinformatics/Asthma_Cell_Atlas with the overall top 50 of differential interactions provided in Suppl. Fig. 2.

**Figure 1.**
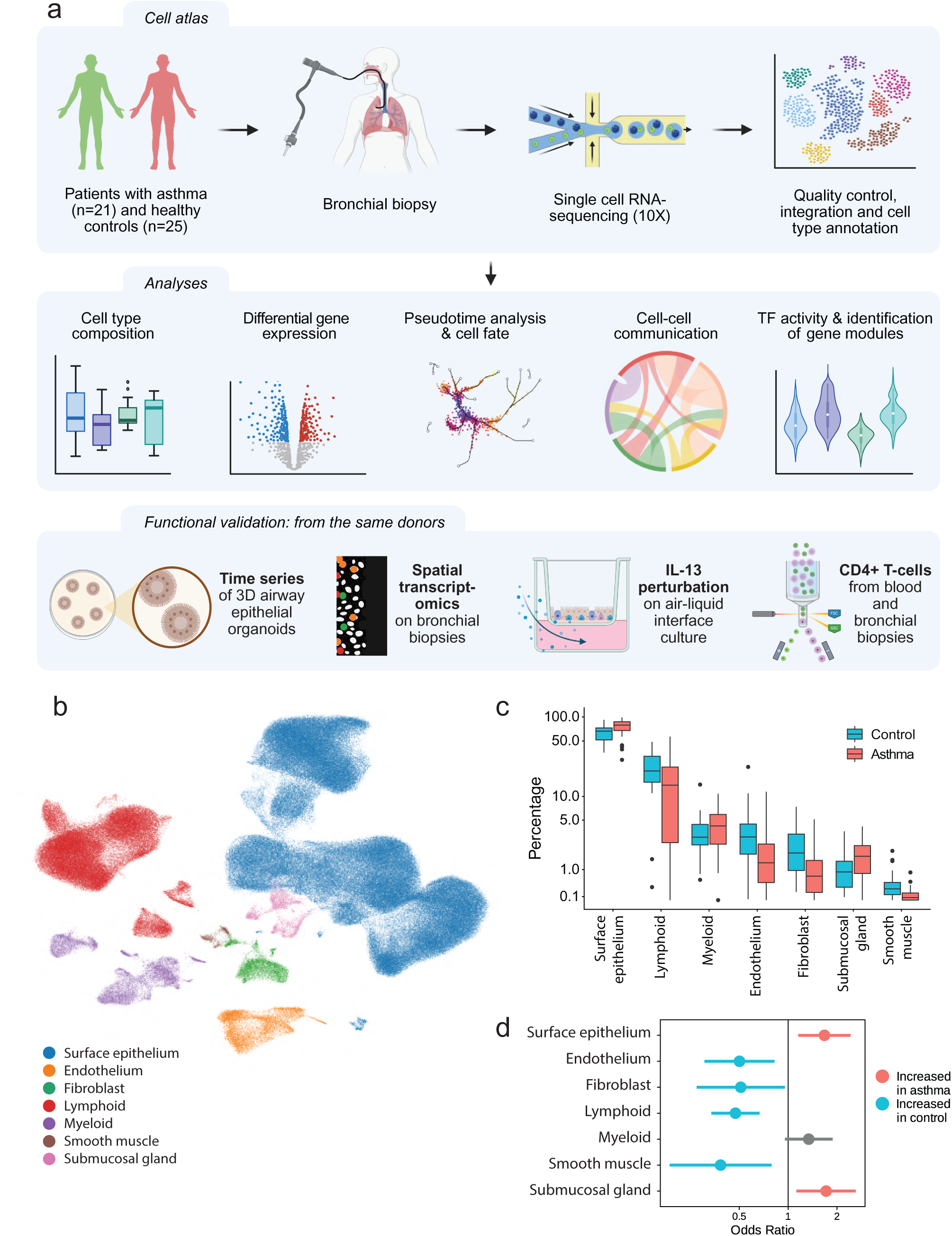
Generation and analysis of the Airway Cell Atlas of Childhood-onset Asthma. **a)** Study outline: scRNA-seq data generation from bronchial biopsies of patients with childhood-onset asthma and controls, including spatial transcriptomics; from 3D epithelial organoid and ALI cultures of matched primary bronchial epithelial cells; and from FACS sorted CD4+ T-cells from peripheral blood and bronchial biopsies. **b)** UMAP of the asthma cell atlas of patients with asthma and healthy controls. **c)** Cell type percentage per donor for major cell types in the asthma cell atlas in healthy controls (blue) and patients with asthma (red). **d)** Odds ratios of the cell type abundance in asthma compared to control as analysed in a logistic mixed effects model. Significant association with asthma or control in red or blue, respectively. Note that panel **c** summarizes donor-level percentages, whereas panel d reports results of a mixed effects model at the single cell level.

### The airway wall of patients with asthma is characterized by an increase in hillock-like and hypersecretory goblet cells

We identified 12 distinct subsets of surface airway epithelial cells, including basal (*KRT5*, *S100A2*), secretory (*SCGB1A1*, *BPIFB1*), and multiciliated (*FOXJ1, CAPS*) lineages and their subsets, as well as the rare epithelial cell types pulmonary ionocytes *(FOXI1, CFTR)^22^*, neuroendocrine cells *(SLC6A17)*, and tuft-like cells (*FOXI1*, *DAB1*), which were recently shown to contain both mature tuft cells and tuft-ionocyte progenitor cells^23^ (Fig. 2A, Extended. Fig. 2A, 2B). In addition, we observe a discrete epithelial cell subset expressing *KRT6A*, *KRT4*, and *KRT13*, which has been described previously as KRT13 suprabasal cells^24^. In agreement with the HLCA^20^ and previous publications^25^ we annotate this subset as hillock-like cells to distinguish them from the recently described specialized epithelial cells of the anatomically distinct hillock structures^22,26^. We identify a higher proportion of hillock-like cells in patients with asthma compared to healthy controls (Fig. 2B, 2C). We do not observe differences in the proportions of the other surface or SMG epithelial cell types within the surface epithelium and SMG subsets, respectively, between patients with asthma and controls, including goblet cells, annotated as the *MUC5AC/MUC5B/CEACAM5*-expressing subset of secretory epithelial cells (Fig. 2B, 2C, Suppl. Fig. 3). Cell type specific differential gene expression (DGE) analysis revealed a common IL-13 response in the surface epithelial cell subsets, including well-known IL-13 response genes^27^ such as *ALOX15*, *CCL26*, *CDH26*, *NOS2*, *MUC5AC*, *POSTN, SERPINB2* and *CST1*, in agreement with previous reports^7,13,28^ (Fig. 2D, Extended Fig. 2C, Suppl. Tables 2A-M). In the basal cell subsets, we additionally observe a wider range of differentially expressed genes, with lower expression of *HIF3A*, a transcriptionally induced regulator of hypoxia response, and *ZNF667* and higher expression of *KCNE3*, a voltage-gated potassium channel subunit, the centrosome protein *CEP72* and the hyaluronate receptor *CD44* in asthma, in agreement with earlier observations^29^. DGE and abundance analyses for endothelial and stromal cell subsets did not yield any transcriptional differences, while airway smooth muscle cells were decreased in proportion within the stromal lineage, and arterial endothelium cells were increased in proportion within the endothelial lineage in patients with asthma (Suppl. Fig. 3, Suppl. Tables 3A-C, Suppl. Tables 4A-E).

**Figure 2.**
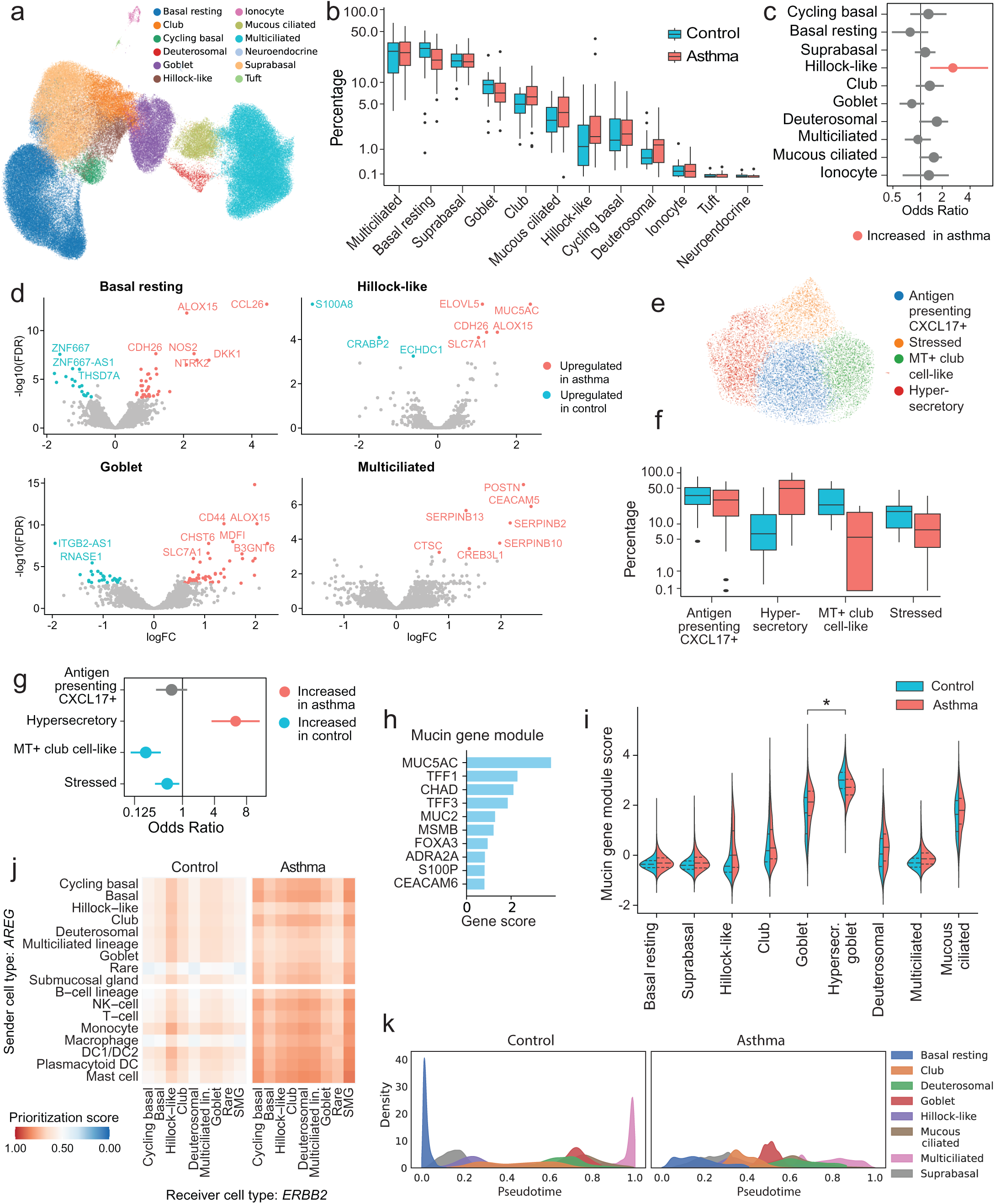
The airway epithelium in asthma has higher differentiation activity and is enriched for hillock-like cells and hypersecretory goblet cells. **a)** UMAP of the surface airway epithelial cells of the asthma cell atlas. **b)** Cell type proportions as percentage of total surface airway epithelium per donor. **c)** Odds ratios of the cell type abundance within the surface airway epithelium in patients with asthma compared to healthy controls (logistic mixed effects model). **d)** Log_10_(fold-change) versus -log_10_(FDR-corrected p-value) of differential gene expression in basal resting, hillock-like, goblet and multiciliated cells between patients with asthma and healthy controls. **e)** UMAP of the goblet cell population, showing four transcriptional phenotypes. **f)** Proportions of these subsets as percentage of all goblet cells per donor. **g)** Odds ratios of the abundance of the subsets within the total goblet cell population in patients with asthma compared to healthy controls (logistic mixed effects model). Note that panels **b** and **f** summarize donor-level percentages, whereas panels **c** and **g** report results of a mixed effects model at the single cell level. **h)** Top 10 gene contribution scores in a mucin production gene expression module, identified using DRVI. **i)** Expression values of this gene module per cell type. Asterisk indicates significant difference (logistic mixed effects model). **j)** MultiNicheNet prioritization score for *AREG* (ligand) to *ERBB2* (receptor) cell-cell communication from epithelial and immune cells as senders to epithelial cell types as receivers in healthy controls (left) and patients with asthma (right). **k)** Distribution of airway epithelium cells across Palantir pseudotime for healthy controls (left) and patients with asthma (right), per cell type.

In contrast to our previous findings^13^, we did not observe a difference in the proportions of goblet cells (*MUC5AC*, *MUC5B, CEACAM5*) and mucous ciliated cells (*FOXJ1*, *MUC5AC*) in this comprehensive group of patients with asthma and healthy controls. Recently, a specific subset of goblet cells characterized by high ribosomal gene expression, termed ‘quiescent goblet’, was identified in bronchial brush samples from patients with asthma^7^. Therefore, we performed further subsetting of the goblet cells through data-driven identification of stable clusters using the Adjusted Rand Index, which rendered four transcriptional phenotypes (Fig. 2E, Extended Fig. 3A). These were annotated as: club-cell like, metallothionein-expressing (*SCGB1A1*, *SCGB3A1*, *SERPINB3*, several *MT* genes); antigen presenting CXCL17+ (*HLA-A/B/C*, *HLA-DR*, *CXCL17*); stressed goblet (*VMP1*, *EZR*, *DNAJA1/DNAJB1*, *HSPA8*, *CDKN1A*, *UBC*); and a ‘quiescent goblet’-like cluster with a high percentage of ribosomal gene expression, and low numbers of read counts and expressed genes (Extended Fig. 3B, 3C, Suppl. Tables 5A-E).

The proportion of the quiescent cluster was strongly increased in patients with asthma, whereas club cell-like MT+ and stressed goblet cell subsets were decreased (Fig. 2F, 2G). We applied DRVI, a disentanglement variational autoencoder^30^, to perform data-driven identification of interpretable gene expression programs in our atlas (Extended Fig. 4A), which revealed one latent factor composed of mucin genes, including *MUC5AC*, *MUC2* and transcription factors such as *FOXA3*, *TFF1* and *TFF3,* of which *MUC5AC* and *FOXA3* are known asthma genes^6^ (Fig. 2H). The ‘quiescent’ cluster of goblet cells showed increased activity for this mucus-production gene module (Fig. 2I, Suppl. Fig. 4A). The transcriptional features of this subset, i.e. low diversity of expressed genes and number of reads and high expression levels of mucin and ribosomal protein genes, reflect high protein production of a small set of mucin genes. Therefore, we annotated this cluster as hypersecretory goblet cells, and propose that it contains those secretory cells producing high amounts of mucus, which would be identified as goblet cells in immunohistochemical stainings. In line with this, PAS stainings show an increased proportion of goblet cells in the airways of these patients with asthma compared to healthy controls (Extended Fig. 5). Use of this data-driven mucin gene expression module enables improved annotation of goblet and club cell subsets, which are difficult to distinguish based on standard marker gene expression, as classic goblet cell identity is defined on histological features^31^ and the distinction between club and goblet cells is difficult to make based on transcriptomic data^32^. In conclusion, patients with asthma have increased proportions of hillock-like cells and hypersecretory, which we interpret as genuine, goblet cells.

### The airway epithelium is characterized by higher differentiation activity in asthma

Hillock-like epithelial cells are a transitional cell state involved in epithelial repair and may contribute to squamous differentiation of the airway epithelium^26^. To assess whether their increased proportion in patients with asthma corresponds to altered epithelial differentiation, we performed a Palantir pseudotime analysis^33^ on basal, secretory and multiciliated lineage cells (Fig. 2K). In samples from healthy controls, we observed a marked enrichment of cells with pseudotime values at the very start or the very end of the epithelial differentiation trajectory, indicating a high proportion of undifferentiated basal resting cells and fully differentiated multiciliated cells. In total, 51% of cells were assigned a pseudotime value outside of the 0.05-0.95 range of the trajectory in controls, reflecting a homeostatic epithelium. In contrast, the vast majority of epithelial cells in patients with asthma have intermediate pseudotime values, with less than 4% of cells outside the 0.05-0.95 range of the trajectory (Fig. 2K), indicating a significantly more active process of epithelial differentiation compared to healthy controls (Extended Fig. 2D). Analysis of cell-cell communication patterns using MultiNicheNet^34^ identified an increase in AREG and EREG signalling by the epithelial and immune cell subsets to the ERBB2 and ERBB3 receptors on the asthmatic epithelium, which could maintain proliferation and differentiation of the airway epithelium in asthma (Fig. 2J, Suppl. Fig. 5). EGFR pathway activity has previously been shown to be increased in patients with asthma, initially involving EGFR and EGF^35^, but later also involving alternative ligands, such as AREG^36^, and receptors such as ERBB3^37^. Both EGF and AREG were shown to induce bronchial epithelial proliferation and sensitize to IL-13-induced goblet cell metaplasia^36^. Conversely, reduced expression of ERBB2 in airway epithelial cells from patients with asthma was shown to constrain epithelial proliferation and repair^38^. In conclusion, we find that the bronchial epithelium in patients with asthma has an increased differentiation activity.

### An asthma-associated basal cell subtype is characterized by expression of Laminins, MMP10, MMP13 and TSLP

As we observed an active differentiation process in the asthmatic epithelium, originating from basal cells, we contrasted the gene modules identified by DRVI expressed in basal cells between patients with asthma and controls (Extended Fig. 4A). One gene module expressed in basal cells appeared to be strongly associated with early pseudotime in asthma, but less so in control (Fig. 3A, Extended Fig. 4B, 4C.). The top genes contributing to this DRVI latent factor 5 include laminins *LAMC2*, *LAMA3,* metalloproteases *MMP10*, *MMP13*, the dystonin gene *DST*, the cytokeratin *KRT17*, and the alarmin *TSLP* (Fig. 3B). *KRT17* expression was recently found to mark a subset of uncommitted basal cells with the capacity to initiate metaplastic differentiation and epithelial repair programs^39^, while the laminins *LAMC2* are *LAMA3* are core components the Laminin-332 glycoprotein structure in the basement membrane^40^. Airway basal cells are anchored to these laminins through the hemidesmosomes, which connect to the cytoskeleton through dystonin^41,42^. Increased expression of *LAMC2* and *LAMA3* is associated with epithelial migration ^43,44^. In addition, this latent factor includes expression of the alarmin *TSLP*, which induces the type-2 immunity upon epithelial damage^45^, as well as the metalloproteinases *MMP10* and *MMP13*, which are involved in ECM remodelling in several lung diseases^46–48^. Taken together, we interpret this latent factor as a gene expression module associated with an epithelial repair response.

**Figure 3.**
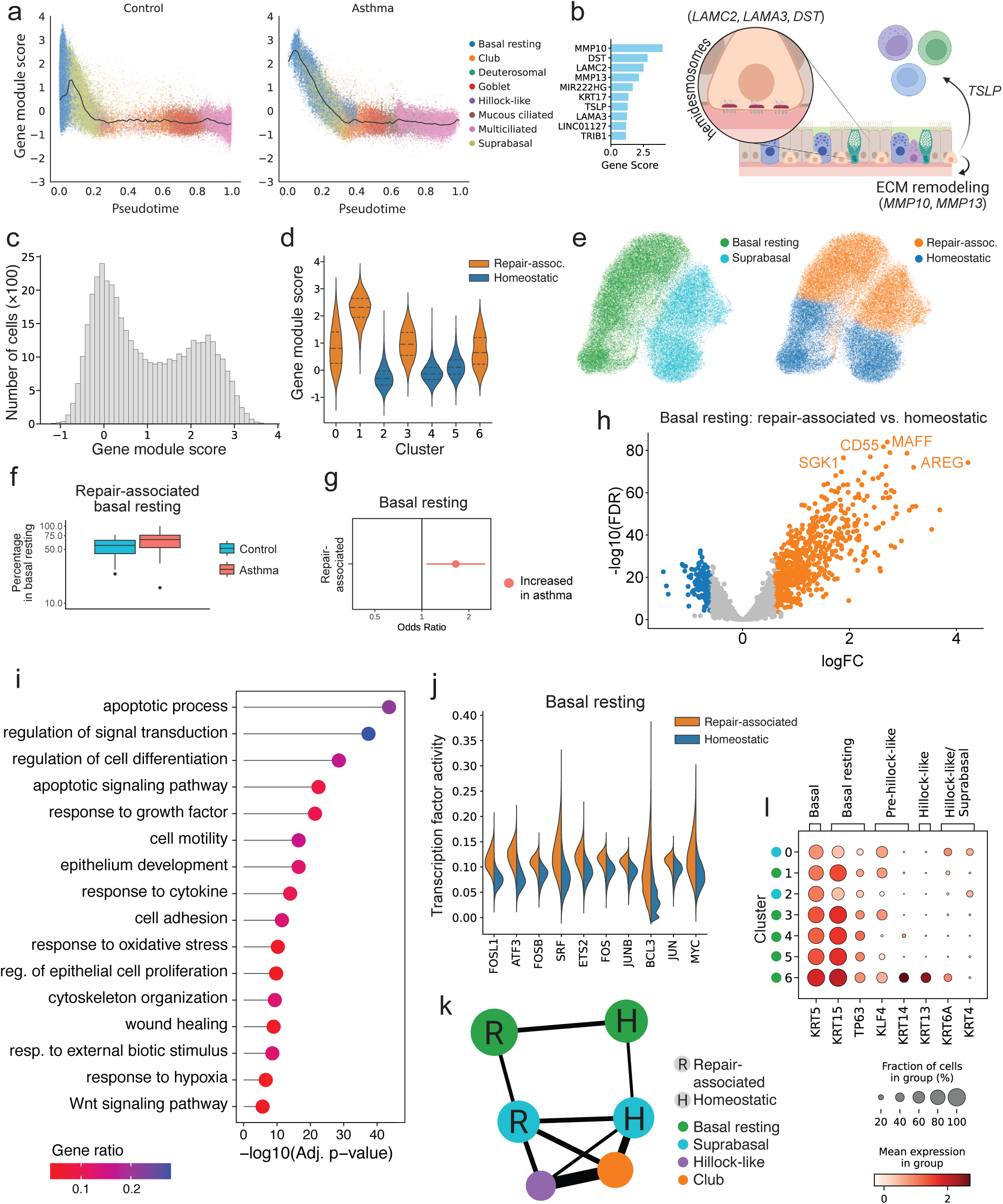
Repair-associated basal cells are one of two major basal cell subsets in the airways and are increased in patients with asthma. **a)** Airway epithelial repair gene module expression values across Palantir pseudotime, the black line shows the binned median value. **b)** Top 10 gene contribution scores in the repair gene module, reflecting ECM remodelling, hemidesmosomal anchoring to the basement membrane, and TSLP production. **c)** Distribution of repair gene module expression values in basal resting cells, and **d)** per basal cell subcluster, annotated as repair-associated basal (orange) and homeostatic basal (blue). **e)** UMAPs of the basal cell subset, displaying basal and suprabasal cells (left) versus repair-associated and homeostatic basal cells (right). **f)** Repair-associated cells as percentage of total basal resting cells per donor. **g)** Odds ratios of the abundance of repair-associated cells within the basal resting cell subset (logistic mixed effects model). Note that panel **f** summarizes donor-level percentages, whereas panel **g** reports results of a mixed effects model at the single cell level. **h)** Log_10_(fold-change) versus -log_10_(FDR-corrected p-value) of of differential gene expression between repair-associated and homeostatic basal resting cell subsets, corrected for disease status. **l)** Gene set enrichment analysis of DE genes between repair-associated and homeostatic basal resting cell subsets. **j)** Transcription factor activity per cell for repair-associated (orange) and homeostatic (blue) basal resting cells, using pySCENIC. **k)** PAGA graph of cell type connectivity, reflecting transcriptional similarity and/or transitions between cell types. Repair-associated and homeostatic basal subsets indicated with R and H, respectively; circle size reflects subset size. **l)** Marker gene expression in basal cell subclusters, green and blue markers indicate basal resting and suprabasal cell clusters, respectively.

We identified a bimodal distribution of the expression of this gene module within the basal cell subset of the asthma cell atlas (Fig. 3C, Suppl. Fig. 6A), in line with basal cell heterogeneity previously observed between the ventral versus dorsal location in the airways^39^. Therefore, we annotated basal resting and suprabasal cells with high activity of this gene module as ‘repair-associated’ and those without activity of the gene module as ‘homeostatic’ (Fig. 3D, 3E). While repair-associated basal resting cells are present both in patients with asthma and healthy controls, abundance analysis revealed that their proportion is increased in asthma compared to controls (Fig. 3F, 3G).

The repair-associated and homeostatic basal resting cells are transcriptionally highly distinct: DGE analysis with asthma as a covariate reveals a total of 3110 differentially expressed genes, including cytokeratins *KRT8* and *KRT17*, transcription factors such as *KLF4, FOSL1, FOSB, FOS, JUNB*, and *SOX9*, the EGFR ligand *AREG,* chemo/cytokines such as *CXCL8, CXCL3, CCL2* and *IL18,* the alarmin *TSLP,* the serine protease inhibitor *SERPINE1* (PAI-1), integrins *ITGA6*, *ITGB6* and *ITGB1*, laminins *LAMC1*, *LAMC2* and *LAMA3* and pro-survival factors such as *MYC*, *PIM1* and *MCL-1* (Fig. 3H, Suppl. Fig. 6B, Suppl. Table 6A-B). Expression of *AREG* in the repair-associated basal subset might contribute to the survival and proliferation signalling in the airway epithelium in asthma identified in Figure 2J. In line with these findings, enrichment analysis shows that these genes are associated with a wide range of biological processes, including epithelium development, regulation of epithelial cell proliferation, cell adhesion and cell migration (Fig. 3I, Suppl. Table 6C-D), indicating involvement of these cells in epithelial repair. Transcription factor activity analysis using pySCENIC^49^ confirms increased activity of the mentioned transcription factors in repair-associated compared to homeostatic basal resting cells (Fig. 3J, Suppl. Fig. 6C).

We next examined whether repair-associated basal resting cells are a transient cell state or a stable cell identity reflecting fundamental heterogeneity of basal cells within the airway epithelium. A PAGA graph shows strong transcriptional similarity between the repair-associated basal resting and suprabasal cells, as well as between homeostatic basal resting and suprabasal cells, without transitions across this transcriptional divide. These results suggest a lack of transitions between the repair-associated and homeostatic basal cell subsets (Fig. 3K).

We then performed DGE analysis within the repair-associated basal cells between patients with asthma and healthy controls (Extended Fig. 4D, Suppl. Tables 6E-H). In addition to a number of IL-13-induced genes also observed as differentially expressed in DGE analyses of all basal resting or suprabasal cells, we identified increased expression of *NTRK2,* encoding the TrkB receptor that can support anchorage-independent cell survival^50^ and enhances cholinergic neuroplasticity contributing to AHR in asthma^51^. We also found *LOXL4*, encoding a lysyl oxidase involved in ECM stabilization^52^, to be upregulated in repair-associated basal resting cells in patients with asthma, further supporting a role for this subset in wound repair processes in asthma specifically. Notably, these repair-associated basal cells display increased expression of the KLF4 transcription factor (Fig. 3L), which has been shown to drive basal to hillock-like cell differentiation in the context of squamous lung cancer^53^. In line with this, we observe a small subset of *KRT14*+ cells within the repair-associated basal cells, 1.7% of all basal resting cells. These *KRT14*+ basal cells retain expression of basal resting cell markers (*TP63*, *KRT15*), but also express low levels of early hillock-like marker genes (*KRT14*, *KRT6A*, *KRT13*) in addition to *KLF4* (Fig. 3L).

In conclusion, we have identified a repair-associated subset of basal resting cells that is higher in patients with asthma and that couples features of epithelial repair to airway inflammation (Extended Fig. 4E) and remodelling. Moreover, we observed increased expression of hillock-like differentiation factor KLF4 in this subset compared to homeostatic basal cells, and identified a subset of *KRT14*+ cells within the repair-associated basal cells that shows signs of differentiation into hillock-like cells.

### KRT14+ basal cells undergo cell-autonomous differentiation into hillock-like cells in patients with asthma

To study whether the increased proportion of repair-associated basal epithelial cells observed in the lower airways of patients with asthma represents a cell-autonomous effect, or is dependent on chronic inflammation or other exogenous effects, we generated time series scRNA-seq data from primary bronchial epithelial cells (PBECs) of our study participants cultured *ex vivo* in 3D spheroid cultures^54^ (Fig. 4A, Suppl. Table S1B). We selected 3D epithelial spheroid culture models as these are characterized by a lack of multiciliated cells or fully differentiated goblet cells, highlighting the less differentiated epithelial cell types and their transitions, including basal cells. We subsetted the basal cells into cycling cells, repair-associated basal cells (including a *KRT14*+ subset) and homeostatic basal cells, and additionally identified hillock-like cells, secretory cells and a small number of cycling secretory cells, as well as a cluster composed of the rare epithelial cell types, i.e. ionocytes, and tuft and neuroendocrine cells (Fig 4B, Extended Fig. 6A, 6B).

**Figure 4.**
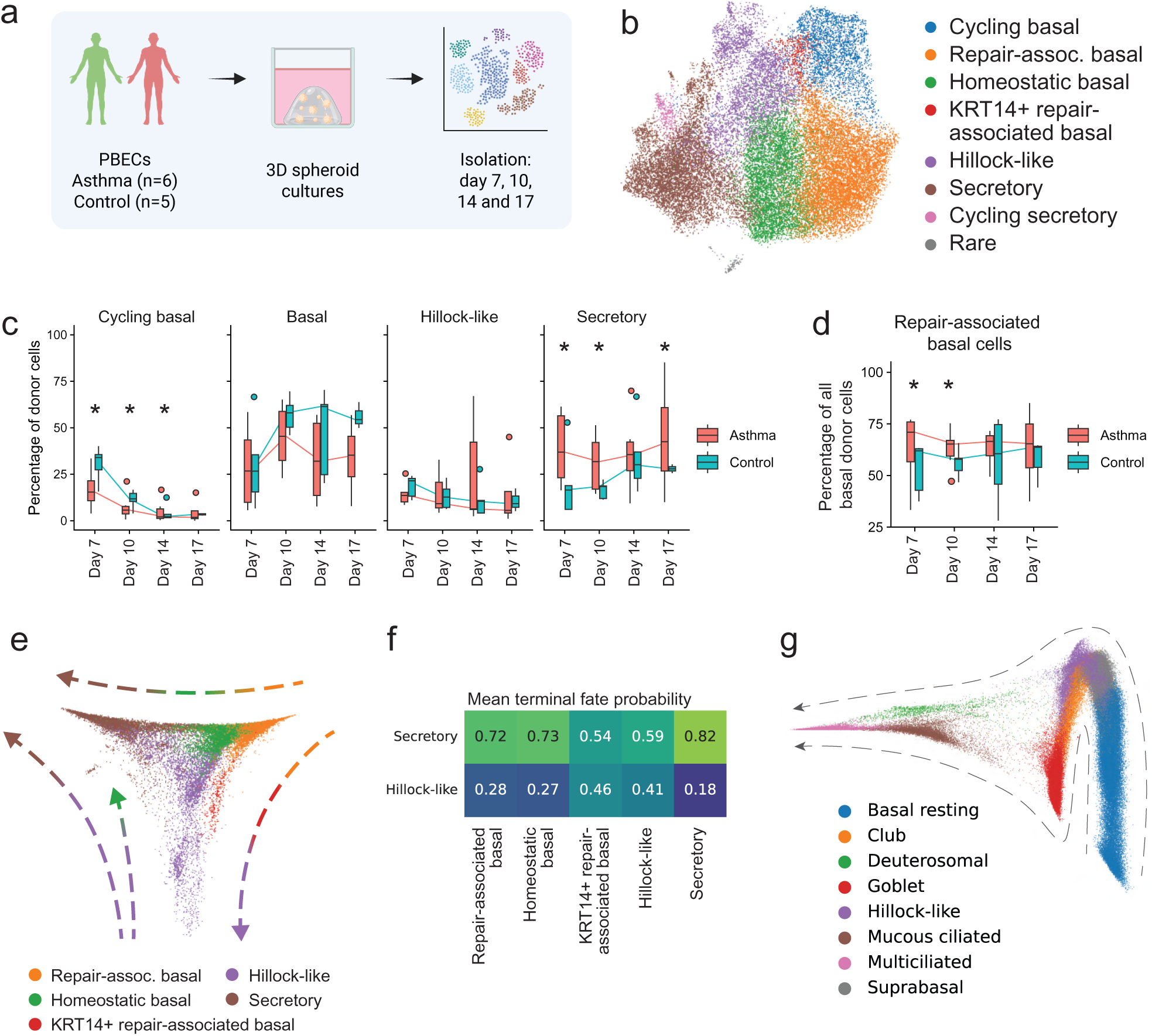
Repair-associated basal cell differentiation into hillock-like cells is a cell-autonomous trait in patients with asthma retained in *ex vivo* cultures. **a)** Experimental design of 3D epithelial spheroid culture time series experiment. **b)** UMAP of the primary bronchial epithelial cells from the spheroid cultures. **c)** Cell type percentage per sample and day, and **d)** proportion of repair-associated cells, including *KRT14*+ repair-associated cells, as percentage of the basal cell subset per sample and day. Asterisks indicate a significant difference between cultures derived from asthma (red) and control (blue) donors, logistic mixed effects model. **e)** Diffusion map showing a pseudotemporal ordering of the epithelial cells in 3D spheroid cultures, arrows indicate the observed transitions between cell subsets. **f)** Mean probability per cell subset of differentiation into either a secretory or hillock-like terminal fate (CellRank) in the 3D spheroid epithelial cell culture model. **g)** Diffusion map of the airway epithelial cells in the *in vivo* asthma cell atlas, arrows indicate observed transitions.

Time-resolved cell type abundance analysis shows an increased proportion of cycling basal cells in healthy controls compared to patients with asthma on day 7, 10 and 14, with cycling cell proportions decreasing over time in both asthma and control (Fig. 4C). The proportion of basal cells was increased on day 10, following the peak in proliferating basal cells on day 7, but was not significantly different between asthma and control. Within the subset of non-cycling basal cells we observed more repair-associated basal cells in 3D epithelial organoids derived from patients with asthma compared to those from healthy controls on days 7 and 10 (Fig. 4D), showing that this difference is retained in *ex vivo* cultures. We also observe a higher proportion of secretory cells in epithelial cell cultures from patients with asthma on days 7, 10, and 17 (Fig. 4C). DGE analyses on these cell types did not show transcriptomic differences between PBECs from patients with asthma compared to healthy controls with the exception of higher expression of *INHBA* in repair-associated basal cells from patients with asthma (Suppl. Tables 7A-I).

To study the underlying differentiation trajectories of these cells, we performed diffusion pseudotime analysis on the data from the 3D spheroid epithelial cultures. A data-driven diffusion map shows a pseudotemporal ordering of cells that, starting from basal cells, bifurcates into two branches of more differentiated cells: hillock-like cells and secretory cells respectively. A number of transitions can be observed in the diffusion map. Firstly, the subset of KRT14+ basal cells originating from the repair-associated basal cells are transitioning into hillock-like cells. Secondly, hillock-like cells display transitions towards both homeostatic basal cells and towards secretory cells. Finally, we also see direct transitions from both the repair-associated and homeostatic basal cells towards the secretory branch (Fig. 4E). Cell fate analysis using CellRank^55^ on the combined asthma and control PBEC dataset shows that the probability of a hillock-like terminal fate for *KRT14*+ basal cells is similar to that of hillock-like cells themselves, further supporting the *KRT14*+ to hillock-like cell transition (Fig. 4F). These data indicate that secretory cells in the airway epithelium can differentiate from either homeostatic basal cells, through suprabasal cells, or from repair-associated basal cells through *KRT14*+ basal cells and hillock-like cells.

To confirm the existence of this alternative basal to secretory cell differentiation trajectory through *KRT14*+ basal cells and hillock-like cells *in vivo*, we performed the same diffusion pseudotime analysis on the airway epithelial cells of the bronchial biopsy data of asthma cell atlas. This analysis shows that in the lower airways, basal resting cells differentiate into either a goblet or multiciliated endpoint, and that the hillock-like cells form a parallel route next to the canonical suprabasal-club trajectory (Fig. 4G). We do not observe large proportions of squamous cells in our cultured primary epithelial cells or in the asthma cell atlas. The data obtained from these 3D spheroid epithelial cultures also indicate that the increased proportion of repair-associated basal cells in patients with asthma is a cell-autonomous feature retained *ex vivo*, driven by either genetic susceptibility or epigenetic programming, and independent from ongoing airway inflammation *in situ*.

### Hillock-like cells are governed by two distinct transcription factor programs, and differentiate into basal, club, and goblet cells

We next sought to better characterize the hillock-like cell differentiation trajectory in the 4th-5th generation airways of patients with asthma and controls. Hillocks are morphologically defined, discrete aciliated structures present in the airways, which have a stratified epithelium. In contrast, pseudostratified *KRT13*+ epithelial cells with similar transcriptional programs have been described, but their function remains unclear^7,24,26^. Spatial transcriptomic analysis of bronchial biopsies from three patients with asthma and three healthy controls using SCRINSHOT^56^ shows that hillock-like cells, characterized by expression of *KRT6*, *KRT13A*, *KRT4* and/or *SPRR3*, commonly occur in both asthma-derived and healthy surface airway epithelium, but do not appear to form stratified hillock structures (Fig. 5A). Therefore, we describe this subset as hillock-like rather than hillock cells.

**Figure 5.**
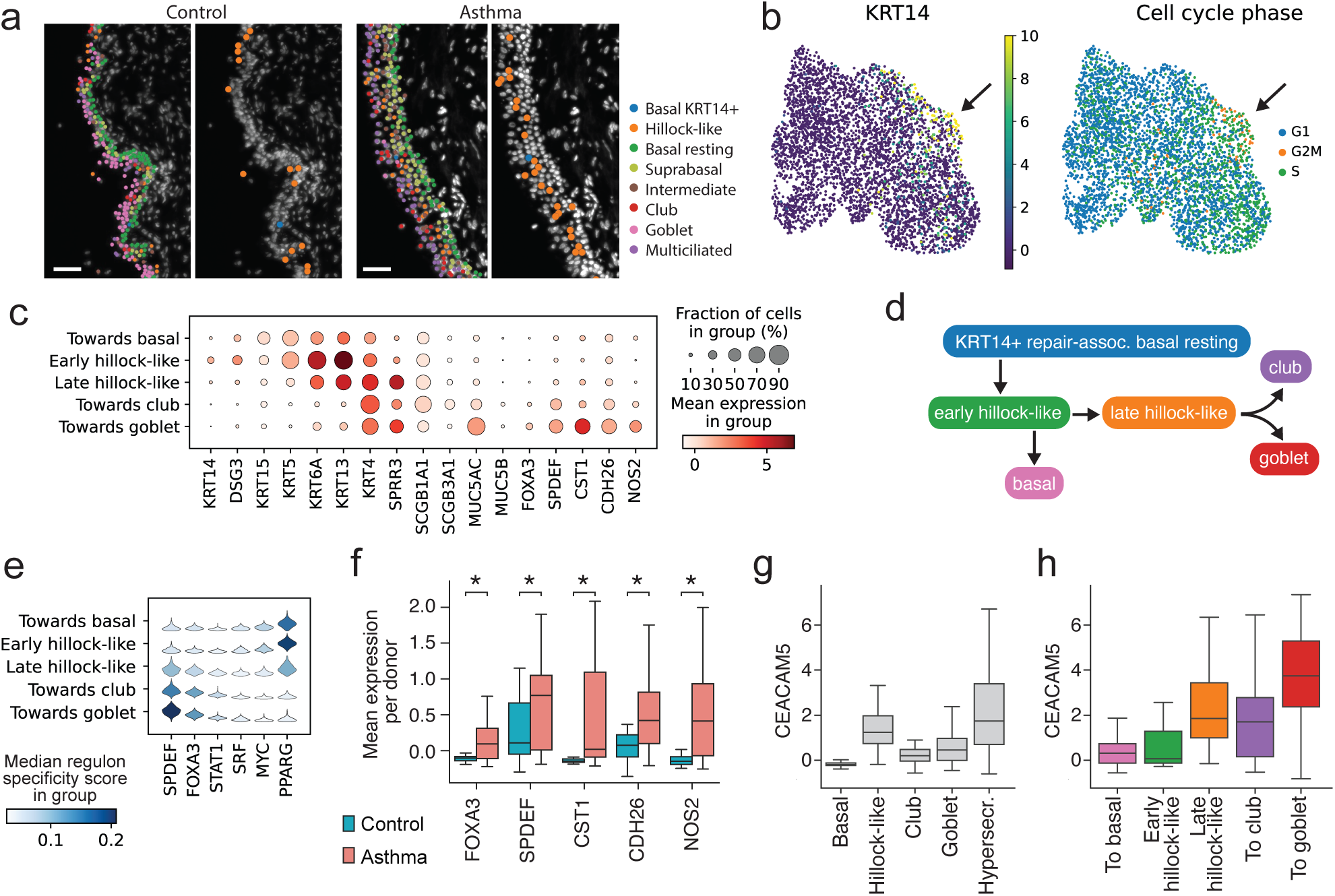
Hillock-like cells transition into basal and club cells, and into goblet cells in the presence of IL-13. **a)** Representative images of SCRINSHOT analyses identifying hillock-like and *KRT14*+ basal cells, alongside the other surface epithelial cell types, in bronchial biopsies of patients with asthma and healthy controls. **b)** UMAPs of the hillock-like cell subset in the asthma airway cell atlas, showing *KRT14* expression (left) and phase of cell cycle (right). Arrows indicate a subset of *KRT14*+ cells with increased cell cycle activity. **c)** Marker gene expression levels for five transcriptionally distinct subsets of hillock-like cells: two core hillock-like cell clusters and three clusters with expression indicative of differentiation to a basal, club or goblet cell identity. **d)** Proposed hillock-like cell differentiation model from *KRT14*+ basal resting cells. **e)** Transcription factor (TF) activity per hillock-like subgroup for *six TFs*, determined using pySCENIC. **f)** Mean gene expression level per donor of IL-13 response genes *FOXA3*, *SPDEF*, *CST1*, *CDH26* and *NOS2* within the hillock-like cells. **g)** Mean *CEACAM5* gene expression per donor in the main airway epithelial cell types, and **h)** in the five hillock-like cell subsets.

Further analysis of the hillock-like cell subset in the asthma cell atlas shows that hillock-like cells are transcriptionally heterogeneous. A small number of *KRT14*+ hillock-like cells show high cell cycle activity (Fig. 5B). Additionally, 26% of cycling basal cells express *KRT13* or *KRT14* (Suppl. Fig. 4B, 4C). This suggests a proliferative phase as *KRT14*+ basal cells transition to the hillock-like state, similar to the proliferation of hillock basal cells observed in stratified hillock structures^26^. We observe subsets of hillock-like cells with particularly strong expression of early hillock-like marker genes (*KRT6A*, *KRT13*) or expression of more mature hillock-like marker genes (*KRT4*, *SPRR3*); but also clusters of cells with basal (*KRT5*, *KRT15*), club (*SCGB3A1*), or goblet (*MUC5AC*) marker gene expression (Fig. 5C), suggestive of hillock-like cell differentiation towards those cell types (Fig. 5D). PAGA analysis of the hillock-like subset supports this interpretation (Suppl. Fig. 5E). Spatial transcriptomic analysis identified subsets of hillock-like cell populations with expression of *KRT6A*, *KRT13*, or *KRT4*, localised sparsely and predominantly in the intermediate layers of airway epithelium between basal and multiciliated cells, and occasionally in the SMG. In addition, *KRT14+* cells, while located mostly in the basal layer of the SMG duct, were also observed as individual cells in the surface airway epithelium (Extended. Fig. 7, Suppl. Fig. 7). Protein stainings for KRT5 and KRT14 showed that KRT14 expression was retained in large patches of the airway epithelium (Suppl Fig. 8), which were observed more often in the bronchial epithelium of patients with asthma than in controls pySCENIC^49^ analysis reveals that these subsets are governed by two distinct sets of transcription factors: early hillock-like cells and those differentiating towards a basal cell transcriptional profile show high activity of transcription factors PPARγ, MYC and SRF; whereas hillock-like cells with club or goblet cell markers show increased activity for transcription factors SPDEF, FOXA3 and STAT1 (Fig. 5E). The mature hillock-like subset has features of both. A transcription-factor-activity-based embedding of surface epithelium cell types clearly shows that transcription factor activity of these two subpopulations of hillock-like cells overlaps with that of suprabasal and secretory cells, respectively (Extended. Fig. 8A). While hillock-like cells differentiating towards goblet only show a trend towards an increased proportion (p=0.068) in patients with asthma (Extended Fig. 8B, 8C), we do observe increased expression of *FOXA3* and *SPDEF*, transcription factors associated with goblet cell differentiation, and *CST1*, *CDH26* and *NOS2* (Fig. 5F), which are well known IL-13 response genes^27^, in targeted comparisons of the expression of these genes using a mixed effects regression model. Expression of these genes is specifically high in the hillock-like subset differentiating towards a goblet cell identity, and within that subset remains increased in patients with asthma compared to controls (Extended. Fig. 8D).

**Figure 6.**
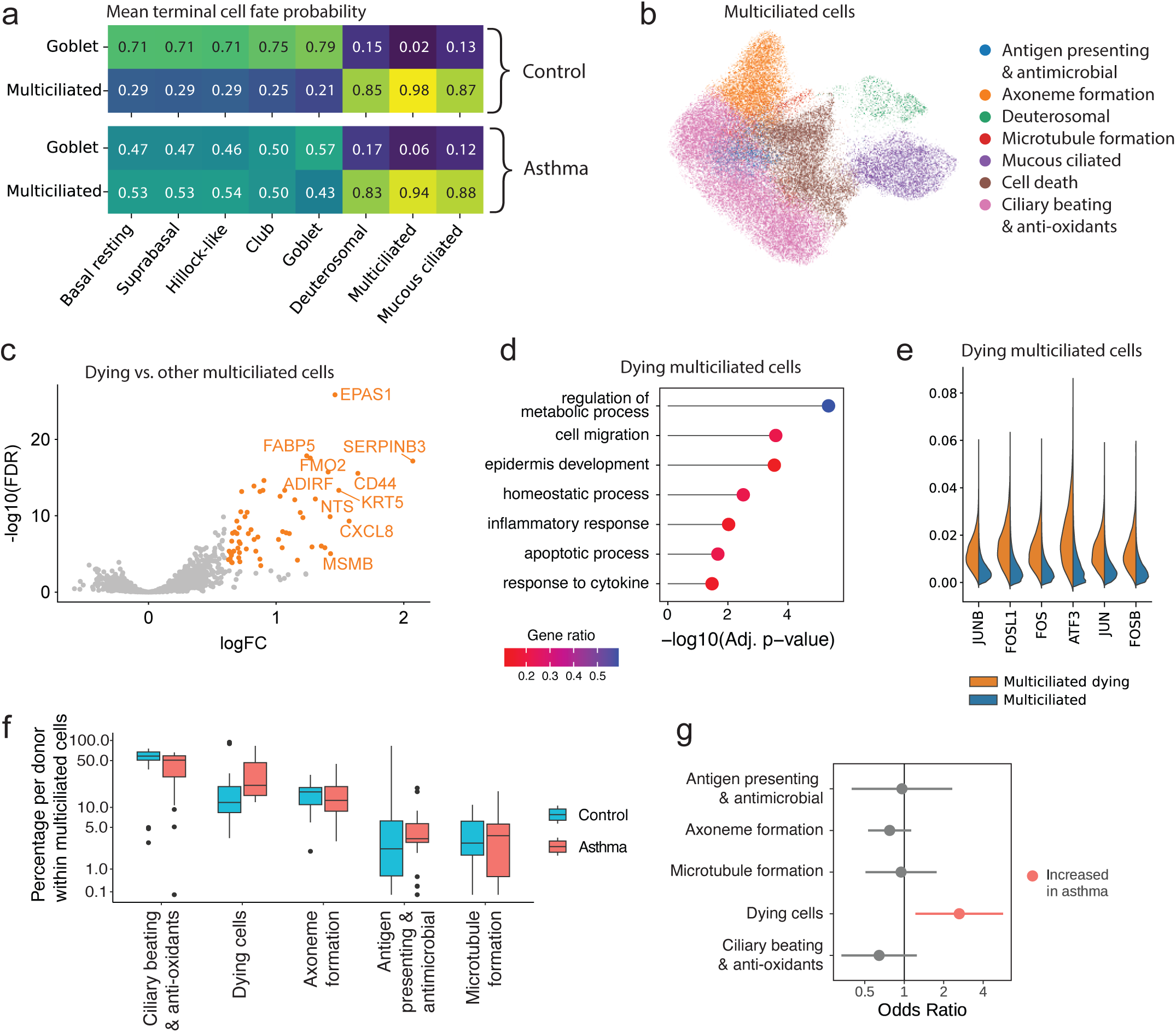
Increased multiciliated cell death in the airway epithelium in patients with asthma. **a)** Mean probability per cell typeof differentiation into either a goblet or multiciliated terminal cell fate (CellRank) in the asthma cell atlas. **b)** UMAP of the subsets of the multiciliated lineage cells, with 7 transcriptional phenotypes, including deuterosomal and mucous ciliated cells. **c)** Log_10_(fold-change) versus -log_10_(FDR-corrected p-value) of differential gene expression between dying multiciliated and all other multiciliated cells, corrected for disease status. **d)** Gene set enrichment analysis of the differentially expressed genes. **e)** Transcription factor activities per cell type, determined using pySCENIC. **f)** Multiciliated cell subsets as percentage of all multiciliated cells per donor. **g)** Odds ratios of the abundance of multiciliated cell subsets within the total multiciliated cell population (logistic mixed effects model). Note that panel **f** summarizes donor-level percentages, whereas panel **g** reports results of a mixed effects model at the single cell level.

**Figure 7.**
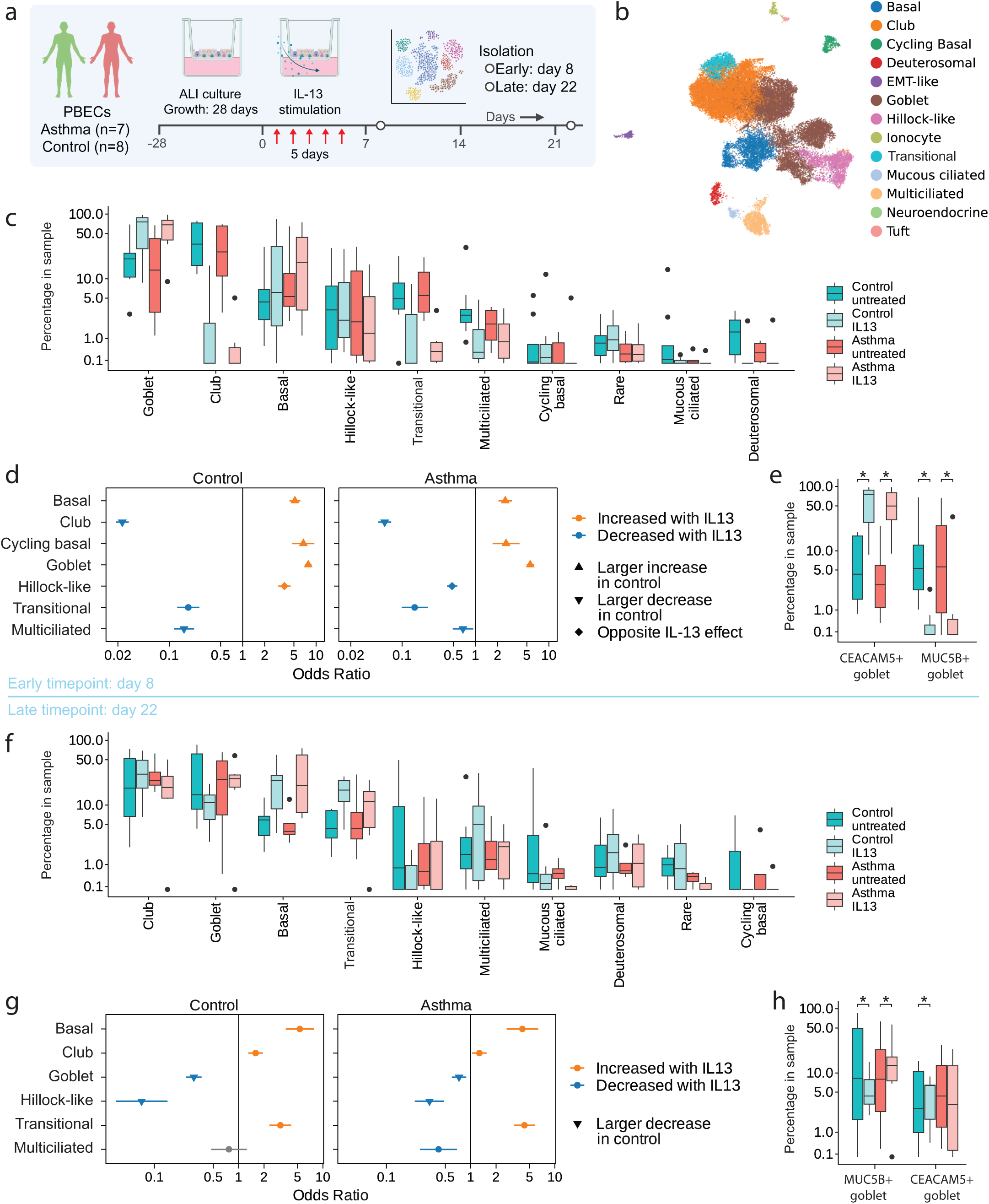
IL-13 induces short-term goblet cell hyperplasia and long-term loss of ciliated cells in ALI cultured PBECs from patients with asthma. **a)** Experimental design of IL-13 perturbation of matched primary bronchial epithelial cells grown in air-liquid-interface (ALI) cultures with short-term and long-term read-outs. **b)** UMAP of the ALI cultured cells. **c)** Cell type proportion as percentage of all cells per sample at 8 days after the start of IL-13 perturbation. **d)** Odds ratios of the cell type abundance at 8 days after IL-13 perturbation compared to untreated samples (logistic mixed effects model), for ALI cultures from control (left) and asthma (right) donors. **e)** Cell type proportion of CEACM5+ and MUC5B+ goblet cell subsets as percentage of all goblet cells per sample. Asterix indicates significant differences in cell type proportion between treatment conditions within a disease group (logistic mixed effects model). Panels **f)**, **g)** and **h)** show cell type proportions, odds ratios, and goblet cell subset proportions as in **c**, **d**, and **e**, now at 22 days after the start of IL-13 perturbation. Note that panel **c** and **f** summarize donor-level percentages, whereas panel **d** and **g** report results of a mixed effects model at the single cell level.

**Figure 8.**
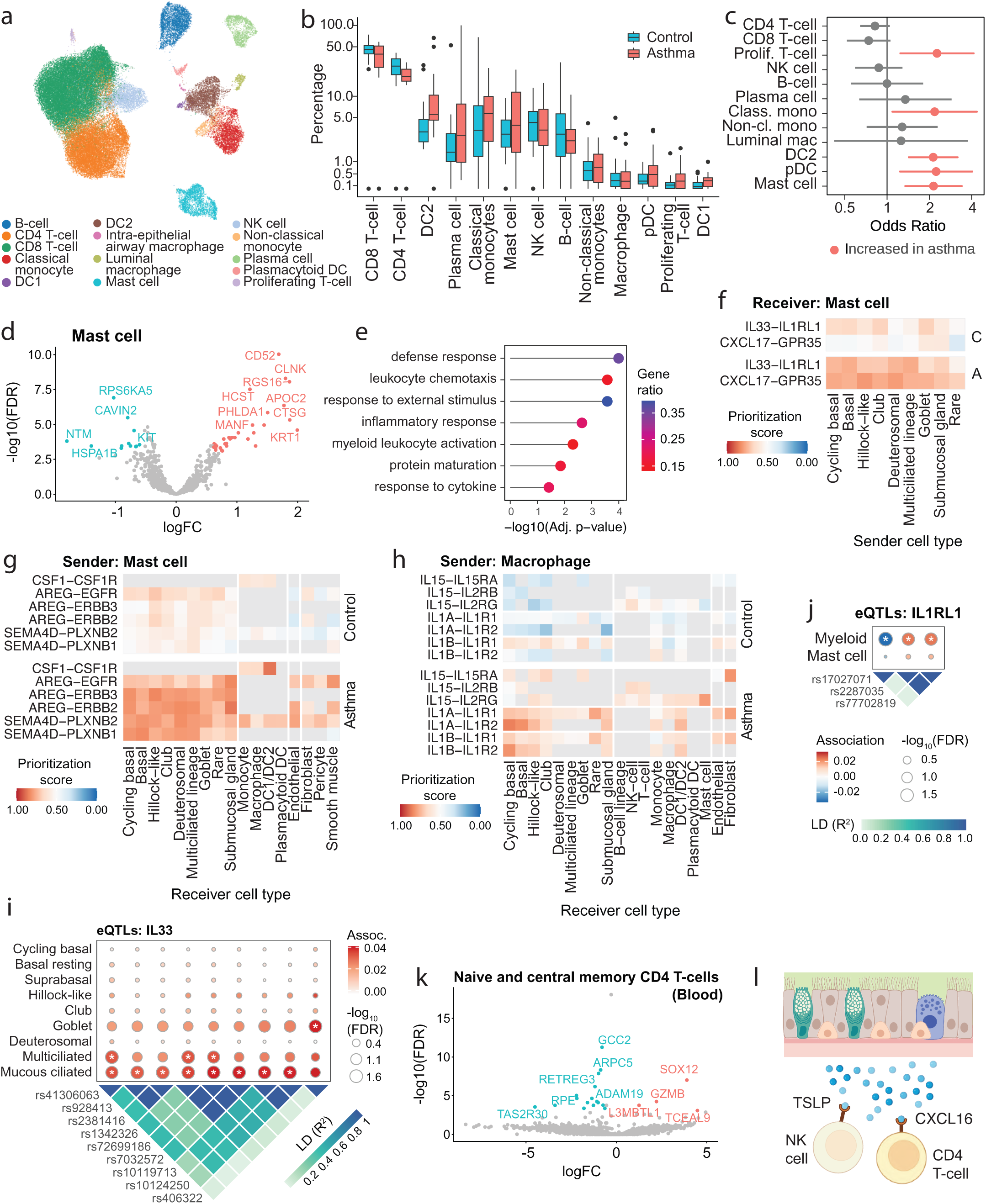
Activation of the immune system in the airway wall in patients with asthma. **a)** UMAP of the immune cell subset of the asthma cell atlas. **b)** Cell type proportions as percentage of all immune cells per donor. **c)** Odds ratios of the cell type abundance in patients with asthma compared to control (logistic mixed effects model). Note that panel **b** summarizes donor-level percentages, whereas panel **c** reports results of a mixed effects model at the single cell level. **d)** Log_10_(fold-change) versus -log_10_(FDR-corrected p-value) of differential gene expression in mast cells between patients with asthma and controls. **e)** Gene set enrichment analysis of the differentially expressed genes. **f)** MultiNicheNet prioritization score for cell-cell communication from epithelial cell types to mast cells in patients with asthma (A) and control (C), ligands and receptors indicated on the y-axis. **g)** MultiNicheNet prioritization score for cell-cell communication from mast cells, and **h)** from macrophages, to epithelial, immune, endothelial, and stromal cell types in patients with asthma and controls. **i)** eQTLs identified for IL-33 gene expression in goblet, multiciliated and mucous ciliated cells, and the linkage disequilibrium (LD) structure of the SNPs. All SNPs except rs406322 are in LD with asthma SNP rs992969 with an R^2^ between 0.46 and 0.97 in CEU ancestry. White asterisks indicate a significant eQTL effect. **j)** eQTLs identified for IL1LR1 gene expression in the myeloid cell subset, with SNP LD structure. **k)** Log_10_(fold-change) versus -log_10_(FDR-corrected p-value) of the differential gene expression between patients with asthma and controls in peripheral blood naïve and central memory CD4+ T-cells. **l)** MultiNicheNet cell-cell communication analysis reveals an increase in *CXCL16-CSCS6* interactions between epithelial cells and CD4+ T-cells in patients with asthma.

Finally, we observe high *CEACAM5* gene expression in both hillock-like cells and hypersecretory goblet cells, but not other goblet cells (Fig. 5G). Specifically, hypersecretory goblet cells from patients with asthma are *MUC5AC*+/*CEACAM5*+/*MUC5B*-, whereas those from healthy controls are *MUC5AC*+/*MUC5B*+ with lower *CEACAM5* expression (Extended. Fig. 8E). Within the hillock-like cells, *CEACAM5* expression is specifically high in the subset differentiating towards a goblet cell identity, suggesting that hillock-like cell differentiation contributes to the increase in hypersecretory goblet cells in patients with asthma (Fig. 5H), implicating the non-canonical differentiation trajectory in goblet cell hyperplasia In conclusion, we identify a small transitional subset of repair-associated, *KRT14*+ basal resting cells marked by cell cycle activity, that differentiates into hillock-like cells, connecting the increase in basal resting repair-associated cells in patients with asthma with the increased proportion of hillock-like cells. These hillock-like cells differentiate into basal, club and goblet cells, with the latter showing strong expression of IL-13 induced genes such as *NOS2* and *CDH26.* Expression of these genes may create a positive feedback loop that contributes to the disease process, as *NOS2* contributes to inducible nitric oxide production and airway obstruction^57^, and *CDH26* has been shown to enhance sensitivity of airway epithelial cells to IL-13-induced IL4R signalling^58^ and to contribute to amplification of type-2 inflammation in the airways of patients with asthma^59^.

### Highly active epithelial differentiation towards a multiciliated fate in asthma is associated with an ongoing loss of multiciliated cells

To further explore the altered differentiation trajectories of the airway epithelium in asthma, we performed cell fate analysis using goblet and multiciliated cells as terminal states, as confirmed by diffusion pseudotime (Fig. 4G). This revealed that in healthy donors, basal, hillock-like and club cells have an approximately 71%-75% probability of a goblet terminal fate. Goblet cells have a corresponding 79% probability of remaining a goblet cell, whereas the deuterosomal, mucous ciliated and multiciliated cells show a 83%-94% probability of a multiciliated terminal fate, reflecting the irreversibility of the multiciliated differentiation process (Fig. 6A)^60,61^. In patients with asthma, we observed an almost 2-fold higher probability of basal, suprabasal and club cells to adopt a multiciliated terminal fate compared to controls. The cell subsets of the multiciliated lineage show fate probabilities in patients with asthma that are comparable to those in healthy controls (Fig. 6A).

Despite the increased probability of a multiciliated cell fate (Fig. 6A) and the ongoing differentiation process in the airway epithelium of patients with asthma (Fig. 2K), we did not observe a higher proportion of multiciliated cells compared to controls (Fig. 2B), suggesting an ongoing loss of multiciliated cells in asthma. We therefore examined the transcriptional heterogeneity within the multiciliated cells, identifying five stable subsets (Extended Fig. 3D). Based on gene expression, four of these could be annotated as multiciliated cell subsets with a transcriptional phenotype of antigen presentation & antimicrobial activity; axoneme formation; microtubule formation; or ciliary beating & antioxidant activity (Fig. 6B, Extended Fig. 3F). The fifth subset showed a comparatively low number of expressed genes and reads, and a high percentage of mitochondrial reads, indicating low transcriptional activity often associated with dying cells (Extended Fig. 3E). DGE analysis on this subset reveals 144 significantly upregulated genes compared to the other multiciliated cells, including the chemokine *CXCL8* and *CCL5*, as well as cathepsin C (*CSTC*) (Fig. 6E, Suppl. Tables 8A-F). Enrichment analysis of the genes specific for this multiciliated cell subset identifies multiple biological processes, including apoptosis, migration, epidermis development and response to cytokine (Fig. 6C, Suppl. Table 8G). We hypothesize that this multiciliated cell subset consists of dying or apoptotic cells, with expression of pro-inflammatory chemokines. Additionally, transcription factor analysis showed an increase in activity of various AP-1 associated transcription factors, including JUNB, FOSL1, FOS, ATF3, JUN and FOSB (Fig. 6E). Abundance analysis revealed an increase of the number of dying multiciliated cells in asthma compared to healthy controls (Fig. 6F, 6G).

Due to the lack of fully differentiated cells in the 3D epithelial organoid time series data, these experiments can not be used to assess whether hillock-like cells differentiate into multiciliated cells. However, we do find that the gene signature of dying multiciliated cells is increased in repair-associated compared to homeostatic basal cells (Extended Fig. 3G), including *EPAS1* and *KLF4*, which are overexpressed in the dying multiciliated cells compared to the other multiciliated cell subsets, suggesting a potential lineage relationship with the repair-associated basal cells. We also found a strong correlation between the percentage of repair-associated basal resting cells and dying multiciliated cells per donor (Extended Fig. 3H); and observe that the small number of hillock-like cells that overlaps with the multiciliated cell population in the transcription-factor-activity-based embedding, predominantly clusters with the dying multiciliated cells (Suppl. Fig. 4F). The dying multiciliated cells have high expression of *CTNNB1* (beta-Catenin) and *TCF7*, key mediators of canonical Wnt signalling, which has been shown to drive hillock-like cell differentiation^62^ (Suppl. Table 8F). Taken together, these results show a higher proportion of dying multiciliated cells in the airway epithelium in patients with asthma, which could explain the highly active epithelial differentiation in these patients (Fig. 2K), and which contributes to airway inflammation through *CXCL8* and *CCL5* expression, recruiting neutrophils and eosinophils respectively.

### In vitro stimulation of PBECs with IL-13 results in loss of multiciliated cells, with decreased recovery in cultures from donors with asthma

Our data indicate that epithelial differentiation trajectories are altered, alongside a universal IL-13 response signature in the airway epithelium in asthma (Fig. 2D) consistent with chronic IL-13 activity *in vivo*. To further study the effect of IL-13 exposure on the epithelium of patients with asthma, we conducted an *ex vivo* IL-13 perturbation experiment in air-liquid-interface (ALI) cultured PBECs. ALI-differentiated PBECs from patients with asthma and healthy controls were treated with IL-13 for 5 days, and harvested for scRNA-seq at 3 and 17 days after the end of the IL-13 exposure (Fig. 7A, Suppl. Table S1C). We identified basal, cycling basal, hillock-like, club, goblet, deuterosomal, multiciliated and mucous ciliated cells, as well as neuroendocrine cells, tuft-like cells and ionocytes in these ALI cultures (Fig. 7B, Extended Fig. 9A, 9B). Additionally, we identified a subset of transitional epithelial cells and a cluster of cells undergoing epithelial-to-mesenchymal transition (EMT), neither of which correspond to any cell subset observed in the *in vivo* asthma cell atlas.

Unstimulated ALI cultures from PBECs of patients with asthma and controls showed similar cell type proportions, although we observed a higher proportion of deuterosomal cells in ALIs from controls (Extended Fig. 9C). Transepithelial electrical resistance (TEER) was measured each week of ALI culture. In line with previous reports^9,10^, we observed that barrier function was reduced in PBECs from patients with asthma when compared to controls. Furthermore, we found that IL-13 induced loss of barrier function in PBECs of patients with asthma, but not of healthy controls (Extended Fig. 9D). At the early timepoint, there was an IL-13 induced increase in basal, cycling basal, and goblet cells and a decrease in club, multiciliated and transitional cells in both asthma and control (Fig. 7C, 7D). These differences were stronger in control, which could reflect an altered sensitivity to IL-13 between PBECs from patients with asthma and healthy controls. Hillock-like cells are increased upon IL-13 stimulation in control PBECs, but decreased in asthma. The increase in goblet cells is due to a strong increase in proportion of a *MUC5AC*+/*CEACAM5*+ subset that resembles the hypersecretory goblet cells found *in vivo* in patients with asthma, whereas a subset of *MUC5AC*+/*MUC5B*+/*CEACAM5*^low^ goblet cells was decreased after IL-13 (Fig. 7E, Extended Fig 9C).

At the late timepoint, the increased proportion of basal cells in IL-13 treated PBECs persists, accompanied by an increase in club and transitional cells in both asthma and healthy controls (Fig. 7F, 7G, Extended Fig. 9D). Goblet and hillock-like cells were decreased in IL-13 stimulated cultures at the late timepoint, with a stronger decrease in control PBECs. This, together with the short-term response, suggests that the increase in hillock-like cells in asthma *in vivo* is not likely to be due to IL-13. The IL-13 induced decrease in multiciliated cell proportions observed at the early timepoint was no longer observed in ALI cultures from healthy controls at the late timepoint, whereas it was retained in ALI cultures from patients with asthma (Fig. 7G).

DGE analysis of IL-13 treated ALI cultures compared to untreated control cultures shows an early IL-13 response in basal and *CEACAM5*+ goblet cells in healthy control-derived PBECs, with a similar but more modest induction of these genes in the same cell types in PBECs from patients with asthma (Extended Fig. 9F, 9G, Suppl. Tables 9A-F). Notably, in ALI cultures from both asthma and controls we observed increased expression of *SPRR3* in IL-13 treated CEACAM5+ goblet cells, linking their increased abundance in the ALI cultures to hillock-like-to-goblet cell differentiation. At the late timepoint, we observed a strong, and relatively comparable, transcriptional response in club and *CEACAM5*+ goblet cells in ALI cultures from both patients with asthma and controls (Extended Fig. 9F, G, Suppl. Tables 9G-X) Three-day IL-13 treatment of ALI cultured PBECs has previously been shown to induce mislocalization of basal bodies and loss of cilia, repress FOXJ1 activity and induce MUC5AC expression in FOXJ1+ cells, independent of NOTCH activity^63,64^. We confirm that IL-13 stimulation resulted in a short-term loss of multiciliated cells in both asthma and control PBECs, but where healthy control derived ALI cultures recovered their multiciliated cell proportions, ALIs with PBECs from patients with asthma did not. This suggests that while we observe ongoing differentiation *in vivo* in patients with asthma, likely as a result of high multiciliated cell death, recovery of multiciliated cell proportions may be hampered by chronic IL-13 exposure. Finally, IL-13 stimulation of both asthma and control PBECs resulted in a marked short-term increase in specifically *CEACAM5*+ goblet cells, whose gene expression profile suggests an origin in the hillock-like epithelial differentiation pathway in the asthma cell atlas biopsies (Fig. 5H). Therefore, we propose that the IL-13 induced goblet cell metaplasia in patients with asthma, observed as an increase in hypersecretory goblet cells *in vivo* (Fig. 2H, 2I), mainly occurs through the non-canonical hillock-like trajectory.

### Mast cell-epithelial crosstalk is amplified in asthma, driving a pro-inflammatory feedback loop

We found that the airway epithelium in asthma displays a pro-inflammatory phenotype, with increased expression of chemokines and alarmins in several disease-associated epithelial cell subsets that are part of the non-canonical epithelial differentiation trajectory. We therefore also performed detailed analysis of the immune cells in the asthma cell atlas, with in total 64,476 cells annotated to 14 cell types (Fig. 8A, Extended Fig. 10A, 10B). In line with the presence of chronic airway inflammation, we observed a higher proportion of classical monocytes, type 2 dendritic cells (DC2s), plasmacytoid dendritic cells (pDCs), mast cells, and proliferating T-cells in patients with asthma (Fig. 8B, 8C). Previous studies by us and others revealed higher proportions of mast cells and B lymphocytes in the airways of patients with asthma^7,13^. We are the first to show increased proportions of pDCs in the airway wall in patients with asthma in the absence of exacerbations or allergen challenges. In previous studies, pDCs were shown to be increased in BAL or induced sputum of patients with asthma after allergen challenge^65,66^ and in induced sputum during exacerbations^67^. At baseline, DC2s were found to be increased in the airway epithelium in type-2 high but not in type-2 low asthma^68^. Upon allergen challenge, DC1s were shown to be higher in BAL^69^ and DC2s were higher in BAL and bronchial biopsies^69^ as well as in bronchial brushes^7^ of patients with asthma compared to controls. Overall, the increased proportions of classical monocytes and different DC subsets in the airway wall of patients with asthma at baseline indicates increased immune surveillance^70^, reflecting a primed immune system despite the absence of active triggers. Several factors produced by the epithelial cell subsets of the non-canonical trajectory could contribute to an increased number and activity of innate immune cells, such as *TSLP, CXCL8, CXCL3, CCL2* and *SERPINE1* expressed by the repair-associated basal cells and *CCL5* and again *CXCL8* expressed by the dying multiciliated cells.

In addition to increased numbers of dendritic cells, we mainly observe interactions between the airway epithelium and the mast cells and T-cell subsets. DGE analysis, followed by enrichment analysis, showed a clear activated phenotype of mast cells in patients with asthma compared to those of healthy controls (Fig. 8D, 8E, Suppl. Tables 10A-K). Cell-cell communication analysis predicted that mast cells from patients with asthma receive more IL-33 signalling, originating from the basal and multiciliated subsets, and more *CXCL17* signalling from all epithelial cell subsets (Fig. 8F). In turn, the mast cells interact with the epithelium by sending pro-inflammatory and remodelling-associated ligands, including AREG and SEMA4D (Fig. 8G), thereby contributing to the proliferation and survival signalling potentially sustaining epithelial differentiation (Fig. 2J). Mast cells also interact through CSF1 with monocytes, macrophages and DCs. This might contribute to the higher number of DCs as observed in our data, but also to pro-inflammatory signalling by macrophages through interleukins IL-1A, IL-1B and IL-15, which is increased in patients with asthma, and predicted to mainly act on the basal and rare epithelial cells, DCs, mast cells and fibroblasts (Fig. 8H). Together these results begin to chart a self-sustaining chronic inflammatory cascade in the airway wall in patients with asthma.

Interestingly, the IL-33-dependent interactions between the airway epithelium and mast cells are in part governed by asthma-associated genetic variants. Cell-type-specific cis-eQTL analysis reveals an higher expression of the *IL33* gene in mucous ciliated and multiciliated cells (Fig 8I), including a cis-eQTL effect of SNPs in linkage disequilibrium with the well-known asthma-associated SNP rs992969^71^. In addition, we observed a goblet-cell specific association of SNP rs406322 and *IL-33* gene expression (Fig. 8I). The expression levels of the *IL1RL1* gene, encoding one of the two chains of the heterodimeric IL-33 receptor^71^, are also associated with SNPs in LD with known asthma-associated genetic signals in myeloid cells including mast cells, although this association did not reach significance due to lack of power (Fig 8J).

In the asthma cell atlas, all T-cells have a comparatively low RNA content per cell, which hampers identification of subsets of CD4+ or CD8+ T-cells at high resolution. To examine CD4+ T-cell subsets in further detail, we generated a combined dataset of FACS-purified CD4+ T-cells obtained from both bronchial biopsies and peripheral blood of these same donors (Suppl. Table S1D), making use of the comparatively sensitive SmartSeq2 scRNA-seq platform. CD4 T-cells from 14 patients with asthma and 13 healthy controls were included. After quality control this resulted in 3,660 and 3,708 total cells from blood and biopsies, respectively. We were able to annotate these in more detail, yielding 6 CD4+ T-cell subsets labeled as naive/central memory, effector memory, CD45RA+ effector memory (EMRA, in blood), tissue resident memory (TRM, in biopsies), cytotoxic (in blood), and regulatory CD4 T-cells. The TRM subset was further divided into cycling, cytotoxic, and resting/effector subsets (Suppl. Fig. 9A-C). Based on marker gene expression, we were unable to distinguish naive from central memory CD4 T-cells.

Abundance analysis on this data identified a higher proportion of regulatory T-cells cells within the CD4+ T-cells of patients with asthma compared to healthy controls, as well as a trend towards a higher proportion of cytotoxic CD4+ T-cells in bronchial biopsies of patients with asthma (p=0.066) (Extended Fig. 10C-G). Presence of cytotoxic *GZMB*+ CD4+ T-cells has previously been described in BAL of patients with severe asthma^72^. Additionally, DGE analysis of the naive/CM subset in blood showed an increase in *GZMB* expression in patients with asthma, suggesting increased cytotoxic activity in central memory CD4 T-cells (Fig. 8K, Suppl. Tables 11A-I). Interestingly, cell-cell communication analysis of the asthma cell atlas also shows an increase in pro-inflammatory CXCL16 signalling from the epithelium towards T-cells in patients with asthma, particularly in CD4+ T-cells (Fig. 8I, Extended Fig. 10H). CXCL16/CXCR6 signalling has been shown to be critical to attract and retain tissue-resident memory T-cells to the airways^73^.

In contrast, the same analysis shows an asthma-specific interaction of the TSLP-producing basal cells with NK cells, although TSLP-IL7Ra interactions with CD4 and CD8 T cells are also higher in asthma (Fig. 8I, Extended Fig. 10I). NK cells contribute to resolution of inflammation in asthma through induction of eosinophil apoptosis^74^, which was impaired in patients with severe asthma^75^. It remains unknown whether TSLP modulates this NK cell function in asthma. TSLP is well known to induce Th2 cell activity in CD4 T cells^45^.

### Non-canonical differentiation trajectory-associated cell states correlate with disease severity and FeNO

Finally, we evaluated whether the transitional and terminal cell states in the hillock-like epithelial differentiation trajectory were associated with specific clinical phenotypes in asthma (Fig. 9A). While nominal p-values should be interpreted with caution, a pattern arises in which the percentage of repair-associated basal resting cells and hypersecretory goblet cells per donor are positively correlated with age, disease severity and exhaled nitric oxide in patients with asthma, suggestive of a contribution towards a type-2 inflammation (Fig. 9A). This association is parallelled by the IL-13 driven upregulation of iNOS in hillock-like cells as they transition into goblet cells (Fig. 5C, Extended Fig. 8D). The proportion of hypersecretory goblet cells also increases with age in healthy control donors. Additionally, the proportion of *KRT14*+ basal resting and hillock-like cells per patient with asthma are correlated with smoking history (pack-years and past smoking, no current smokers were included in the cohort), in line with smoking-induced injury resulting in squamous metaplasia ^76^. Finally, we observe a positive association of hypersecretory goblet cell proportion with small airway obstruction (r5-r20), while it is negatively correlated with atopy, as measured by skin prick tests. This may point to the existence of atopy-independent disease mechanisms.

**Figure 9.**
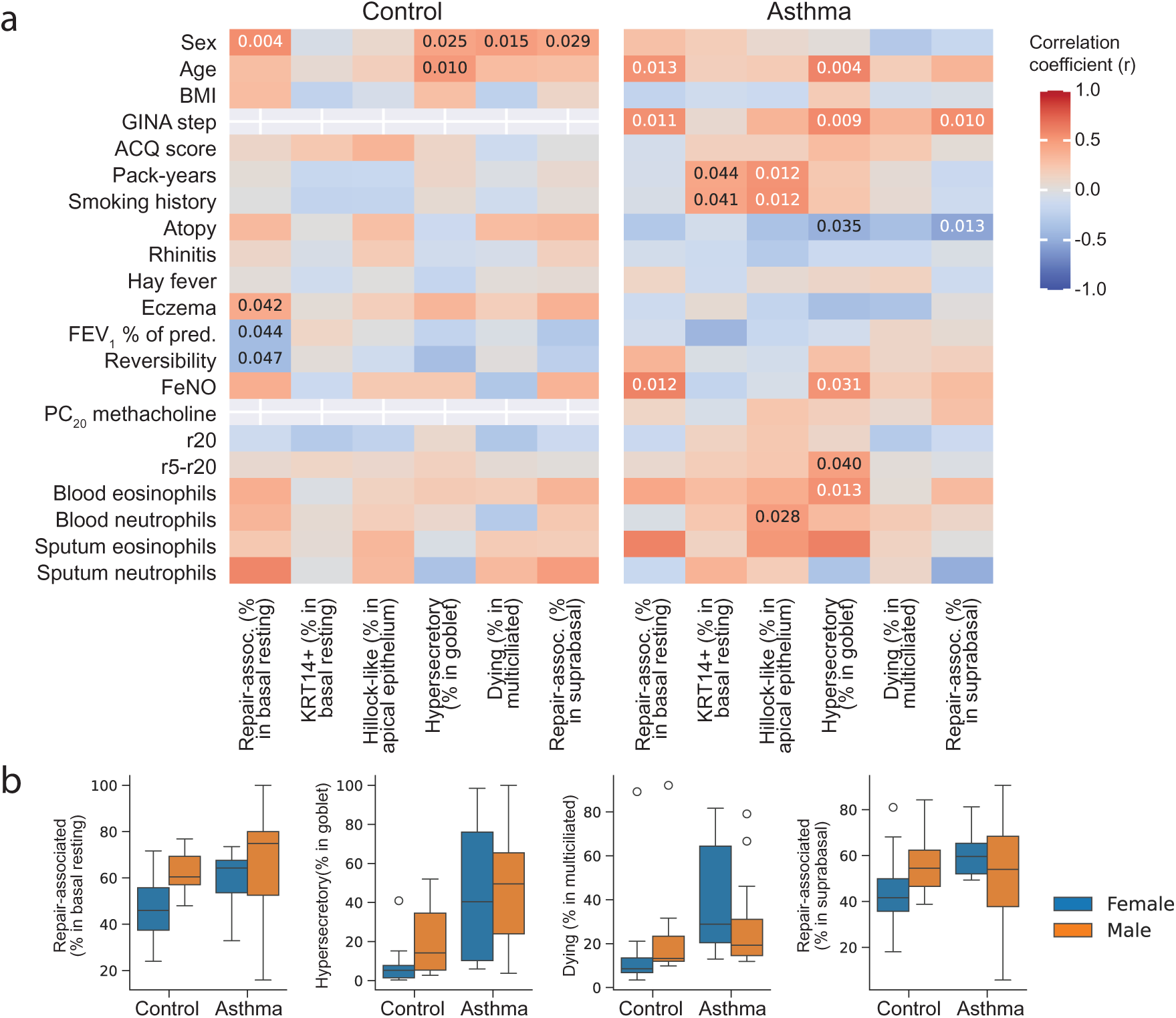
The non-canonical epithelial differentiation trajectory is associated with clinical and lung function parameters. **a)** Spearman correlation between selected clinical characteristics of the patients with asthma and healthy controls in the airway cell atlas and their epithelial cell type and cell state composition. Numbers indicate a nominally significant correlation (p-value ≤ 0.05). **b)** The proportion of repair-associated basal resting cells, hypersecretory goblet cells, and dying ciliated cells or repair-associated suprabasal cells as proportion of their parent cell type in male and female patients with asthma and healthy controls.

Interestingly, the percentage of repair-associated basal resting and suprabasal cells, *KRT14*+ basal resting cells, hypersecretory goblet cells, and dying multiciliated cells are all correlated with biological sex in healthy controls, but not in patients with asthma (Fig. 9A). In healthy controls, men have a higher proportion of these asthma-associated cell subsets than women, whereas this dimorphism is not observed in patients with asthma (Fig. 9A, 9B). Finally, we observe a positive association of repair-associated basal resting cell percentage in healthy controls and the prevalence of eczema, and negative associations with both the forced expiratory volume at one second (FEV_1_) and airflow reversibility. In conclusion, the proportion of the bronchial epithelial cell subsets in the non-canonical differentiation trajectory are associated with relevant clinical features of asthma including GINA score, parameters of small airway function and FeNO.

## Discussion

Here, we present a comprehensive single cell reference atlas of the lower airway wall in adults with childhood-onset asthma and matched healthy controls, identifying cellular mechanisms underpinning chronic airway inflammation and remodelling. We find that in patients with asthma, the airway epithelium has a phenotype of chronic repair, with increased differentiation through a novel non-canonical trajectory involving a *KRT14*+ proliferative basal cell state that transitions into a hillock-like epithelial cell state that is strongly increased in the pseudostratified bronchial epithelium of patients with asthma. We describe a fundamental heterogeneity of basal cells, with a repair-associated basal subset that forms the start of the non-canonical epithelial differentiation trajectory, and is increased in patients with asthma. We identify a strongly increased proportion of dying multiciliated cells in patients with asthma, alongside a striking increase in both epithelial differentiating activity and multiciliated cell fate decisions. Interestingly, transcriptomic features retained in dying multiciliated cells suggest a potential origin in the non-canonical epithelial differentiation pathway, though this needs further experimental validation. Our *in vitro* epithelial cultures show that the higher proportion of repair-associated basal cells is a cell-autonomous effect, and the increased proportion of hillock-like cells is not due to IL-13 signalling in patients with asthma. In contrast, IL-13 exposure of ALI cultured PBECs induces a transition from hillock-like cells to *CEACAM5*+ goblet cells, a subset also present in bronchial biopsies of patients with asthma, implicating the non-canonical differentiation trajectory in goblet cell metaplasia in asthma. All epithelial cell subsets in the non-canonical differentiation trajectory contribute to airway inflammation and remodelling, with high expression of matrix metalloproteases and metalloendopeptidases MMP10, MMP13 and ADAMTS1, as well as the alarmin TSLP and chemokines such as CCL2, CCL5, CXCL3 and CXCL8, recruiting monocytes, eosinophils, neutrophils and memory T-cells. This aligns with strongly increased proportions of proliferating T-cells, monocytes, pDCs, DC2s and mast cells that characterize the airways wall in patients with asthma, all of which contribute to reciprocal interactions between the airway epithelium and the immune cells, maintaining chronic inflammation of the airways. Association of the cell types in this novel epithelial differentiation trajectory with clinical parameters such as disease severity, FeNO levels and small airway function underscore the clinical relevance of these findings.

We observe an asthma-associated basal cell subset characterised by a fundamental change in overall transcriptional activity, including high expression of KLF4, SOX9 and AP-1 family members. Basal cell heterogeneity in the lower airways has been described before^39,62,77,78^, based on transcriptomic and functional features^78^ as well as on anatomical location within the airways^39,62^. Interestingly, basal cells from the dorsal bronchial wall are enriched for a subset of KRT17-expressing cells and give rise to a strongly increased proportion of KRT13+ cells in culture^39,62^. Also in the dorsal airway wall, patches of KRT13+ epithelial cells with squamous differentiation have been observed^62^. A dedicated epithelial subset named hillock cells was recently described in discrete anatomical structures of stratified epithelium with squamous morphology called hillocks^26^. In both the dorsal patches of so-called aberrant epithelium with high numbers of KRT13+ cells and in the hillock structures, an increased proportion of proliferating cells was observed^26,62^, which aligns with our observation of a *KRT14*+ proliferative transitional cell state in the differentiation towards hillock-like cells in the pseudostratified epithelium.

The increased proportion of repair-associated basal cells in bronchial epithelial cells from patients with asthma is retained in 3D spheroid epithelial cultures, indicating that this is a cell-autonomous feature independent of external factors. It remains unknown what triggers the outgrowth of this specific basal cell subset. In organoid cultures of human fetal lung buds (9 weeks post conception), hypoxia induced pronounced differentiation of hillock-like cells, which were mapped to one of three differentiation trajectories from basal cells, with two other branches also present under normoxia leading to secretory and neuroendocrine cells, respectively^79^. DamID-Seq and RNA-seq identified *KLF4* as an HIF2α target in hypoxic conditions^79^. We observe increased expression of *KLF4* throughout the non-canonical epithelial differentiation trajectory, and expression of *EPAS1* (HIF2α) and *KLF4* is higher in repair-associated basal cells and dying multiciliated cells, indicating that hypoxia could contribute to hillock-like differentiation in the pseudostratified epithelium of the lower airways in patients with asthma. However, the repair-associated basal cells also express *SOX9*, which may indicate an origin in the myoepithelial cells from the submucosal gland^80,81^. Although these cells from the SMG express *KRT14*, the *KRT14*+ cells in our dataset do not express SOX9, making it unlikely that these are myoepithelial cells. We observe increased numbers of KRT14+ surface epithelial cells in bronchial biopsies of patients with asthma, although these are also present in biopsies from healthy controls. Given the presence of a small, but highly proliferative, *KRT14*-expressing epithelial transitional cell state as part of the non-canonical epithelial differentiation trajectory, we propose that epithelial patches of KRT14+ cells reflect regions where epithelial repair through this pathway has recently occurred, for instance after viral infection, and where presence of the KRT14 protein remains after differentiation of these cells through the proliferative *KRT14+* basal cell state. In line with this, *KRT14* expression is limited in histologically normal regions of human tracheal epithelium, but was observed in hyperplastic, disrupted or metaplastic regions^82^.

Repair-associated basal cells express high levels of laminins that collectively form the laminin-322 protein in the basal lamina to which basal cells adhere through hemidesmosomes, which contain a6b4 integrins^83^. Expression of beta-6 integrin is only induced after epithelial damage ^84^ and contributes to IL-4 driven migration of airway epithelial cells^85^. Laminin-322 levels in the basement membrane are higher in patients with allergic asthma^86^, and laminin-322 supports motility of bronchial epithelial cells more than other ECM components^43^, with knockdown of *LAMC2* expression being sufficient to impair migration in several epithelial cancer models^40^.

What is the role of the non-canonical differentiation pathway in the known epithelial changes in asthma? We observe that IL-13 exposure induces differentiation of hillock-like cells into CEACAM5-positive goblet cells, a feature we find to be characteristic of goblet cells in patients with asthma. In a meta-analysis of asthma-specific gene expression in nasal and bronchial brushes from a total of 355 cases and 193 controls, *CEACAM5* was one of the genes with highest expression in patients with asthma compared to controls^87^, indicating that this goblet cell phenotype is commonly present in the asthmatic airways. Additionally, our findings of a hypersecretory subset of CEACAM5+ goblet cells, combined with dying multiciliated cells, provide insight into a mechanism that may contribute to mucus plugging in the airways of patients with asthma. Through the non-canonical differentiation pathway, and under the influence of IL-13, hillock-like cells in patients with asthma preferentially transition into this specific goblet cell population characterized by high levels of mucin production.

Our findings are relevant to the mechanisms underlying disease onset in at-risk individuals. Childhood-onset asthma has a strong genetic component, and recent analyses in a previously established tracheal scRNA-seq dataset^25^ as well as in nasal epithelial cells^88^ report strong expression of childhood-onset asthma genes in a *KRT6A*/*KRT13*/*SPRR3*-positive epithelial cell subset resembling hillock-like cells^88^ or in the entire basal/hillock-like/squamous trajectory^89^. *In vitro* analyses show that the increased proportion of repair-associated basal epithelial cells in patients with asthma is cell-autonomous, in line with a genetic predisposition driving this basal cell state. Onset of disease in childhood is often associated with severe lower respiratory tract infections by rhinovirus (RV) or respiratory syncytial virus (RSV)^90^, which induce extensive epithelial damage. We hypothesize that in children with susceptibility for asthma, the epithelial repair response is dominated by high activity of the repair-associated basal cells, proliferation of the KRT14+ transitional cells and high numbers of hillock-like cells. Through factors such as TSLP and MMPs, this may set off an inflammatory cascade with innate and adaptive immune cell activation culminating in chronic airway inflammation and remodelling. In line with this, expression of hillock-like cell genes was higher in bronchial biopsies from severely wheezing infants who developed asthma at school age compared to transient wheezers, in absence of any signs of eosinophilic infiltration or type-2 inflammation^89^. This indicates that the altered epithelial repair precedes, and possibly contributes to, type-2 inflammation in childhood-onset asthma.

Our study underscores the relevance of the alarmins IL-33 and TSLP to this process. We find that the IL-33 pathway is regulated by asthma-associated SNPs in a cell type specific fashion. Moreover, TSLP expression is a core feature of the repair-associated basal epithelial cell subset, which is strongly increased in patients with asthma. In mouse models, both Tslp and Il-33 have been shown to drive differentiation and activation of pathogenic Th2 cells^91,92, 93^, which are increased in the bronchial airway wall in patients with asthma, as we have previously shown^13^. Both TSLP and IL-33 have been shown to contribute to a checkpoint for the tissue-specific induction of type-2 immune responses by Th2 cells and ILC2s^94^. Interestingly, RV infection was sufficient to break inhalation tolerance and induce type-2 immunity in the lungs through TSLP and IL-33 release ^95^. Taken together, these findings support a model where the asthma-associated epithelial repair response through the non-canonical epithelial differentiation trajectory drives activation of the type-2 immune response, which leads to airway inflammation and remodelling, and chronic disease.

In summary, we present the asthma cell atlas: a novel, comprehensive and freely accessible resource for the study of the cellular and molecular changes in the lower airways in patients with asthma. We use this resource to establish a novel, non-canonical bronchial epithelial differentiation pathway associated with asthma, which contributes to airway inflammation by immune cell recruitment and activation, and to airway wall remodelling through release of matrix remodelling factors and by driving goblet cell hyperplasia. All cell states of this non-canonical pathway are increased in asthma, and their proportions are associated with clinical parameters such as disease severity and small airway function, underscoring the clinical relevance of our findings.

## Code availability

All code is available at: https://github.com/Nawijn-Group-Bioinformatics/Asthma_Cell_Atlas/.

## Data availability

Data is available upon reasonable request to.

## Methods

### Patient recruitment and ethical approval

All tissue donors were volunteers specifically recruited for participation in the ARMS study (NCT03141814)^1^. This study was approved by the Institutional Review Board (Medical Ethics Committee; METc) of the University Medical Center Groningen (ABR nr. NL 53173.042.15), and all subjects gave their written informed consent prior to participation. All subjects had a smoking history of ≤10 pack-years and were included in the age range of 18-65 years. Further inclusion criteria for control subjects were: no history of asthma or other respiratory diseases and lack of asthma-related medication; negative provocation test (i.e. PC_20_ methacholine >8 mg/mL, and adenosine 5′-monophosphate >320 mg/mL with a 2 minute protocol); lack of pulmonary obstruction (i.e. forced expiratory volume in 1 s (FEV_1_)/forced vital capacity (FVC) ≥70%), absence of lung function impairment (i.e. >90% FEV_1_% predicted) and AHR (PC20 >16mg/mL). Further inclusion criteria for patients with asthma: asthma diagnosis confirmed by objective measurements, i.e. either an increase in FEV_1_ of ≥12% from the initial value and ≥200 mL after inhaling 400 μg of salbutamol, or airway hyperresponsiveness (AHR) to methacholine (provocative concentration causing a 20% drop in FEV_1_ (PC_20_) ≤8 mg/mL); age of onset of asthma symptoms of ≤20 years; a documented asthma diagnosis during childhood; current use of inhaled corticosteroids with or without β2-agonists due to respiratory symptoms; and a positive provocation test (PC_20_ methacholine of ≤8mg/mL with a 2 minute protocol). Additionally, patients with asthma had clinically stable disease without asthma exacerbation or respiratory infection symptoms for at least six weeks prior to inclusion in the study. See Table S1A for subject characteristics.

Six weeks before the start of the study, patients with asthma stopped use of inhaled corticosteroids. All subjects were clinically characterized with pulmonary function and provocation tests and underwent a fiberoptic bronchoscopy under conscious sedation according to a standardized protocol. The bronchoscopy was postponed for ≥6 weeks for subjects who developed upper respiratory symptoms. For this study, macroscopically adequate endobronchial brushes and biopsies were collected between the third and sixth generation of the right lower and middle lobe. Extracted biopsies and brushes were placed in HBSS (Lonza, BE10543F) supplemented with 10% Penicillin (10000 U/mL)/Streptomycin (10000 μg) (Gibco) and kept on ice before being processed within 1 hour.

See supplementary tables S1A, S1B and S1C, S1D, S1E for clinical characteristics of the donors included in the airway cell atlas, 3D organoid or air-liquid-interface ALI culture models and the sorted CD4+ T-cell samples.

**Table S1A:** Clinical characteristics of the patients with asthma and healthy control donors included in the airway cell atlas of childhood-onset asthma. Sex, GINA step, smoking history and atopy are presented as: number (percentage) and tested with a Fisher’s exact test. BMI, FEV_1_ % of predicted, FEV_1_/FVC and neutrophil counts are presented as mean (SD) and tested with a t-test. All other variables are presented as median [IQR] and tested with a Mann-Whitney U test. Atopy is defined as a positive skin prick test.

|  |  | Control (n=25) | Asthma (n=21) | p-value |
| --- | --- | --- | --- | --- |
| Age |  | 55 [50, 59] | 52.00 [48, 61] | 0.74 |
| Sex | *Male* | 11 (44.0) | 13 (61.9) | 0.25 |
|  | *Female* | 14 (56.0) | 8 (38.1) |  |
| BMI |  | 25.4 (3.7) | 27.9 (4.7) | 0.05 |
| GINA step | *1* | - | 5 (25.0) | 1.00 |
|  | *2* | - | 5 (25.0) |  |
|  | *3* | - | 4 (20.0) |  |
|  | *4* | - | 6 (30.0) |  |
| ACQ score |  | 0.0 [0.0, 0.0] | 0.8 [0.3, 1.4] | <0.01 |
| Smoking history | *Ex-smoker* | 8 (32.0) | 3 (14.3) | 0.19 |
|  | *Never smoker* | 17 (68.0) | 18 (85.7) |  |
| Pack-years |  | 2.0 [1.0, 4.3] | 2.0 [1.5, 6.0] | 0.75 |
| Atopy | *Yes* | 9 (36.0) | 17 (81.0) | <0.01 |
|  | *No* | 16 (64.0) | 4 (19.0) |  |
| FEV<sub>1</sub> % of predicted |  | 113.23 (11.41) | 83.08 (13.49) | <0.01 |
| FEV<sub>1</sub>/FVC |  | 0.76 (0.06) | 0.64 (0.08) | <0.01 |
| Reversibility |  | 1.66 [0.22, 2.92] | 6.81 [5.76, 12.38] | <0.01 |
| FeNO |  | 16.60 [10.50, 20.70] | 35.55 [19.87, 51.50] | <0.01 |
| PC<sub>20</sub> methacholine mg/mL |  | - | 0.53 [0.20, 1.06] | - |
| r5 |  | 0.32 [0.27, 0.35] | 0.47 [0.33, 0.52] | <0.01 |
| r20-r5 |  | 0.02 [-0.01, 0.04] | 0.11 [0.03, 0.17] | <0.01 |
| Blood eosinophils × 10<sup>9</sup>/L |  | 0.12 [0.08, 0.17] | 0.20 [0.14, 0.29] | <0.01 |
| Blood neutrophils × 10<sup>9</sup>/L |  | 3.04 (1.20) | 3.45 (0.71) | 0.19 |
| Sputum eosinophils × 10<sup>9</sup>/L |  | 0.35 [0.05, 0.92] | 1.70 [1.00, 8.00] | 0.01 |
| Sputum neutrophils × 10<sup>9</sup>/L |  | 48.16 (21.99) | 58.69 (13.61) | 0.23 |

**Table S1B:** Clinical characteristics of the patients with asthma and healthy control donors included in the 3D epithelial organoid cultures dataset. Sex, GINA step, smoking history and atopy are presented as: number (percentage) and tested with a Fisher’s exact test. BMI, FEV_1_ % of predicted, FEV_1_/FVC and neutrophil counts are presented as mean (SD) and tested with a t-test. All other variables are presented as median [IQR] and tested with a Mann-Whitney U test. Atopy is defined as a positive skin prick test.

|  |  | Control (n=5) | Asthma (n=6) | p-value |
| --- | --- | --- | --- | --- |
| Age |  | 59 [58, 59] | 51.5 [48.25, 55.5] | 0.10 |
| Sex | *Male* | 4 (80.0) | 3 (50.0) | 0.55 |
|  | *Female* | 1 (20.0) | 3 (50.0) |  |
| BMI |  | 26.0 (3.2) | 31.1 (8.4) | 0.23 |
| GINA step | 1 | - | 1 (16.7) | 1.00 |
|  | 2 | - | 1 (16.7) |  |
|  | 3 | - | 1 (16.7) |  |
|  | 4 | - | 3 (50.0) |  |
| ACQ score |  | 0.0 [0.0, 0.0] | 1.4 [1.2, 1.8] | 0.01 |
| Smoking history | *Ex-smoker* | 3 (60.0) | 0 (0.0) | 0.06 |
|  | *Never smoker* | 2 (40.0) | 6 (100.0) |  |
| Pack-years |  | 2.0 [1.5, 3.0] | - | - |
| Atopy | *Yes* | 3 (60.0) | 5 (83.3) | 0.55 |
|  | *No* | 2 (40.0) | 1 (16.7) |  |
| FEV<sub>1</sub> % of predicted |  | 116.06 (11.36) | 80.45 (22.47) | 0.01 |
| FEV<sub>1</sub>/FVC |  | 0.75 (0.06) | 0.65 (0.11) | 0.09 |
| Reversibility |  | 2.10 [1.07, 5.40] | 6.04 [4.49, 15.02] | 0.36 |
| FeNO |  | 13.72 [12.26, 15.19] | 18.50 [15.46, 20.05] | 0.25 |
| PC<sub>20</sub> methacholine mg/mL |  | - | 0.32 [0.19, 1.16] | - |
| r5 |  | 0.28 [0.26, 0.33] | 0.42 [0.35, 0.52] | 0.07 |
| r20-r5 |  | 0.02 [0.02, 0.04] | 0.08 [0.03, 0.16] | 0.27 |
| Blood eosinophils × 10<sup>9</sup>/L |  | 0.11 [0.10, 0.17] | 0.20 [0.15, 0.39] | 0.17 |
| Blood neutrophils × 10<sup>9</sup>/L |  | 2.76 (0.55) | 3.73 (1.27) | 0.15 |
| Sputum eosinophils × 10<sup>9</sup>/L |  | 1.00 [1.00, 1.00] | 3.65 [0.98, 11.12] | 0.35 |
| Sputum neutrophils × 10<sup>9</sup>/L |  | 73.00 (2.83) | 58.50 (16.07) | 0.30 |

**Table S1C:** Clinical characteristics of the patients with asthma and healthy control donors included in the airway-liquid-interface cultures dataset. Sex, GINA step, smoking history and atopy are presented as: number (percentage) and tested with a Fisher’s exact test. BMI, FEV_1_ % of predicted, FEV_1_/FVC and neutrophil counts are presented as mean (SD) and tested with a t-test. All other variables are presented as median [IQR] and tested with a Mann-Whitney U test. Atopy is defined as a positive skin prick test.

|  |  | Control (n=8) | Asthma (n=7) | p-value |
| --- | --- | --- | --- | --- |
| Age |  | 58.0 [52, 59.5] | 51 [45, 63.5] | 0.60 |
| Sex | *Male* | 4 (50.0) | 4 (57.1) | 1.00 |
|  | *Female* | 4 (50.0) | 3 (42.9) |  |
| BMI |  | 24.9 (3.1) | 28.5 (5.4) | 0.13 |
| GINA step | 2 | - | 1 (14.3) | 1.00 |
|  | 3 | - | 1 (14.3) |  |
|  | 4 | - | 5 (71.4) |  |
| ACQ score |  | - | 1.3 [1.2, 1.5] | <0.01 |
| Smoking history | *Ex-smoker* | 1 (12.5) | 0 (0.0) | 1.00 |
|  | *Never smoker* | 7 (87.5) | 7 (100.0) |  |
| Pack-years |  | 2.0 [2.0, 2.0] | - | - |
| Atopy | *Yes* | 4 (50.0) | 3 (42.9) | 1.00 |
|  | *No* | 4 (50.0) | 4 (57.1) |  |
| FEV<sub>1</sub> % of predicted |  | 111.44 (15.01) | 81.58 (13.67) | <0.01 |
| FEV<sub>1</sub>/FVC |  | 0.76 (0.05) | 0.67 (0.09) | 0.04 |
| Reversibility |  | 2.11 [0.16, 2.74] | 10.81 [6.17, 17.04] | <0.01 |
| FeNO |  | 16.30 [14.62, 21.98] | 26.20 [22.35, 30.05] | 0.36 |
| PC<sub>20</sub> methacholine mg/mL |  | - | 0.76 [0.30, 3.01] | - |
| r5 |  | 0.30 [0.26, 0.36] | 0.42 [0.35, 0.50] | 0.05 |
| r20-r5 |  | 0.01 [-0.01, 0.03] | 0.08 [0.03, 0.16] | <0.01 |
| Blood eosinophils × 10<sup>9</sup>/L |  | 0.11 [0.06, 0.18] | 0.27 [0.15, 0.43] | 0.09 |
| Blood neutrophils × 10<sup>9</sup>/L |  | 3.17 (1.01) | 3.82 (0.97) | 0.25 |
| Sputum eosinophils × 10<sup>9</sup>/L |  | 0.10 [0.05, 0.15] | 1.00 [0.50, 13.75] | 0.37 |
| Sputum neutrophils × 10<sup>9</sup>/L |  | 60.15 (24.25) | 58.77 (10.95) | 0.93 |

**Table S1D:** Clinical characteristics of the patients with asthma and healthy control donors included in the CD4+ T-cell dataset. Sex, GINA step, smoking history and atopy are presented as: number (percentage) and tested with a Fisher’s exact test. BMI, FEV_1_ % of predicted, FEV_1_/FVC and neutrophil counts are presented as mean (SD) and tested with a t-test. All other variables are presented as median [IQR] and tested with a Mann-Whitney U test. Atopy is defined as a positive skin prick test.

|  |  | Control (n=13) | Asthma (n=13) | p-value |
| --- | --- | --- | --- | --- |
| Age |  | 57 [54, 59] | 53 [49, 60] | 0.38 |
| Sex | *Male* | 8 (61.5) | 9 (69.2) | 1.00 |
|  | *Female* | 5 (38.5) | 4 (30.8) |  |
| BMI |  | 24.7 (3.8) | 27.3 (6.5) | 0.22 |
| GINA step | 2 | - | 2 (15.4) | 1.00 |
|  | 3 | - | 7 (53.8) |  |
|  | 4 | - | 4 (30.8) |  |
| ACQ score |  | 0.0 [0.0, 0.0] | 1.20 [0.4, 1.4] | <0.01 |
| Smoking history | *Ex-smoker* | 6 (46.2) | 1 (7.7) | 0.07 |
|  | *Never smoker* | 7 (53.8) | 12 (92.3) |  |
| Pack-years |  | 3.5 [1.5, 4.0] | 2.0 [2.0, 2.0] | 0.61 |
| Atopy | *Yes* | 6 (46.2) | 11 (84.6) | 0.10 |
|  | *No* | 7 (53.8) | 2 (15.4) |  |
| FEV<sub>1</sub> % of predicted |  | 113.37 (13.50) | 84.67 (16.97) | <0.01 |
| FEV<sub>1</sub>/FVC |  | 0.73 (0.05) | 0.62 (0.10) | <0.01 |
| Reversibility |  | 4.56 [2.00, 5.65] | 6.53 [3.96, 11.12] | 0.16 |
| FeNO |  | 19.70 [12.90, 23.28] | 30.60 [19.50, 41.48] | 0.26 |
| PC<sub>20</sub> methacholine mg/mL |  | - | 0.88 [0.41, 2.46] | - |
| r5 |  | 0.34 [0.24, 0.40] | 0.38 [0.33, 0.58] | 0.08 |
| r20-r5 |  | 0.02 [-0.02, 0.04] | 0.08 [0.04, 0.19] | 0.02 |
| Blood eosinophils × 10<sup>9</sup>/L |  | 0.11 [0.08, 0.14] | 0.22 [0.18, 0.43] | <0.01 |
| Blood neutrophils × 10<sup>9</sup>/L |  | 2.81 (1.02) | 3.74 (1.15) | 0.04 |
| Sputum eosinophils × 10<sup>9</sup>/L |  | 1.00 [0.40, 1.00] | 4.50 [0.98, 9.07] | 0.12 |
| Sputum neutrophils × 10<sup>9</sup>/L |  | 55.09 (17.61) | 58.76 (26.29) | 0.76 |

**Table S1E:** Clinical characteristics of additional donors included in the asthma cell atlas: patients in remission of asthma, and the patients with asthma treated with ICS. Sex, GINA step, smoking history and atopy are presented as: number (percentage) and tested with a Fisher’s exact test. BMI, FEV_1_ % of predicted, FEV_1_/FVC and neutrophil counts are presented as mean (SD) and tested with a t-test. All other variables are presented as median [IQR] and tested with a Mann-Whitney U test. Atopy is defined as a positive skin prick test.

|  |  | Remission (n=11) | Asthma: ICS (n=7) |
| --- | --- | --- | --- |
| Age |  | 47 [44, 55] | 52 [40.5, 61.5] |
| Sex | *Male* | 6 (54.5) | 4 (57.1) |
|  | *Female* | 5 (45.5) | 3 (42.9) |
| BMI |  | 22.5 (1.9) | 27.2 (3.3) |
| GINA step | 3 | - | 3 (42.9) |
|  | 4 | - | 4 (57.1) |
| ACQ score |  | 0.0 [0.0, 0.0] | 1.3 [1.1, 1.5] |
| Smoking history | *Ex-smoker* | 1 (9.1) | 3 (42.9) |
|  | *Never smoker* | 10 (90.9) | 4 (57.1) |
| Pack-years |  | 5.0 [5.0, 5.0] | 4.0 [2.2, 4.5] |
| Atopy | *Yes* | 8 (72.7) | 4 (57.1) |
|  | *No* | 3 (27.3) | 3 (42.9) |
| FEV<sub>1</sub> % of predicted |  | 91.83 (21.67) | 105.00 (9.87) |
| FEV<sub>1</sub>/FVC |  | 0.69 (0.11) | 0.76 (0.08) |
| Reversibility |  | 5.43 [3.64, 8.06] | 1.46 [-1.06, 3.24] |
| FeNO |  | 15.68 [14.20, 18.10] | 25.94 [15.86, 37.92] |
| PC<sub>20</sub> methacholine mg/mL |  | 0.47 [0.26, 1.56] | 0.77 [0.70, 0.84] |
| r5 |  | 0.33 [0.30, 0.37] | 0.34 [0.28, 0.37] |
| r20-r5 |  | 0.02 [0.00, 0.04] | 0.03 [0.02, 0.06] |
| Blood eosinophils × 10<sup>9</sup>/L |  | 0.17 [0.14, 0.24] | 0.15 [0.10, 0.16] |
| Blood neutrophils × 10<sup>9</sup>/L |  | 2.73 (0.94) | 3.06 (0.84) |
| Sputum eosinophils × 10<sup>9</sup>/L |  | 0.00 [0.00, 0.30] | 1.45 [0.22, 2.70] |
| Sputum neutrophils × 10<sup>9</sup>/L |  | 28.70 (19.85) | 56.22 (24.84) |

### Cell culture

#### 3D culture of primary bronchial epithelial cells

For 6 patients with asthma and 5 healthy controls, primary bronchial epithelial cells (PBECs) obtained from the lung brushes and expanded into submerged cultures were thawed in T25 flasks as previously described^2^. Cells were cultured with Airway Epithelial Cell Growth medium (AEGM; Promocell C-21060), and medium was refreshed twice a week. After expanding them for one passage, cells were seeded in a Matrigel (Corning 354230). After solidification of the matrigel, it was covered by HUB medium^3^. Medium was refreshed twice a week. After two weeks, cells were passaged and seeded again in Matrigel. Briefly, the cells were first washed with ice cold HBSS (Lonza, BE10543F). After centrifugation for 5 min at 300 g, cells were resuspended TrypLE (Gibco 1260413) and incubated for 15 min at 37 °C. Next, trypsin-EDTA (0,25%) (Gibco 25200-056) was added and cells were incubated in this 1:1 TrypLE:trypsin-EDTA mixture for another 15 min at 37 °C. After centrifugation for 5 min at 300 g, cells were counted and seeded in Matrigel as described above.

#### Air-liquid-interface culture and IL-13 perturbation of primary bronchial epithelial cells

For 7 patients with asthma and 8 healthy controls, PBECs obtained from lung brushes were isolated by centrifugation and expanded in adherent, submerged growth conditions in Purecol/fibronectin/BSA coated flasks in AEGM with supplement mix (Promocell, C-21060). After two passages, cells were frozen down in liquid nitrogen. For experiments, cells were thawed and expanded submerged in AEGM medium for an additional two passages.

PBECs were seeded in Purecol/fibronectin/BSA pre-coated polyester transwell 0.4 µm pore inserts (Corning, 3470) at a density of 270 k/cm^2^. Cells were cultured under submerged conditions with 500 µl in the basolateral and 200 µl in the apical side of AEGM. After three days, apical medium was removed and basolateral medium was replaced with 1:1 ALI medium (AEGM:Dulbecco’s Modified Eagle Medium (DMEM, Lonza) 1:1 freshly supplemented with 15 ng/mL of retinoic acid Sigma, R2625). ALI medium was refreshed three times per week. After 4 weeks of air-liquid interface (ALI) culture, cells were apically and basolaterally stimulated for 5 days with 10 ng/mL of interleukin 13 (Immunotools, 11340137) or saline in ALI medium. 100 µl apical stimulation was added for 4 hours in ALI medium and then removed. After 5 days, the medium was refreshed with ALI medium and cells were cultured for another 20 days.

### Trans epithelial electrical resistance (TEER)

After airlift at the beginning of the ALI culture period, TEER measurements were taken weekly to assess the epithelial barrier integrity using a Millicel ERS-2 Volt ohmmeter (Merck, MERS00002). Briefly, 100 µl of AEGM without supplement mix was added apically and the electrode was inserted in the fluid. Electrical resistance was measured across the cell layer growing on the membrane of the transwell insert (in Ohm) and corrected for the electrical resistance of an empty insert.

### Tissue processing for single cell sequencing

#### Bronchial biopsies

A single cell suspension was obtained from the bronchial biopsies as previously described^4^. Briefly, after finely chopping the biopsies using a single edge razor blade, the tissue was dissociated for one hour into a mixture of 1 mg/mL collagenase D and 0.1 mg/mL DNase I (Roche) in HBSS (Lonza, BE10543F) at 37 °C in a water bath with shaking. Filtration of the suspension was performed through a 70 μm nylon cell strainer (Falcon) and cleared of red blood cells using a red blood cell lysis buffer (eBioscience). After counting with hemotocytometer, the single cell suspension was immediately loaded onto the 10X Genomics Chromium system, according to the manufacturer’s instructions (10X Genomics).

#### CD4+ T-cells for SmartSeq2 analysis

Bronchial biopsy and whole blood samples from the same patients were processed as previously described^4^. Briefly:

### CD4 T-cells from peripheral blood

Lithium heparin-anticoagulated whole blood (500 μl) was lysed using an ammonium chloride-potassium solution (155 mM ammonium chloride (NH4Cl), 10 mM potassium bicarbonate (KHCO3), 0,1 mM EDTA). Cells were centrifuged for 5 min at 4 °C, 550 g, after which the cell pellet was washed twice with PBS containing 1% BSA, followed by staining for cell surface markers. Blood leukocytes were stained with CD4 APC-Cy7, CD3 PerCP Cy5.5, CD8 APC, and CD45RA-PE (eBioscience) for 30 min at 4°C and washed twice with PBS containing 1% BSA. Propidium iodide was added 5 min before sorting.

### CD4 T-cells from bronchial biopsies

Airway wall biopsy single-cell suspensions (see above) were stained for 30 min at 4 °C with CD3 PerCP Cy5.5, CD45 BB515, CD4 APC-Cy7 (BD), and CD8 PE and washed twice with PBS containing 1% BSA. Propidium iodide was added 5 min before sorting.

### FACS sorting of individual CD4 T-cells for SmartSeq2

Lymphocytes were selected in the FCS/SSC plot. These were then selected on single, live cells for blood or single, live, CD45+ for lung. The sorted cells were positive for CD3 and CD4. All cells were sorted in individual wells of a 96-well plate using a MoFlo Astrios (Beckman Coulter) using Summit Software (Beckman Coulter).

#### 3D organoid cultures

On days 3, 7, 10, 14 and 17 after seeding, the 3D spheroid cultures were harvested following the same protocol described above, and processed for single cell-RNA sequencing (scRNA-seq) using the 10X Genomics Chromium system, according to the manufacturer’s instructions (10X Genomics).

#### Airway liquid interface cultures

ALI cultures were harvested for scRNA-seq either 3 or 17 days after the last IL-13 stimulation. Cells were removed from the insert using 0.25% trypsin-EDTA (Gibco 25200056) for 10 minutes at 37 °C and washed with HBSS (Lonza, BE10543F). PBECs were labelled with donor-specific hashtags using TotalSeq anti-B2M antibodies for demultiplexing during data processing. Next, cells from different donors were pooled and prepared for scRNA-seq library prep using Chromium 5’ V1.1 chemistry according to the instructions of the manufacturer (10X Genomics). RNA sequencing was performed using a Novaseq S6 with 26 cycles for read 1 and 91 cycles for read 2. Paired-end sequencing was performed aiming for 30.000 reads/cell for gene expression libraries and for 10.000 reads/cell for hashtag libraries.

### Single cell sequencing of 10XChromium libraries

Libraries for bronchial biopsies were prepared according to standard protocol of the Chromium single-cell 3′ kit v2, v3, or v3.1 (10X Genomics, CG000204). For the PBECs cultures (3D and ALIs), the single-cell Chromium Next GEM Single Cell V(D)J Reagent Kits v1.1 with Feature Barcode technology for Cell Surface Protein was used (10X Genomics, CG000208). Quality and concentrations of the libraries were assessed on TapeStation (Agilent). Paired-end 150 bp sequencing was performed with Illumina sequencing aiming for 30000 reads per cell.

### Library preparation and sequencing for SmartSeq2

Library preparation was performed with minor modifications from the published SmartSeq2 protocol^5^. In short, single cells were flow sorted onto individual wells of 96 or 384 wells containing 4 μl (96 wells) or 1 μl (384 wells) of lysis buffer (0.3% triton plus DNTPs and OligoDT). After sorting, plates were frozen and stored at −80 °C until further processing. PCR with reverse transcription (25 cycles) and Nextera library preparation performed as described in the published protocol^5^. Paired-end reads from SmartSeq2 were mapped to the human genome (GRCh38) using GSNAP with default parameters^6^. Then, uniquely mapped reads were counted using htseqcount (http://www-huber.embl.de/users/anders/HTSeq/).

### SCRINSHOT

Experimental procedure in SCRINSHOT was performed as previously described^8,9^. Briefly, tissue sections from fresh-frozen biopsies of healthy and diseased patients (generation 4-5 airway) were fixed for 10 minutes in 4% paraformaldehyde, treated with 1M HCl, blocked and incubated with padlock probe mix. DNA probes to detect expressed genes were used at 1-2 padlocks per gene, the rest was applied at 3-4 padlocks per gene. Ligation with SplintR, and probe rolling circle amplification using Phi29-polymerase were followed by fixation of and detection cycles. Detection probes in the form of fluorophore-labeled 15-25 nucleotide oligos were applied at 30 °C in 30% formamide solution, followed by washes at 30 °C in 20% formamide solution. All samples were processed with an epithelial cell type marker panel in a single experiment. The panel was composed of the following marker genes: *SCGB3A1, SCGB1A1, CAPS, MUC5AC, KRT5, LCN2, KRT13, KRT6A, KRT14, SPRR3, KRT15, IL33, KRT4, ITGA6, LTF*. It was applied to sections of biopsies from three asthmatic patients, two healthy controls and one healthy lung. Images were taken at 20x magnification using as a Z-stack with 11 steps of 0.8 μm (to cover the whole 10 μm thickness) at a widefield microscope (Zeiss Axio Observer Z.2, Carl Zeiss Microscopy GmbH, with a Colibri led light source, equipped with a Zeiss AxioCam 506 Mono digital camera and an automated stage).

### Histological imaging after SCRINSHOT

Biopsy sections were washed in PBS, and standard hematoxylin and eosin staining procedure was applied. Samples were washed, mounted in glycerol and imaged under light microscope. Images then were aligned with SCRINSHOT images from consecutive sections.

### Immunohistochemistry

PAS staining was performed using the Sakura Tissue-Tek Prisma Plus Automated Slide Stainer. Slides were deparaffinised and rinsed in demiwater. Stained for 10 min in 1% periodic acid, washed in demiwater and stained 15 min in Schiffs reagent. After another rinse step the slides were stained with Haematoxilin for 5 min and rinsed in water. After dehydration the slides were mounted and covered with a cover glass.

IHC for KRT14 was performed on three µm thick formalin-fixed paraffin tissue slides. Slides were deparaffinised and rinsed in demiwater. Antigen retrieval was performed by incubating the slides for 15 min in Tris-HCl buffer pH 9 in microwave at 300 W. After endogenous peroxidase blocking, the slides were incubated with a primary antibody (mouse anti-human Keratin14, Sigma C8791) 1:500 diluted in 1% BSA/PBS. After washing with PBS the slides were incubated with a secondary Multimer HRP reagent. DAB (3,3’-diaminobenzidine) was used to visualise the positive cells. After Haematoxilin staining, the slides were dehydrated, mounted, and covered with a cover glass.

### Data pre-processing

#### In vivo cell atlas, 3D organoid and ALI culture datasets

FastQ files were aligned to the GRCh38 reference using CellRanger (10X Genomics, v6.1.1). Per sample, FastCAR (v0.2) was used to correct for ambient RNA with the recommend.empty.cutoff function, generating the ambient profile according to default parameters^10^. For the ALI culture samples, the lower number of samples allowed for manual determination of the optimal cut-off per sample. Libraries with 500 UMIs were removed. Doublet detection was scored using Scrublet (v0.2.3) with default parameters. The organoid data was demultiplexed using SoupOrCell, whereas the air-liquid-interface data was demultiplexed using a combination of SoupOrCell and hashtags to allow for maximum recovery of cells. Initial quality control (QC) was performed by filtering out libraries with >25% mitochondrial reads and/or <200 expressed genes. The data was normalized using a sequencing depth of 10000 and log transformed, followed by highly variable gene (HVG) selection (4000 genes). Per sample, preliminary cell type annotation was performed using leiden clustering as implemented in Scanpy: clusters with clear marker gene expression were annotated, while the remaining clusters were labeled as unknown.

#### CD4+ T-cell dataset

Pre-processing of the CD4+ T-cell dataset was performed as previously described^4^, followed by the same initial per-sample QC filters for >25% mitochondrial reads and/or <200 expressed genes, log-normalisation and HVG selection, and a preliminary cell type annotation was performed on the combined data.

#### SCRINSHOT

Analysis of SCRISNHOT data was performed as described previously^9^. Projection and stitching, followed by image export was performed in Zen (2.3 lite). Tiling of SCRINSHOT images was performed in Fiji (ImageJ 1.53c), SCRINSHOT signal (dot) detection was done in CellProfiler (3.1.9). Automated nuclei segmentation was performed in BIAS on tiled images (via Image Filters function) using deep neural network model (DiscovAIR Segmentation, v.1.0) in Segmentation function with scaling 1.50-2.00, detection confidence 1%, contour confidence 50%. Nuclei regions were then expanded by 2 µm (without overlaps) to recapitulate cell ROIs. Assignment of dots to cells was performed in Fiji, as described previously^8^.

### Dataset integration, QC and cell type annotation

#### In vivo cell atlas

The single cell integration benchmarking package (scIB, v1.1.3) was used to determine the choice of integration method, testing scGen (v2.1.0), scVI (v0.18.0), scANVI (v0.18.0), Harmony (v0.1.7) and Scanorama (v1.7.3). scANVI was chosen to integrate the samples, specifying the sample ID as batch. After integration and leiden clustering, 147,918 cells belonging to low quality clusters were removed. The remainder was again integrated using scANVI. Next, QC and annotation were performed in an iterative manner on multiple levels of increasing cell type label resolution. Briefly, per level the neighbours graph and leiden clustering were calculated and clusters were assessed on quality metrics (number of counts and genes, mitochondrial count percentage), for doublet indications doublets (conflicting cell type marker gene expression, aided by scrublet doublet scores), and for sample-drivenness, followed by a new neighbours graph and clusters. Once all clusters passed this QC, these clusters were annotated based on established marker genes from the Human Lung Cell Atlas and other literature, and where needed by comparison to available reference datasets. Then, per cell type label a subset was created in which the same QC steps and the next level of annotation were performed, to a maximum of four levels. All subsets were merged and the remaining 327,077 cells that passed QC were integrated using scanVI on the final cell type labels.

#### Other datasets

The 3D organoid culture, ALI culture, and CD4+ T-cell datasets were integrated using scANVI. The culture datasets were integrated on sample ID, the CD4+ T-cell dataset on donor ID. QC and cell type annotation were performed in the same iterative fashion as described for the *in vivo* data, but was limited to two levels of annotation. Where necessary, additional rounds of integration were performed to improve the final annotation quality. The organoid and ALI culture datasets were again integrated on the final cell type labels using scANVI to aid embedding-based analyses. Samples taken at day 3 of the 3D organoid culture time series experiment were found to contain substantial proportions of fully differentiated, dying cells, remaining from the previous culturing passage. Therefore, these samples were excluded from all analyses.

#### SCRINSHOT

Low positive cells with less than 4 reads per cell were excluded. Normalization by total count was applied. Leiden clustering with 10 nearest neighbors, resolution 0.5 and pc number 6 was used for the combined dataset. Clusters were manually annotated according to expected marker genes. Manual cell type assignment for hillock-like cells was performed by positivity (≥2 transcripts per cell) for any of the following: *KRT4*, *SPRR3*, *KRT6A*, *KRT13*, and negativity (<5 transcripts per cell) for all of the following: *SCGB1A1*, *SCGB3A1*, *LTF*, *CAPS*, *KRT15*, *KRT5*, *MUC5AC*, *LCN2*. Other markers were not considered. Resulting cells could be further split into subcategories: *KRT6A*-positive and *KRT4*-positive. Among those, a few cells expressed high *KRT13*, and were manually separated (≥3 transcripts per cell) Additional cell type was assigned by *KRT14* positivity (≥2 transcripts per cell) and negativity for (<8 transcripts per cell) for all of the following: *SCGB1A1*, *SCGB3A1*, *LTF*, *CAPS*, *KRT15*, *KRT5*, *MUC5AC*, *LCN2*. This cell type was annotated *’KRT14*+ basal’. Annotated cell types were mapped by ROI coordinates using TissuUmaps.

### Computational analyses

#### Differential abundance analysis

We tested for differences in cell type composition with a logistic mixed effects regression model, using R package lme4 (v1.1-33). Cell types with 200 cells were tested (100 cells for the ALI culture dataset) as follows: *cell_type ∼ 1 + disease_status + BMI + (1 | donorID)*, where cell type is encoded as 0 or 1, and BMI was included as a covariate for the cell atlas, 3D organoid culture and sorted CD4+ T-cell datasets, with donor ID as a random effect. Differential cell type abundance in the 3D organoid time series data was tested for each timepoint separately. To assess differential cell type abundance for asthma PBMCs compared to control in the ALI dataset, this model was run including batch as a random effect, including early and late timepoint samples. To assess the effect of IL-13 on asthma compared to control PBECs in the ALI culture dataset, the model was amended to include the treatment group and interaction effect, and run for early and late timepoints separately: *cell_type ∼ 1 + treatment + disease_status + treatment:disease_status + (1 | donorID) + (1 | batch)*.

#### Differential gene expression analysis

Differential gene expression (DGE) was tested on pseudobulks generated per cell type and sample using R package decoupler (v1.5.0), including cells with ≥500 UMIs and samples with ≥10 cells per cell type. To mitigate cross-lineage effects of ambient RNA, per level 1 annotation label we removed all genes with 90% of UMIs in the cell atlas expressed outside of the tested subset. Following TMM normalisation, DGE analysis was performed using quasi-likelihood testing of generalized linear models (*glmQLFit*, *glmQLTest* functions) as implemented in R package EdgeR (v3.40.0) for all cell types with ≥3 pseudobulk sampled per disease group. For the 3D organoid culture datasets, samples were included across timepoints. BMI was included as a covariate in analyses of the airway cell atlas, ALI culture dataset, 3D organoid culture and sorted CD4+ T-cell datasets. When comparing transcriptional phenotypes within a cell type, disease status was included as a covariate. P-values were FDR corrected.

Following DGE analysis, gene set enrichment analysis was performed using a hypergeometric test as implemented in gProfiler on gene sets in the GO:BP database. Where enrichment analysis resulted in many GO terms with overlapping definitions, we clustered the results on their differentially expressed genes, and selected the lowest p-value representative term per cluster for visualisation.

#### Gene module identification

For data-driven identification of gene expression modules, we used the Disentangled Representation Variational Inference^11^ (DRVI, v0.1.2) run on the HVG non-normalized counts of all cells in the airway cell atlas. Donor ID was included as a covariate. The number of latent factors was set at 64, with default encoder and decoder dimensions, training the model for 400 epochs. Contributing genes per latent factor were identified using DRVI’s *traverse_latent* and *calculate_differential_vars* functions, plotting the top 10 contributing genes per module for interpretation.

#### Identification of within-cell-type subsets

To assess transcriptional heterogeneity within the goblet cell and multiciliated lineage subsets, we clustered each subset of cells across a range of 15 resolution values between 0.2 and 3.0, increasing in increments of 0.2. Each subset was randomly subsampled to 90% of its cells ten times, to identify a meaningful clustering resolution by calculating the Adjusted Rand Index (ARI), implemented in the sklearn Python package (v1.6.1), across the ten subsets per resolution. The resolution with the highest ARI score was used to cluster and annotate the full subsets of goblet cells and multiciliated lineage cells, respectively.

Conversely, after a binomial distribution of DRVI-derived gene module activity was found in the basal cell subset, a high- and low-activity basal cell subset were annotated at a resolution most suitable to distinguish these groups. Similarly, after observing heterogeneous expression of basal, hillock-like, club and goblet cell type marker genes within the hillock-like cell subset, hillock-like cell subsets were annotated at a resolution that best separated these distinct expression profiles.

#### Pseudotime and cell fate mapping

Pseudotime across the epithelial differentiation trajectory was calculated on the basal, hillock-like, secretory and multiciliated lineage subsets in the cell atlas, excluding cycling basal cells using Palantir (v1.3.3). This was done using 20 diffusion components calculated on the integrated embedding in asthma and control respectively, as a combined analysis did not yield a reliable model, selecting a basal resting cell and goblet and multiciliated cells as the starting and terminal points, respectively. Cell type density across pseudotime was visualised using a kernel density estimate plot (Seaborn, v0.13.2). Diffusion maps for both the epithelial cell types in the cell atlas, as well as the whole 3D organoid culture dataset were generated by calculating diffusion pseudotime for 15 components using the Scanpy (v 1.10.4) implementation. A PAGA graph was generated using the PAGA implementation in Scanpy, on the basal resting, suprabasal, hillock-like and club cells in the cell atlas.

Cell fate probabilities per cell type were estimated with CellRank (v2.0.6) using the pseudotime kernel on the Palantir pseudotime for the cell atlas data, and diffusion pseudotime for the 3D organoid data. As initial and terminal states, we specified one detected macrostate of basal resting cells and two macrostates of goblet and multiciliated cells in the cell atlas. In the 3D organoid dataset, two groups of 30 outlier cells, driven by donor- or quality effects, were initially detected as dominant macrostates and therefore removed. After re-calculating diffusion pseudotime, cell fate probabilities were estimated using one initial macrostate composed of basal cells, and two macrostates that were detected in the hillock-like and secretory subsets each, which were combined per cell type to estimate an overall cell fate probability per label. Finally, the average terminal cell fate per cluster was calculated using CellRank’s *aggregate_fate_probabilities* function.

#### Transcription factor activity analysis

Gene regulatory networks were inferred from scRNA-seq data using the pySCENIC pipeline (v0.11.2) for the epithelial and immune cell subsets separately, applied to a Loom-formatted expression matrix of 4000 highly variable genes per lineage. TF–target interactions were identified using the grn module with a hg38 transcription factor list (from the Aerts Lab resources: https://resources.aertslab.org) and a tree-based method (GENIE3/GRNBoost). These interactions were refined by motif enrichment using cisTarget ranking databases and motif annotations from the same resource, with dropout masking enabled. Regulon activity was subsequently quantified per cell using AUCell and used for further downstream analysis.

#### Cell-cell interaction analysis

We examined differential cell-cell communication patterns in patients with asthma compared to healthy controls using MultiNicheNet^12^ (v2.1.0) including BMI as a covariate. Per cell type, we included samples with ≥10 cells per donor, excluding condition specific cell types, with recommended values for *min_sample_prop* and *fraction_cutoff*, and using empirical p-values. Ligand-target inference was performed using the top 250 predicted target genes per interaction, for upregulating ligands.

To interpret these results, we examined the top 50 prioritized cell-cell interactions per cell type as the receiver and as the sender, in both patients with asthma and healthy controls, as well as the overall top 50 interactions in asthma and control, respectively, and compared cell states by examining those cell-cell interactions with the highest differences in prioritization score within the disease groups. Individual results are shown as the MultiNicheNet prioritization scores in asthma and in control per ligand/receptor and sender/receiver combination, reflecting differential expression in patients with asthma compared to control, as well as cell type specificity, downstream signalling activity, and the fraction of samples with adequate expression. An additional MultiNicheNet analysis was performed on lower resolution cell type labels, used for greater clarity of plotting where appropriate.

#### eQTL analysis

Genotyping was performed in two batches on Illumina Infinium GSA-24 v3.0 arrays. Merged genotypes were filtered to exclude SNPs with >10% missingness, after which samples with >10% missingness were removed. Subsequently we filtered SNPs with a MAF <5% and samples with heterozygosity rate ±3 standard deviations. Finally, we filtered variants with the HWE p-value cut-off set to 1e-6.

From the cell atlas, the mean normalized expression per donor and cell type was determined using the function *pp.pseudobulk* from the package decoupler, including donors with ≥5 cells from all disease groups in the cell atlas, including patients in remission of asthma or using inhaled corticosteroids, to increase statistical power. Cell types with ≥40 pseudobulk samples were tested for genes expressed in at least a third of donors and with a mean pseudobulk expression of >0.1. Expression per gene was quartile normalized per cell type. Per subset we used TensorQTL (v1.0.10) with the --cis-nominal/--cis setting and the default 1MB window on both sides of the gene, including age, sex, disease group, BMI, and the first principal component of the variation of expression subset in that cell type as covariates. From this analysis we extracted the results of the genes *IL33* and *TSLP*.

#### Additional statistical methods

Gene module and transcription factor activity scores as well as differences in expression of targeted genes in the hillock-like subset were tested using a mixed effects model including donor ID as a random factor using the R package lme4 (v1.1-33), or Python package statsmodels (v0.14.4). TEER data was analysed using a mixed effects model with donor ID as a random effect, as implemented in Graphpad PRISM 9.

## References

1. Kato, A. & Kita, H. The immunology of asthma and chronic rhinosinusitis. Nat. Rev. Immunol. 25, 569–587 (2025).

2. Porsbjerg, C., Melén, E., Lehtimäki, L. & Shaw, D. Asthma. Lancet 401, 858–873 (2023).

3. Lambrecht, B. N., Ahmed, E. & Hammad, H. The immunology of asthma. Nat. Immunol. 26, 1233–1245 (2025).

4. Wenzel, S. E. Asthma phenotypes: the evolution from clinical to molecular approaches. Nat. Med. 18, 716–725 (2012).

5. Sayers, I., John, C., Chen, J. & Hall, I. P. Genetics of chronic respiratory disease. Nat. Rev. Genet. 25, 534–547 (2024).

6. El-Husseini, Z. W., Gosens, R., Dekker, F. & Koppelman, G. H. The genetics of asthma and the promise of genomics-guided drug target discovery. Lancet Respir Med 8, 1045–1056 (2020).

7. Alladina, J., et al. A human model of asthma exacerbation reveals transcriptional programs and cell circuits specific to allergic asthma. Sci. Immunol. 8, eabq6352 (2023).

8. Heijink, I. H. et al. Epithelial cell dysfunction, a major driver of asthma development. Allergy 75, 1902–1917 (2020).

9. Xiao, C. et al. Defective epithelial barrier function in asthma. J. Allergy Clin. Immunol. 128, 549–56.e1–12 (2011).

10. Hackett, T.-L. et al. Intrinsic phenotypic differences of asthmatic epithelium and its inflammatory responses to respiratory syncytial virus and air pollution. Am. J. Respir. Cell Mol. Biol. 45, 1090–1100 (2011).

11. Hewitt, R. J. & Lloyd, C. M. Regulation of immune responses by the airway epithelial cell landscape. Nat. Rev. Immunol. 21, 347–362 (2021).

12. Brusselle, G. G. & Koppelman, G. H. Biologic therapies for severe asthma. N. Engl. J. Med. 386, 157–171 (2022).

13. Vieira Braga, F. A., et al. A cellular census of human lungs identifies novel cell states in health and in asthma. Nat. Med. 25, 1153–1163 (2019).

14. Joulia, R. et al. A single-cell spatial chart of the airway wall reveals proinflammatory cellular ecosystems and their interactions in health and asthma. Nat. Immunol. 26, 920–933 (2025).

15. Wen, H. et al. Nasal airway transcriptome reflects selected asthma-associated gene signatures in the lower airways. Allergy (2026) doi:10.1111/all.70283.

16. Xu, C. et al. Probabilistic harmonization and annotation of single-cell transcriptomics data with deep generative models. Mol. Syst. Biol. 17, e9620 (2021).

17. Luecken, M. D. et al. Benchmarking atlas-level data integration in single-cell genomics. Nat. Methods 19, 41–50 (2022).

18. Heumos, L. et al. Best practices for single-cell analysis across modalities. Nat. Rev. Genet. 24, 550–572 (2023).

19. Luecken, M. D. & Theis, F. J. Current best practices in single-cell RNA-seq analysis: a tutorial. Mol. Syst. Biol. 15, e8746 (2019).

20. Sikkema, L. et al. An integrated cell atlas of the lung in health and disease. Nat. Med. 29, 1563–1577 (2023).

21. Saglani, S. & Lloyd, C. M. Novel concepts in airway inflammation and remodelling in asthma. Eur. Respir. J. 46, 1796–1804 (2015).

22. Montoro, D. T. et al. A revised airway epithelial hierarchy includes CFTR-expressing ionocytes. Nature 560, 319–324 (2018).

23. Shah, V. S. et al. Single cell profiling of human airway identifies tuft-ionocyte progenitor cells displaying cytokine-dependent differentiation bias in vitro. Nat. Commun. 16, 5180 (2025).

24. Deprez, M. et al. A single-cell atlas of the human healthy airways. Preprint at 10.1101/2019.12.21.884759.

25. Yoshida, M. et al. Local and systemic responses to SARS-CoV-2 infection in children and adults. Nature 602, 321–327 (2022).

26. Lin, B. et al. Airway hillocks are injury-resistant reservoirs of unique plastic stem cells. Nature 629, 869–877 (2024).

27. Koh, K. D. et al. Genomic characterization and therapeutic utilization of IL-13-responsive sequences in asthma. Cell Genom. 3, 100229 (2023).

28. Woodruff, P. G. et al. Genome-wide profiling identifies epithelial cell genes associated with asthma and with treatment response to corticosteroids. Proc. Natl. Acad. Sci. U. S. A. 104, 15858–15863 (2007).

29. Lackie, P. M., Baker, J. E., Günthert, U. & Holgate, S. T. Expression of CD44 isoforms is increased in the airway epithelium of asthmatic subjects. Am. J. Respir. Cell Mol. Biol. 16, 14–22 (1997).

30. Moinfar, A. A. & Theis, F. J. Disentangling cellular heterogeneity into interpretable biological factors through structured latent representations. bioRxiv (2024) doi:10.1101/2024.11.06.622266.

31. Rogers, D. F. Airway goblet cells: responsive and adaptable front-line defenders. Eur. Respir. J. 7, 1690–1706 (1994).

32. Blackburn, J. B., Li, N. F., Bartlett, N. W. & Richmond, B. W. An update in club cell biology and its potential relevance to chronic obstructive pulmonary disease. Am. J. Physiol. Lung Cell. Mol. Physiol. 324, L652–L665 (2023).

33. Setty, M. et al. Characterization of cell fate probabilities in single-cell data with Palantir. Nat. Biotechnol. 37, 451–460 (2019).

34. Browaeys, R., et al. MultiNicheNet: a flexible framework for differential cell-cell communication analysis from multi-sample multi-condition single-cell transcriptomics data. bioRxiv (2023) doi:10.1101/2023.06.13.544751.

35. Puddicombe, S. M. et al. Involvement of the epidermal growth factor receptor in epithelial repair in asthma. FASEB J. 14, 1362–1374 (2000).

36. Enomoto, Y. et al. Tissue remodeling induced by hypersecreted epidermal growth factor and amphiregulin in the airway after an acute asthma attack. J. Allergy Clin. Immunol. 124, 913–20.e1–7 (2009).

37. Polosa, R. et al. Expression of c-erbB receptors and ligands in the bronchial epithelium of asthmatic subjects. J. Allergy Clin. Immunol. 109, 75–81 (2002).

38. Inoue, H. et al. Dysfunctional ErbB2, an EGF receptor family member, hinders repair of airway epithelial cells from asthmatic patients. J. Allergy Clin. Immunol. 143, 2075– 2085.e10 (2019).

39. Zhou, Y. et al. Airway basal cells show regionally distinct potential to undergo metaplastic differentiation. Elife 11, e80083 (2022).

40. Rousselle, P. & Scoazec, J. Y. Laminin 332 in cancer: When the extracellular matrix turns signals from cell anchorage to cell movement. Semin. Cancer Biol. 62, 149–165 (2020).

41. Walko, G., Castañón, M. J. & Wiche, G. Molecular architecture and function of the hemidesmosome. Cell Tissue Res. 360, 529–544 (2015).

42. Lau, C. H. et al. Lentiviral expression of wild-type LAMA3A restores cell adhesion in airway basal cells from children with epidermolysis bullosa. Mol. Ther. 32, 1497–1509 (2024).

43. Kligys, K. et al. Laminin-332 and α3β1 integrin-supported migration of bronchial epithelial cells is modulated by fibronectin. Am. J. Respir. Cell Mol. Biol. 49, 731–740 (2013).

44. Moon, Y. W. et al. LAMC2 enhances the metastatic potential of lung adenocarcinoma. Cell Death Differ. 22, 1341–1352 (2015).

45. Stanbery, A. G., Shuchi Smita, Jakob von Moltke, Tait Wojno, E. D. & Ziegler, S. F. TSLP, IL-33, and IL-25: Not just for allergy and helminth infection. J. Allergy Clin. Immunol. 150, 1302–1313 (2022).

46. Kuo, C.-H. S. et al. Contribution of airway eosinophils in airway wall remodeling in asthma: Role of MMP-10 and MET. Allergy 74, 1102–1112 (2019).

47. Nkyimbeng, T. et al. Pivotal role of matrix metalloproteinase 13 in extracellular matrix turnover in idiopathic pulmonary fibrosis. PLoS One 8, e73279 (2013).

48. Christopoulou, M.-E., Papakonstantinou, E. & Stolz, D. Matrix metalloproteinases in chronic obstructive pulmonary disease. Int. J. Mol. Sci. 24, 3786 (2023).

49. Kumar, N., Mishra, B., Athar, M. & Mukhtar, S. Inference of gene regulatory network from single-cell transcriptomic data using pySCENIC. Methods Mol. Biol. 2328, 171–182 (2021).

50. Douma, S. et al. Suppression of anoikis and induction of metastasis by the neurotrophic receptor TrkB. Nature 430, 1034–1039 (2004).

51. Dragunas, G. et al. Cholinergic neuroplasticity in asthma driven by TrkB signaling. FASEB J. 34, 7703–7717 (2020).

52. Boaru, D. L. et al. The role of the LOX family in cancer. Eur. J. Cell Biol. 105, 151527 (2026).

53. Izzo, L. T. et al. KLF4 promotes a KRT13+ hillock-like state in squamous lung cancer. bioRxivorg (2025) doi:10.1101/2025.03.10.641898.

54. Sachs, N. et al. Long-term expanding human airway organoids for disease modeling. EMBO J. 38, (2019).

55. Weiler, P. & Theis, F. J. CellRank: consistent and data view agnostic fate mapping for single-cell genomics. Nat. Protoc. (2026) doi:10.1038/s41596-025-01314-w.

56. Sountoulidis, A. et al. SCRINSHOT enables spatial mapping of cell states in tissue sections with single-cell resolution. PLoS Biol. 18, e3000675 (2020).

57. Dupont, L. L., Glynos, C., Bracke, K. R., Brouckaert, P. & Brusselle, G. G. Role of the nitric oxide-soluble guanylyl cyclase pathway in obstructive airway diseases. Pulm. Pharmacol. Ther. 29, 1–6 (2014).

58. Feng, Y. et al. Cadherin-26 amplifies airway epithelial IL-4 receptor signaling in asthma. Am. J. Respir. Cell Mol. Biol. 67, 539–549 (2022).

59. Esnault, S. et al. Identification of bronchial epithelial genes associated with type 2 eosinophilic inflammation in asthma. J. Allergy Clin. Immunol. 155, 1510–1520 (2025).

60. Revinski, D. R. et al. CDC20B is required for deuterosome-mediated centriole production in multiciliated cells. Nat. Commun. 9, 4668 (2018).

61. Choksi, S. P. et al. An alternative cell cycle coordinates multiciliated cell differentiation. Nature 630, 214–221 (2024).

62. Zhang, Y. et al. Human airway basal cells undergo reversible squamous differentiation and reshape innate immunity. Am. J. Respir. Cell Mol. Biol. 68, 664–678 (2023).

63. Gomperts, B. N., Kim, L. J., Flaherty, S. A. & Hackett, B. P. IL-13 regulates cilia loss and foxj1 expression in human airway epithelium. Am. J. Respir. Cell Mol. Biol. 37, 339–346 (2007).

64. Gerovac, B. J. & Fregien, N. L. IL-13 Inhibits Multicilin Expression and Ciliogenesis via Janus Kinase/Signal Transducer and Activator of Transcription Independently of Notch Cleavage. Am. J. Respir. Cell Mol. Biol. 54, 554–561 (2016).

65. Dua, B., Watson, R. M., Gauvreau, G. M. & O’Byrne, P. M. Myeloid and plasmacytoid dendritic cells in induced sputum after allergen inhalation in subjects with asthma. J. Allergy Clin. Immunol. 126, 133–139 (2010).

66. Bratke, K. et al. Dendritic cell subsets in human bronchoalveolar lavage fluid after segmental allergen challenge. Thorax 62, 168–175 (2007).

67. Chairakaki, A.-D. et al. Plasmacytoid dendritic cells drive acute asthma exacerbations. J. Allergy Clin. Immunol. 142, 542–556.e12 (2018).

68. Greer, A. M. et al. Accumulation of BDCA1^+^ dendritic cells in interstitial fibrotic lung diseases and Th2-high asthma. PLoS One 9, e99084 (2014).

69. El-Gammal, A. et al. Allergen-induced changes in bone marrow and airway dendritic cells in subjects with asthma. Am. J. Respir. Crit. Care Med. 194, 169–177 (2016).

70. Vroman, H., Hendriks, R. W. & Kool, M. Dendritic cell subsets in asthma: Impaired tolerance or exaggerated inflammation? Front. Immunol. 8, 941 (2017).

71. Saikumar Jayalatha, A. K., Hesse, L., Ketelaar, M. E., Koppelman, G. H. & Nawijn, M. C. The central role of IL-33/IL-1RL1 pathway in asthma: From pathogenesis to intervention. Pharmacol. Ther. 225, 107847 (2021).

72. Herrera-De La Mata, S., et al. Cytotoxic CD4+ tissue-resident memory T cells are associated with asthma severity. Med (N. Y.) 4, 875–897.e8 (2023).

73. Wein, A. N. et al. CXCR6 regulates localization of tissue-resident memory CD8 T cells to the airways. J. Exp. Med. 216, 2748–2762 (2019).

74. Barnig, C. et al. Lipoxin A4 regulates natural killer cell and type 2 innate lymphoid cell activation in asthma. Sci. Transl. Med. 5, 174ra26 (2013).

75. Duvall, M. G. et al. Natural killer cell-mediated inflammation resolution is disabled in severe asthma. Sci Immunol 2, (2017).

76. Peters, E. J. et al. Squamous metaplasia of the bronchial mucosa and its relationship to smoking. Chest 103, 1429–1432 (1993).

77. Carraro, G. et al. Single-cell reconstruction of human basal cell diversity in normal and idiopathic pulmonary fibrosis lungs. Am. J. Respir. Crit. Care Med. 202, 1540–1550 (2020).

78. Reynolds, S. D. et al. Human bronchial basal cells are a community of variants. iScience 28, 113633 (2025).

79. Dong, Z. et al. Hypoxia promotes airway differentiation in the human lung epithelium. Cell Stem Cell 32, 1705–1722.e9 (2025).

80. Tata, A. et al. Myoepithelial cells of submucosal glands can function as reserve stem cells to regenerate airways after injury. Cell Stem Cell 22, 668–683.e6 (2018).

81. Lynch, T. J. et al. Submucosal gland myoepithelial cells are reserve stem cells that can regenerate mouse tracheal epithelium. Cell Stem Cell 22, 653–667.e5 (2018).

82. Ghosh, M. et al. Human tracheobronchial basal cells. Normal versus remodeling/repairing phenotypes in vivo and in vitro. Am. J. Respir. Cell Mol. Biol. 49, 1127–1134 (2013).

83. Hogan, B. L. M. et al. Repair and regeneration of the respiratory system: complexity, plasticity, and mechanisms of lung stem cell function. Cell Stem Cell 15, 123–138 (2014).

84. Huang, X. Z. et al. Inactivation of the integrin beta 6 subunit gene reveals a role of epithelial integrins in regulating inflammation in the lung and skin. J. Cell Biol. 133, 921–928 (1996).

85. Lee, S.-N. et al. Integrins αvβ5 and αvβ6 mediate IL-4-induced collective migration in human airway epithelial cells. Am. J. Respir. Cell Mol. Biol. 60, 420–433 (2019).

86. Amin, K. et al. Uncoordinated production of Laminin-5 chains in airways epithelium of allergic asthmatics. Respir. Res. 6, 110 (2005).

87. Tsai, Y.-H., Parker, J. S., Yang, I. V. & Kelada, S. N. P. Meta-analysis of airway epithelium gene expression in asthma. Eur. Respir. J. 51, 1701962 (2018).

88. Walsh, J. M. L., et al. Unique nasal cell states induced by common pediatric respiratory viruses. bioRxivorg (2026) doi:10.64898/2026.04.20.719671.

89. Kersten, E. T. G. et al. Childhood-onset asthma is characterized by airway epithelial hillock- to-squamous differentiation in early life. bioRxiv (2023) doi:10.1101/2023.07.31.549680.

90. Jackson, D. J. & Gern, J. E. Rhinovirus infections and their roles in asthma: Etiology and exacerbations. J. Allergy Clin. Immunol. Pract. 10, 673–681 (2022).

91. Khan, M. et al. Single-cell and chromatin accessibility profiling reveals regulatory programs of pathogenic Th2 cells in allergic asthma. Nat. Commun. 16, 2565 (2025).

92. Rochman, Y. et al. TSLP signaling in CD4+ T cells programs a pathogenic T helper 2 cell state. Sci. Signal. 11, eaam8858 (2018).

93. Meisel, C. et al. Regulation and function of T1/ST2 expression on CD4+ T cells: induction of type 2 cytokine production by T1/ST2 cross-linking. J. Immunol. 166, 3143–3150 (2001).

94. Van Dyken, S. J. et al. A tissue checkpoint regulates type 2 immunity. Nat. Immunol. 17, 1381–1387 (2016).

95. Mehta, A. K. et al. Rhinovirus infection interferes with induction of tolerance to aeroantigens through OX40 ligand, thymic stromal lymphopoietin, and IL-33. J. Allergy Clin. Immunol. 137, 278–288.e6 (2016).

## References for the Methods

1. Wen, H. et al. Nasal airway transcriptome reflects selected asthma-associated gene signatures in the lower airways. Allergy (2026) doi:10.1111/all.70283.

2. Gay, A. C. A. et al. Airway epithelial cell response to RSV is mostly impaired in goblet and multiciliated cells in asthma. Thorax 79, 811–821 (2024).

3. Sachs, N. et al. Long-term expanding human airway organoids for disease modeling. EMBO J. 38, e100300 (2019).

4. Vieira Braga, F. A. et al. A cellular census of human lungs identifies novel cell states in health and in asthma. Nat. Med. 25, 1153–1163 (2019).

5. Picelli, S. et al. Full-length RNA-seq from single cells using Smart-seq2. Nat. Protoc. 9, 171–181 (2014).

6. Wu, T. D. & Nacu, S. Fast and SNP-tolerant detection of complex variants and splicing in short reads. Bioinformatics 26, 873–881 (2010).

7. van den Brink, S. C. et al. Single-cell sequencing reveals dissociation-induced gene expression in tissue subpopulations. Nat. Methods 14, 935–936 (2017).

8. Sountoulidis, A. et al. SCRINSHOT enables spatial mapping of cell states in tissue sections with single-cell resolution. PLoS Biol. 18, e3000675 (2020).

9. Firsova, A. B. et al. Spatial single-cell atlas reveals regional variations in healthy and diseased human lung. Nat. Commun. 16, 9745 (2025).

10. Berg, M. et al. FastCAR: fast correction for ambient RNA to facilitate differential gene expression analysis in single-cell RNA-sequencing datasets. BMC Genomics 24, 722 (2023).

11. Moinfar, A. A. & Theis, F. J. Disentangling cellular heterogeneity into interpretable biological factors through structured latent representations. bioRxiv (2024) doi:10.1101/2024.11.06.622266.

12. Browaeys, R. et al. MultiNicheNet: a flexible framework for differential cell-cell communication analysis from multi-sample multi-condition single-cell transcriptomics data. bioRxiv (2023) doi:10.1101/2023.06.13.544751.

