## Extended Figures for "A repair-associated bronchial epithelial differentiation trajectory through *KRT14*+ basal and hillock-like cells drives airway inflammation and remodelling in childhood-onset asthma"

1a

| Method | Bio conservation: Isolated labels | Bio conservation: KMeans NMI | Bio conservation: KMeans ARI | Bio conservation: Silhouette label | Bio conservation: cLISI | Batch correction: Silhouette batch | Batch correction: iLISI | Batch correction: KBET | Batch correction: Graph connectivity | Batch correction: PCR comparison | Aggregate score: Batch correction | Aggregate score: Bio conservation | Aggregate score: Total |
|---|---|---|---|---|---|---|---|---|---|---|---|---|---|
| **X_scGen** | 0.87 | 1.00 | 1.00 | 0.99 | 1.00 | 0.96 | 0.58 | 1.00 | 1.00 | 1.00 | 0.91 | 0.97 | 0.95 |
| **X_scANVI** | 0.33 | 0.63 | 0.11 | 0.83 | 0.11 | 0.57 | 0.56 | 0.09 | 0.91 | 0.05 | 0.44 | 0.40 | 0.42 |
| **Unintegrated** | 0.74 | 0.56 | 0.21 | 0.76 | 0.39 | 0.00 | 0.00 | 0.19 | 0.89 | 0.00 | 0.22 | 0.53 | 0.41 |
| **X_Harmony** | 1.00 | 0.38 | 0.00 | 0.00 | 0.00 | 0.30 | 1.00 | 0.71 | 0.27 | 0.26 | 0.51 | 0.28 | 0.37 |
| **X-Scanorama** | 0.00 | 0.68 | 0.22 | 1.00 | 0.36 | 0.58 | 0.31 | 0.00 | 0.00 | 0.00 | 0.18 | 0.45 | 0.34 |
| **X_scVI** | 0.10 | 0.00 | 0.10 | 0.93 | 0.15 | 1.00 | 0.41 | 0.04 | 0.62 | 0.13 | 0.44 | 0.26 | 0.33 |

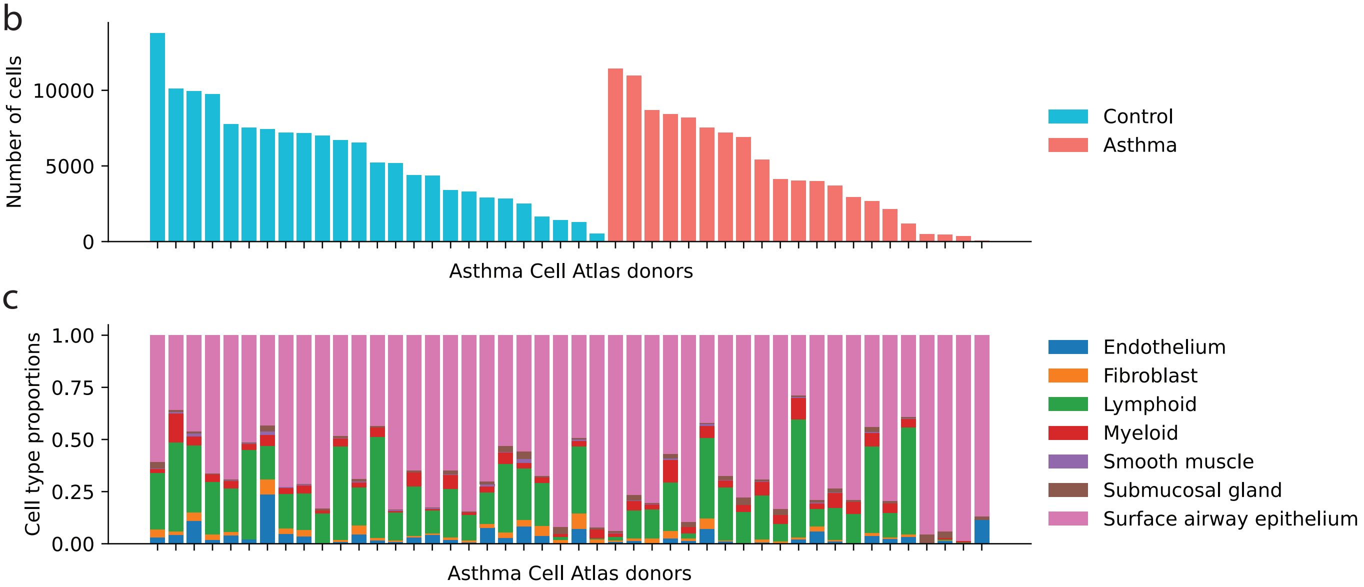


**Extended Figure 1. Integration benchmarking and composition of the airway cell atlas. a)** Overview of integration methods ranked by scIB total score for the asthma cell. Individual metrics for bio conservation and batch correction is indicated as min/max scaled score. **b)** Number of cells per donor in the asthma cell atlas after integration and QC, and **c)** the proportion of major cell type lineages.

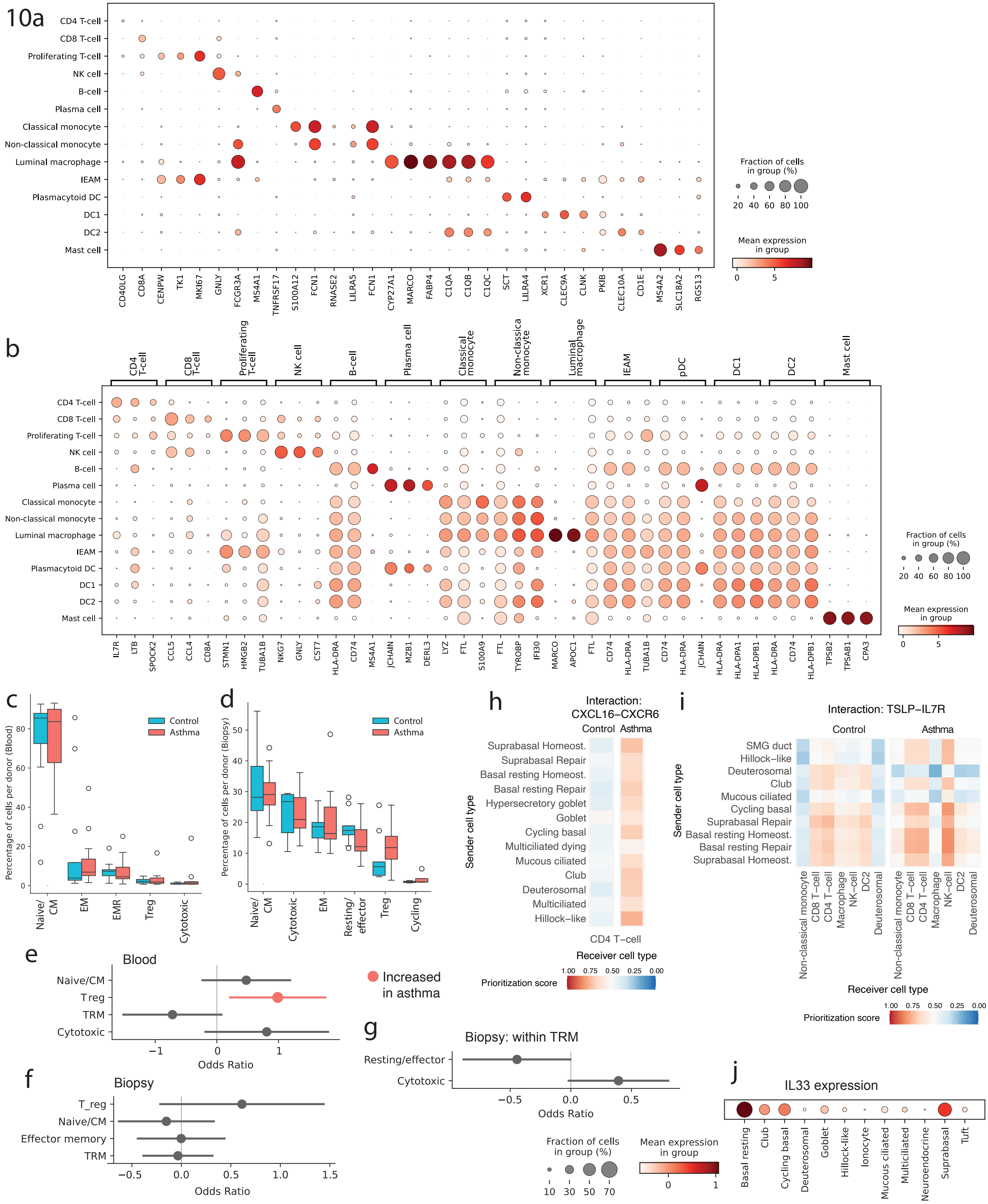


**Extended Figure 10. Immune cell gene expression and T-cell proportions in the asthma cell atlas. a)** Expression of classic cell type marker genes, and **b)** the top 3 cell-type-specific genes per immune cell type as annotated in the asthma cell atlas. **c)** Percentages of CD4+ T-cell subsets as part of total CD4+ T-cell numbers per donor, purified from peripheral blood and **d)** from bronchial biopsies. **e)** Odds ratios of the abundance of CD4+ T-cell subsets in patients with asthma compared to healthy control in peripheral blood and **f)** bronchial biopsies, and **g)** within the tissue resident subset of the bronchial biopsies (logistic mixed effects model at the single cell level). **h)** MultiNicheNet prioritization score for *CXCL16-CXCR6* cell-cell communication to CD4+ T-cells from epithelial cell types (on the y-axis) in asthma and control. **i)** MultiNicheNet prioritization score for *IL7R-TSLP* cell-cell communication from immune cells (x-axis) to epithelial cell types (y-axis) in healthy controls (left) and patients with asthma (right). **j)** IL-33 gene expression in surface epithelium cell types in the asthma cell atlas.

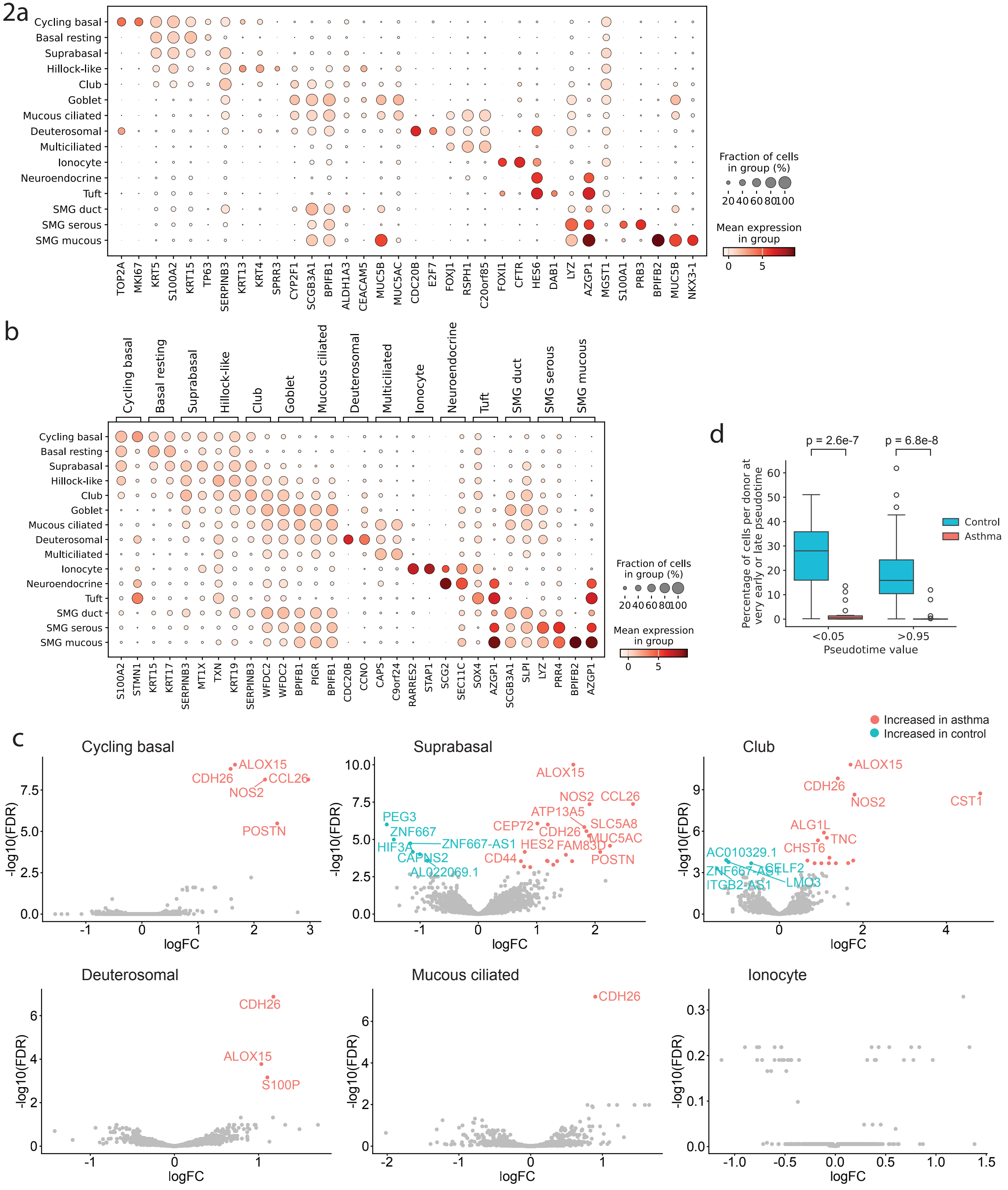


**Extended Figure 2. Cell type specific marker gene expression and asthma-associated differential gene expression. a)** Expression of classic cell type marker genes, and **b)** the top 2 cell-type-specific genes per epithelial cell type as annotated in the asthma cell atlas. **c)** Log10(fold-change) versus -log10(FDR-corrected p-value) of differential gene expression in selected epithelial cell types between patients with asthma and controls. **d)** Surface epithelium cell proportion per donor with a Palantir pseudotime value of <0.05 (left panels) >0.95 (right panels) in controls and patients with asthma, with median and interquartile range.

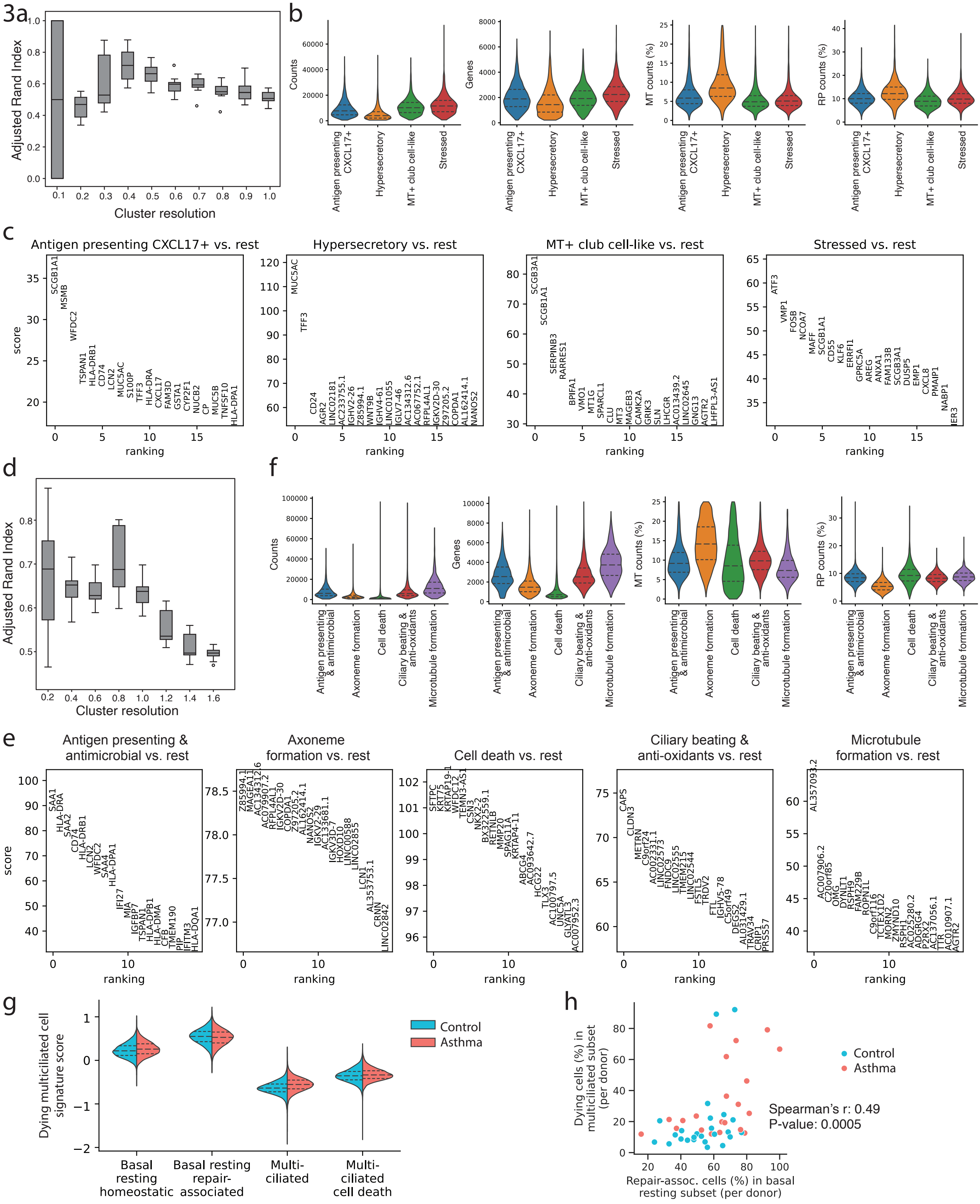

**Extended Figure 3. Subsets of goblet cells and multiciliated lineage cells in the asthma cell atlas. a)** Adjusted Rand Index for the different resolutions of clustering the goblet cell subset in the asthma cell atlas. **b)** QC criteria for the goblet cell subsets. (MT: mitochondrial genes; PR: ribosomal protein-encoding genes.) **c)** Ranked marker gene expression per goblet cell subset (T-test). **d)**, **e)**, and **f)** show the same for the multiciliated cell subset. **g)** Enrichment score for a dying multiciliated cell signature in basal resting cell subsets, multiciliated cells and dying multiciliated cells. **h)** Percentage of repair-associated cells in all basal resting cells versus percentage of dying cells in all multiciliated cells, per donor.

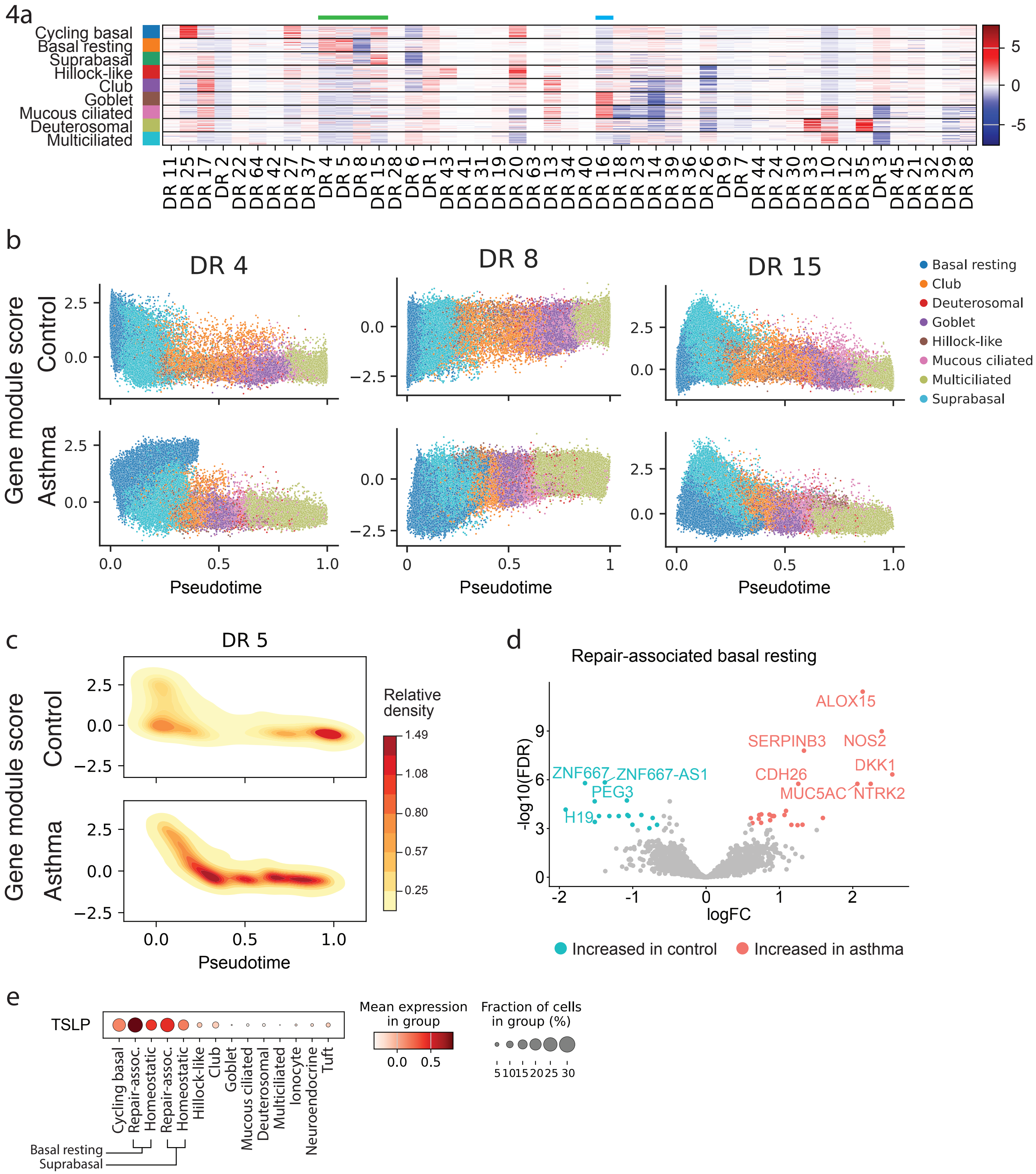


**Extended Figure 4. Data-driven identification of gene programs identifies a latent factor overexpressed in basal resting cells from patients with asthma. a)** Expression levels of latent factors identified using DRVI, per cell type. Green and blue lines indicate factors active in basal cells and a mucin production factor, respectively. **b)** Gene module score for the DR4, DR8 and DR15 latent factors across Palantir pseudotime in the surface epithelium. **c)** Density map of gene module score for latent factor DR5 across Palantir pseudotime. **d)** Log10(fold-change) versus -log10(FDR-corrected p-value) of differential gene expression in repair-associated basal resting cells between patients with asthma and controls. **e)** *TSLP* gene expression in the surface airway epithelial cell types of the asthma cell atlas.

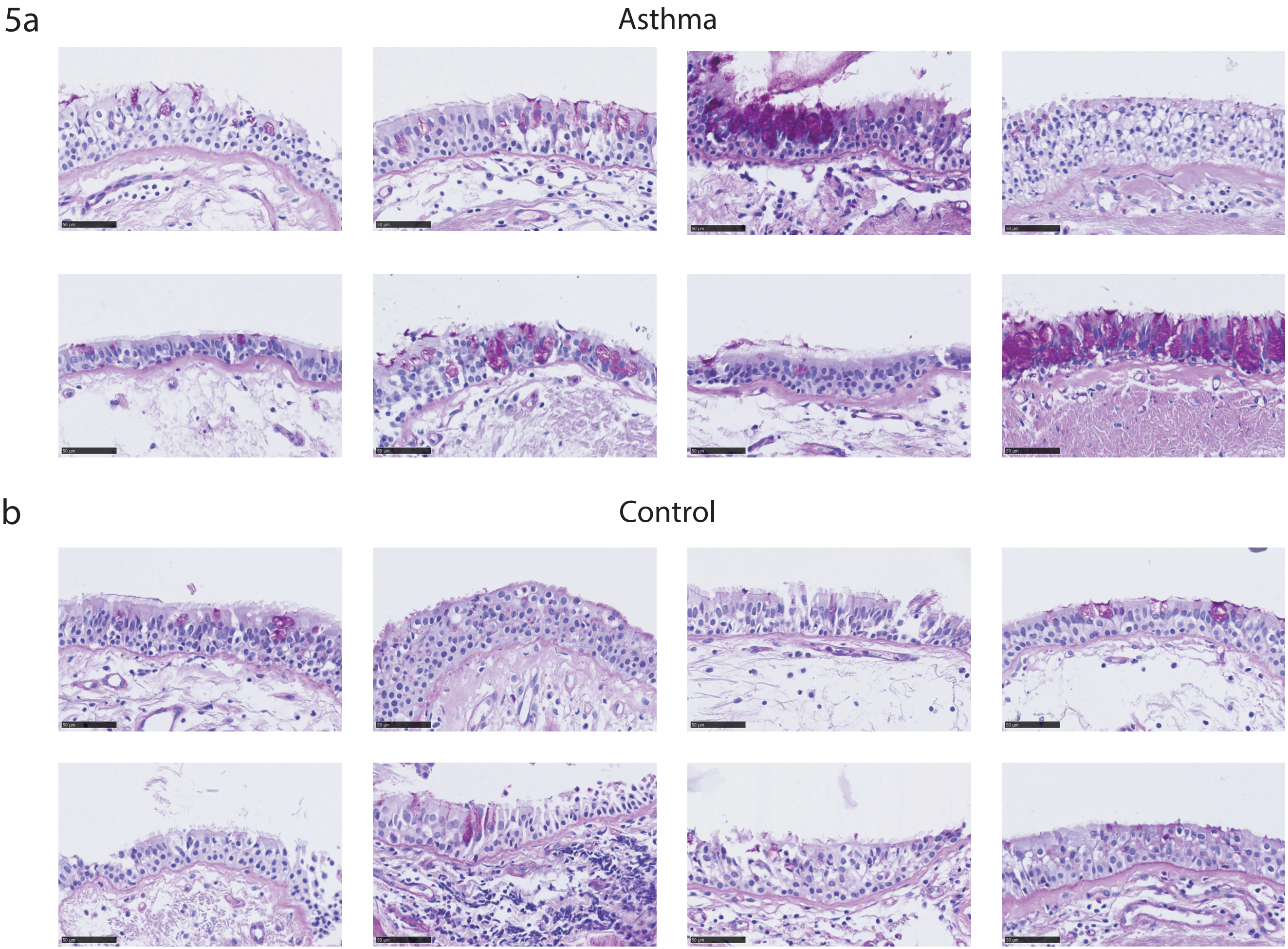


**Extended Figure 5. Goblet cell metaplasia in the airway epithelium of patients with asthma. a)** PAS stainings of the airway epithelium using 4 uM section of formalin-fixed paraffin embedded (FFPE) biopsies from patients with childhood-onset asthma, and **b)** healthy controls of the ARMS cohort. Representative images spanning at least 200 uM of intact airway epithelium are shown from 8 independent donors. Scale bar indicates 50 uM.

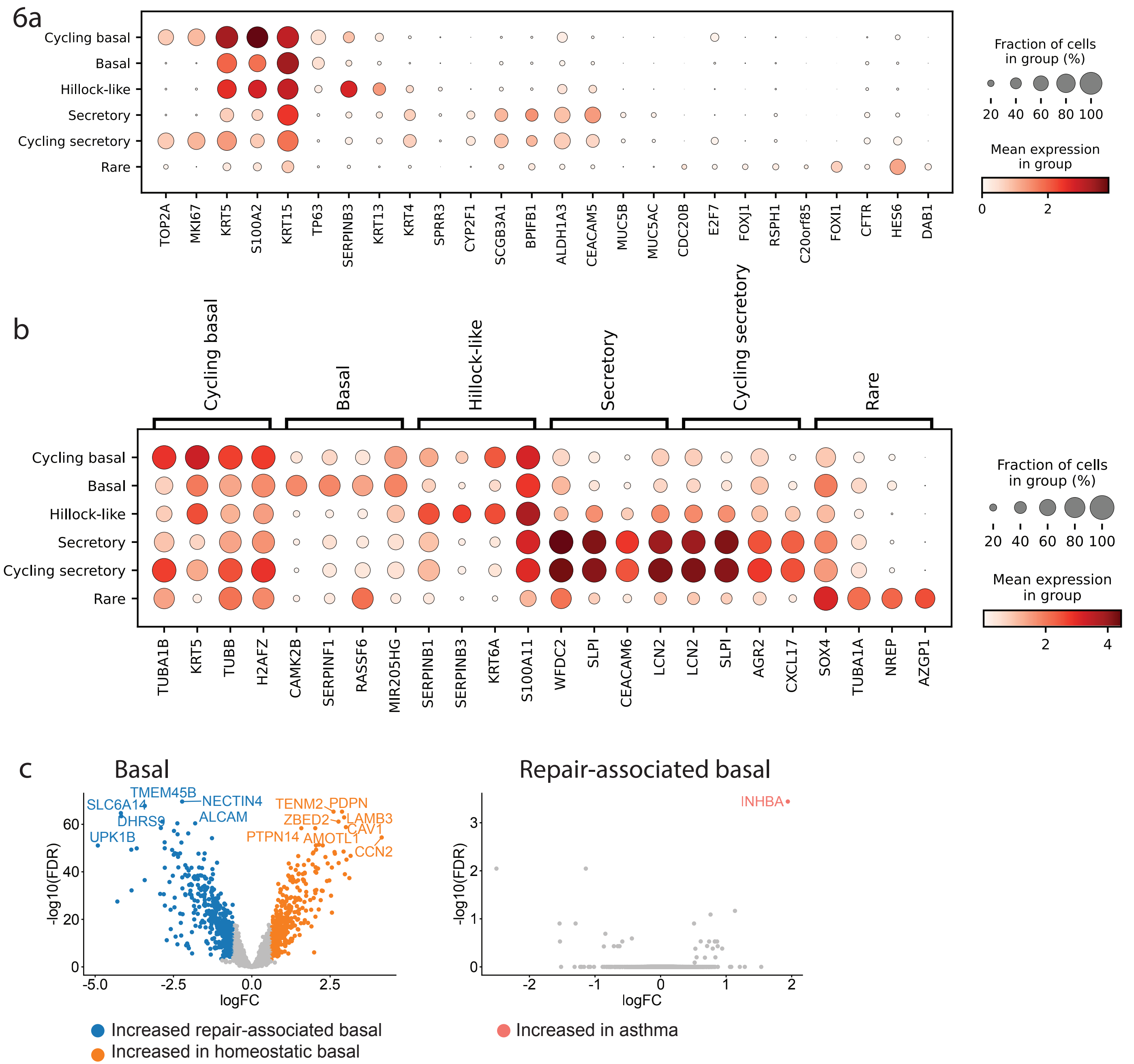


**Extended Figure 6. Gene expression in primary bronchial epithelial cells cultured in 3D epithelial spheroids.** **a)** Expression of classic cell type marker genes, and b) the top 4 cell-type-specific genes per cell type as annotated in 3D epithelial organoid dataset. **c)** Log10(fold-change) versus -log10(FDR-corrected p-value) of differential gene expression in 3D epithelial spheroid cultures between repair-associated basal cells and homeostatic basal cells, and **d)** in repair-associated basal cells between patients with asthma and controls.

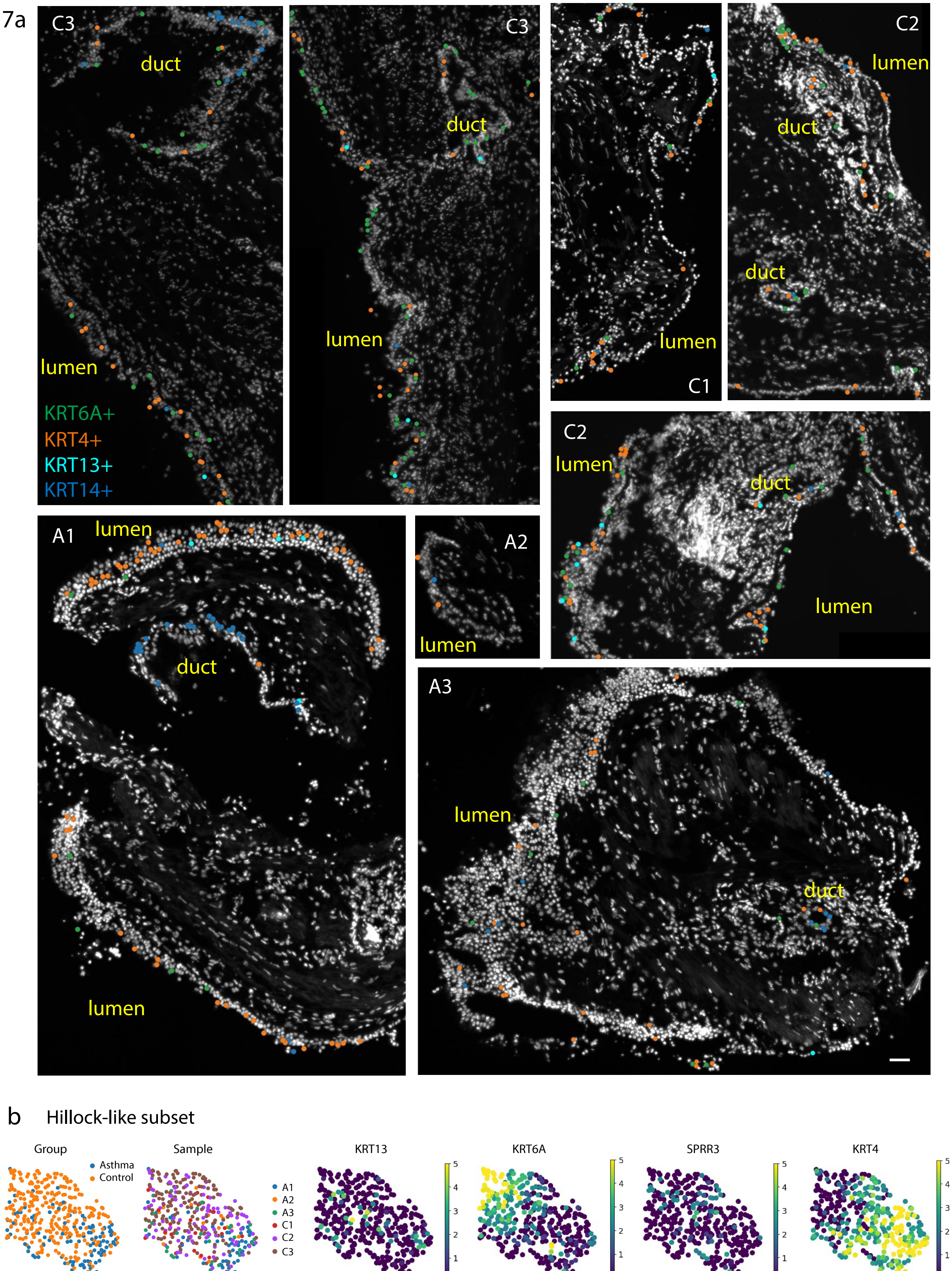


**Extended Figure 7. Distribution and characterization of hillock-like epithelial subpopulations across all analysed samples by spatial transcriptomics. a)** Spatial cell type maps from all analysed bronchial biopsies of patients with asthma (A1-3) and healthy controls (C1-3), showing the subtypes of hillock-like cells on nuclei (DAPI-stained, white) images. Scale bar: 50 $\mu$m. Lumen and submucosal gland ducts labeled in yellow. **b)** UMAPs of hillock-like cells indicating gene expression and sample distribution.

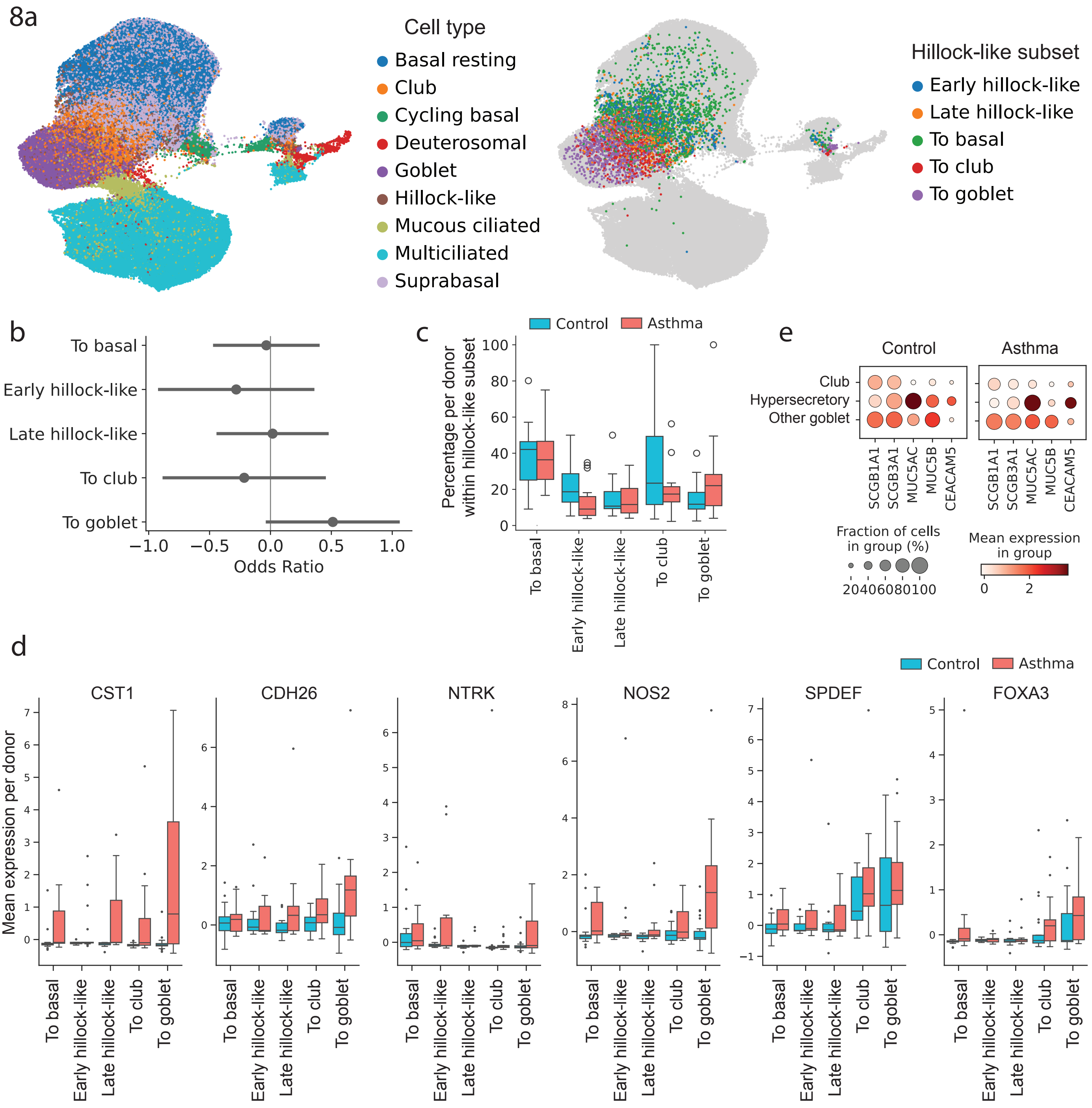


**Extended Figure 8. Transcription factor activity govern Hillock-like cell differentiation patterns. a)** UMAP of an embedding based on transcription factor activity (pySCENIC) of surface airway epithelial cells, showing cell type (left) and hillock-like subset (right) per cell. **b)** Odds ratios of the abundance of the subsets as part of the total hillock-like cells population in patients with asthma compared to healthy controls (logistic mixed effects model at the single cell level), and **c)** their percentages per donor. **d)** Mean expression of *CST1, CDH26, NTRK, NOS2, SPDEF* and *FOXA3* in hillock-like cell subsets per donor in patients with asthma and healthy controls. **e)** Expression of classic club and goblet cell marker genes in club cells, hypersecretory goblet cells and all other goblet cells subsets in patients with asthma and healthy controls.

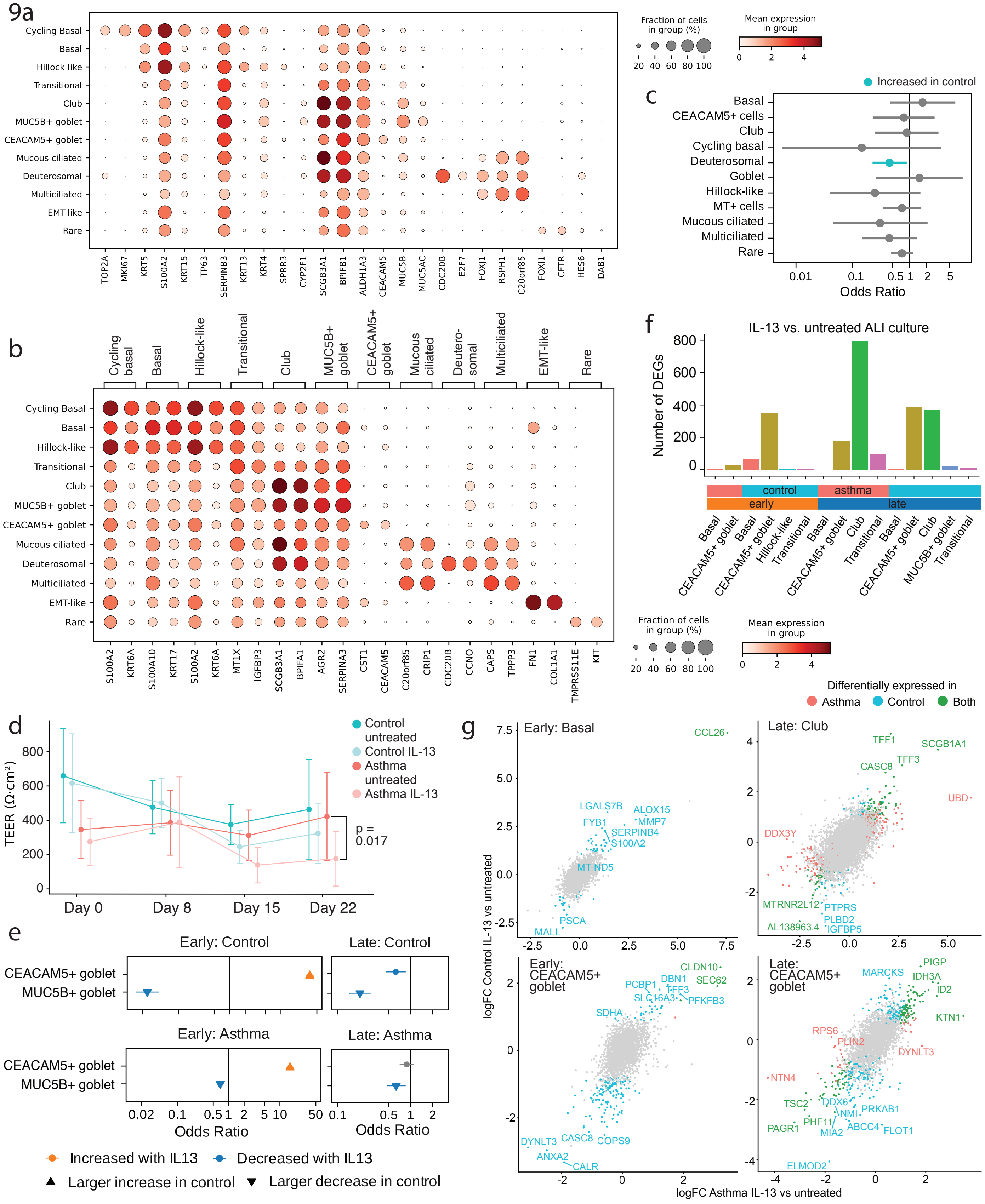


**Extended Figure 9. Gene expression in primary bronchial epithelial cells (PBECs) from patients with asthma and healthy controls in air-liquid interface (ALI) cultures and stimulated with IL-13. a**) Expression of classic cell type marker genes, and **b)** the top 2 cell-type-specific genes per cell type as annotated in the ALI cultured PBECs. **c)** Odds ratios of the abundance of cell types in untreated ALI cultured PBECs from patients with asthma compared to healthy controls (logistic mixed effects model at the single cell level). **d)** Transepithelial electrical resistance (TEER) of ALI cultured PBECs from patients with asthma and healthy controls with and without IL-13. **e)** Odds ratios of the abundance of the two goblet-cell subsets in ALI cultured PBECS from patients with asthma and healthy controls at the early and the late harvest timepoint with compared to without IL-13 stimulation (logistic mixed effects model at the single cell level). **f)** Number of significantly differentially expressed genes between IL-13-stimulated and control treated PBEC cultures per cell type, disease group, and time point. **g)** Log10(fold-change) of differential gene expression induced by IL-13 compared to untreated in patients with asthma versus the same in healthy controls, for basal and *CEACAM5+* goblet cells (early timepoint) and club and *CEACAM5+* goblet cells (late timepoint).
