## Supplementary Figures for "A repair-associated bronchial epithelial differentiation trajectory through *KRT14*+ basal and hillock-like cells drives airway inflammation and remodelling in childhood-onset asthma"

1

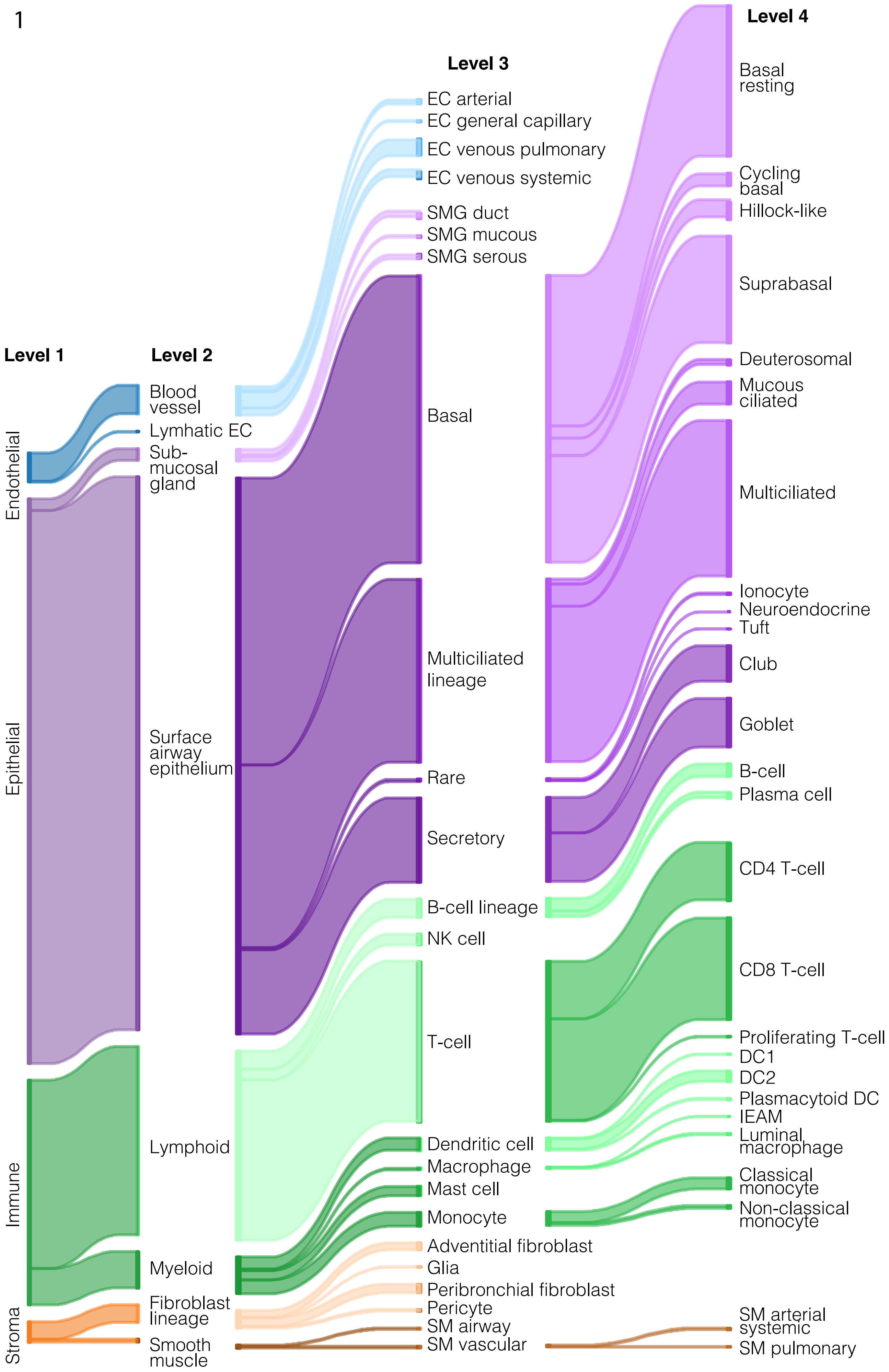


**Supplemental Figure 1. Hierarchical cell type annotation in the asthma cell atlas.** Sankey plot for the 4 levels of hierarchical cell type annotation in the asthma cell atlas, with most high resolution annotations for endothelial cells at level 3, for immune and stromal cells at level 3 and 4, and for epithelial cells at level 4.

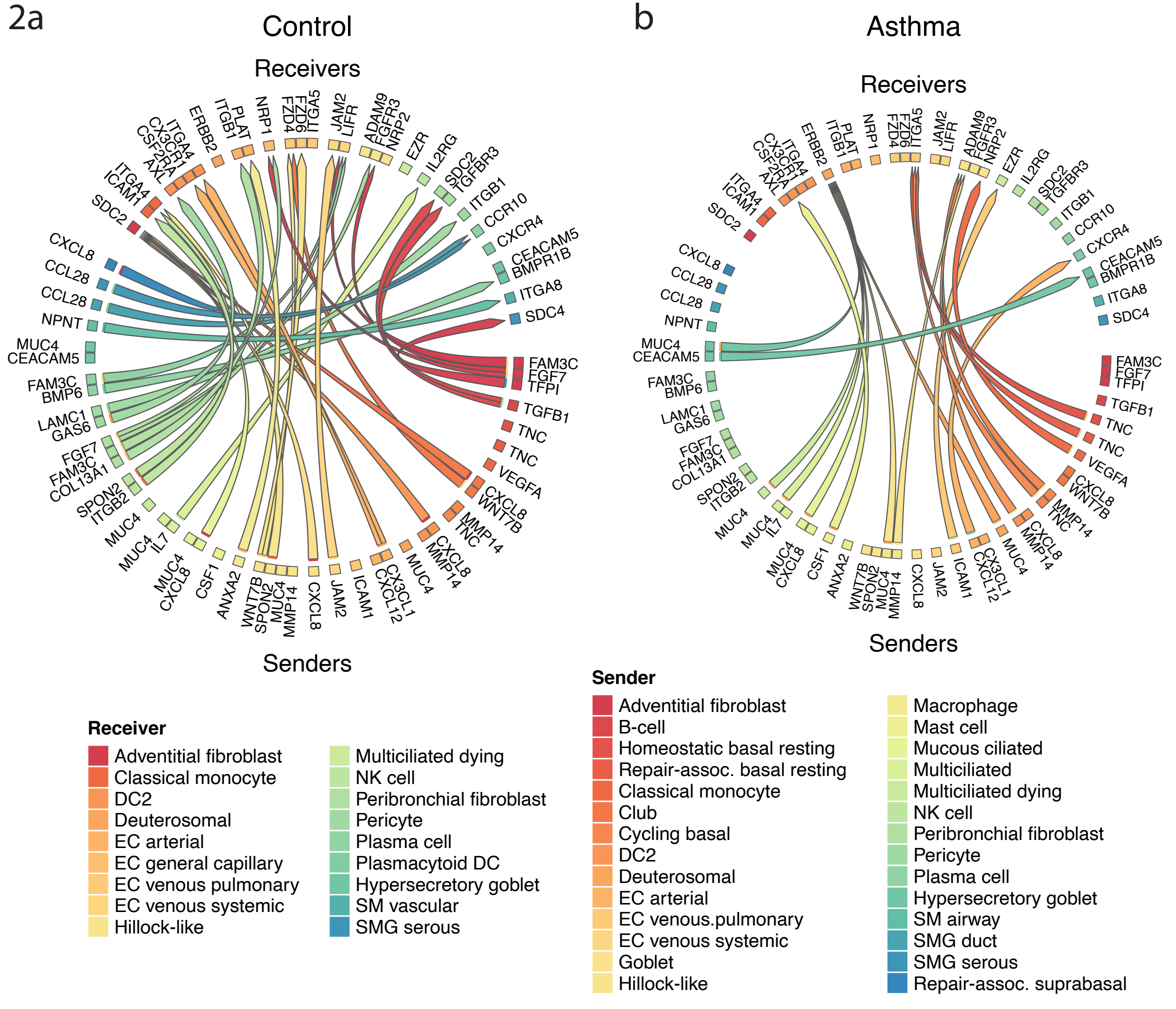


**Supplemental Figure 2. Cell-cell communication patterns identified by MultiNicheNet analysis. a)** Top 50 cell-cell interactions from MultiNicheNet analysis on the full asthma cell in healthy controls, and **b)** in patients with asthma.

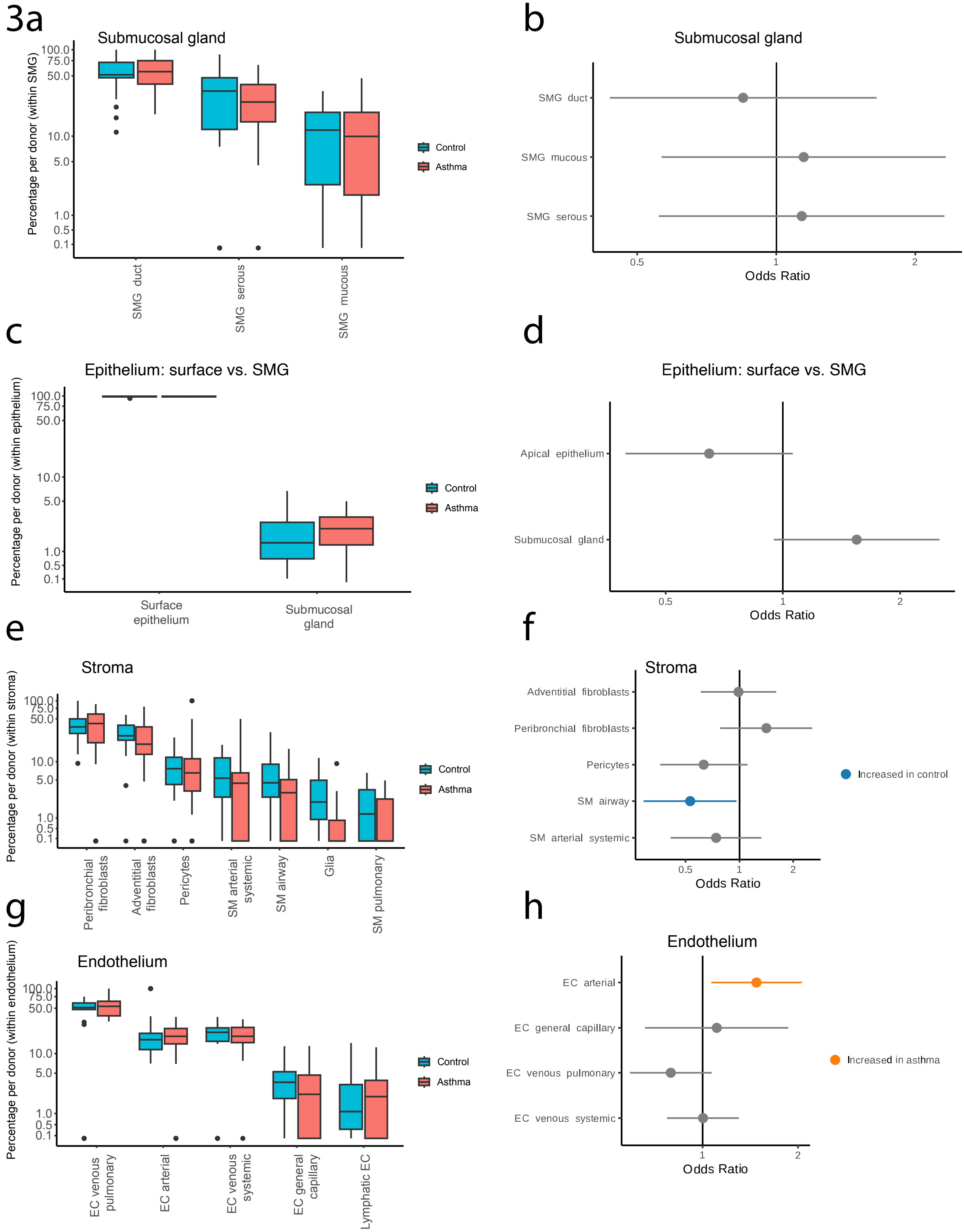


**Supplemental Figure 3. Abundance of submucosal gland, stromal and endothelial cells in the asthma cell atlas. a)** Percentage of epithelial cell subsets of the submucosal glands (SMG) within total SMG epithelial cells per donor in the asthma cell atlas. **b)** Odds ratios of the abundance of SMG cell subsets in patients asthma compared to healthy control (logistic mixed effects model at the single cell level). Panels **c)** and **d)** show the same percentage and odds ratios for the surface epithelium and SMG cells within the epithelium; **e)** and **f)** for stromal cell types; and **g)** and **h)** for endothelial cell types.

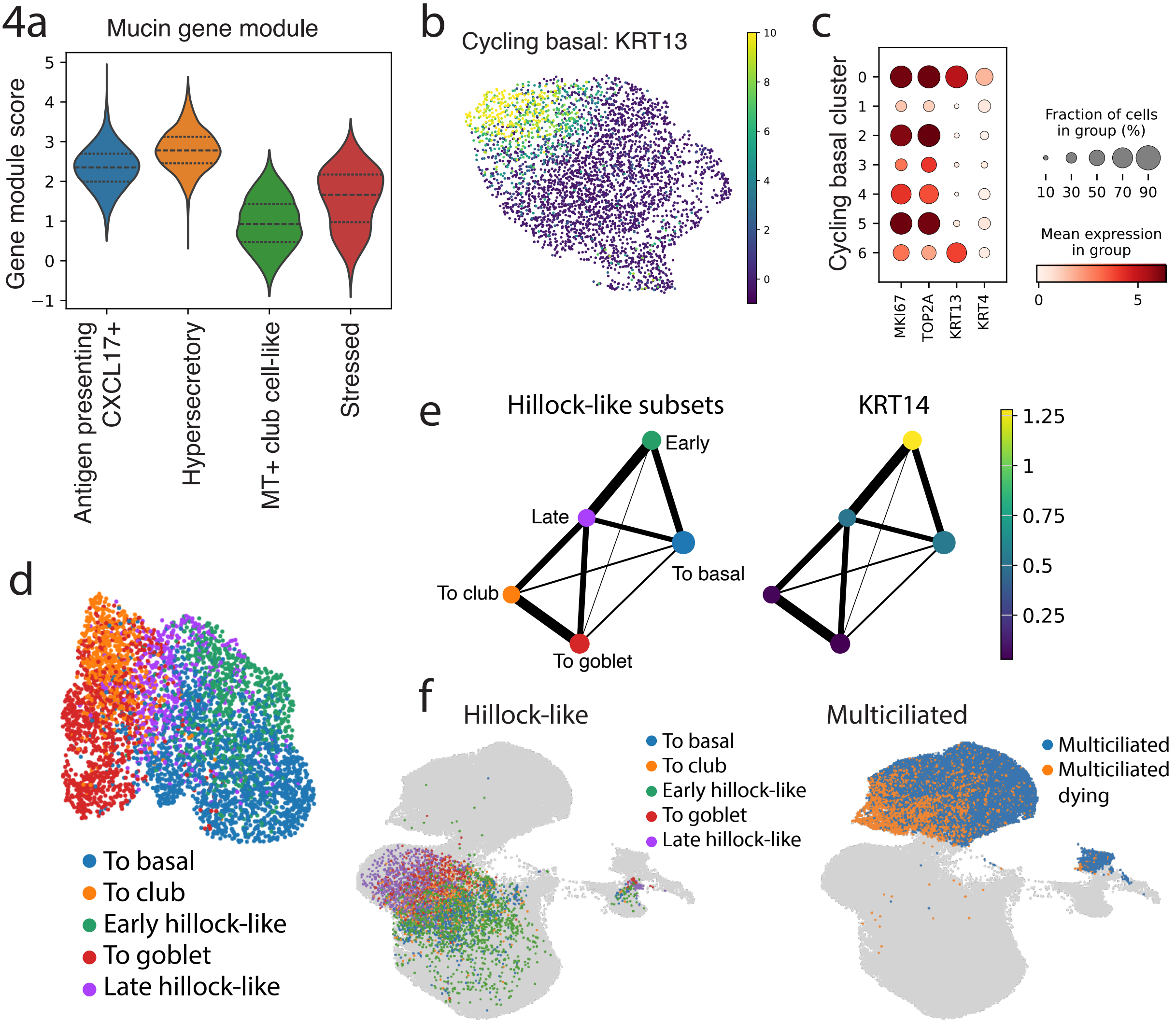


**Supplemental Figure 4. a)** Gene module score for the mucin production DRVI latent factor in the goblet cell subsets. **b)** UMAP of *KRT13* expression within the cycling basal cell subset of surface airway epithelial cells of the asthma cell atlas, and c) gene expression of *MKI67, TOP2A, KRT13* and *KRT4* in clusters of the same subset. **d)** UMAP of the hillock-like cell subsets within the hillock-like cell subset. **e)** PAGA graph of the hillock-like cell subsets coloured by cell subset label and *KRT14* expression. **f)** UMAP of the hillock-like cell subsets in the total population of surface airway epithelial cells (left) and dying multiciliated versus all other multiciliated cells (right).

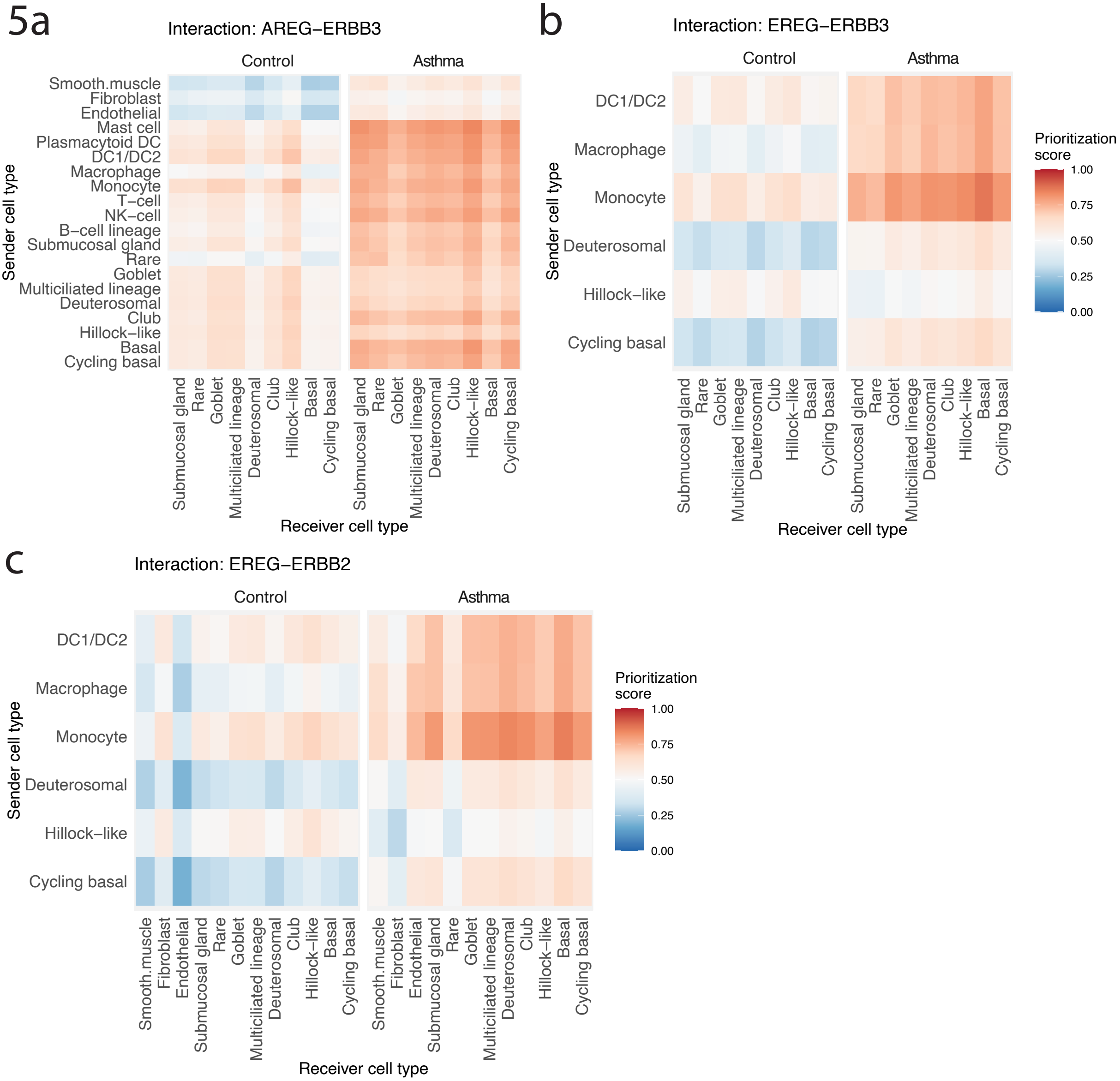


**Supplemental Figure 5. Cell-cell communication involving EGFR family genes and their ligands towards the airway epithelium in the asthma cell atlas. a)** MultiNicheNet prioritization score for *AREG-ERBB3* cell-cell communication between structural and immune cell types as senders (y-axis) and epithelial cell types as receivers (x-axis) in healthy controls and patients with asthma. Panels **b)** and **c)** show the same for *EREG-ERBB3* and *EREG-ERBB2* interactions, respectively.

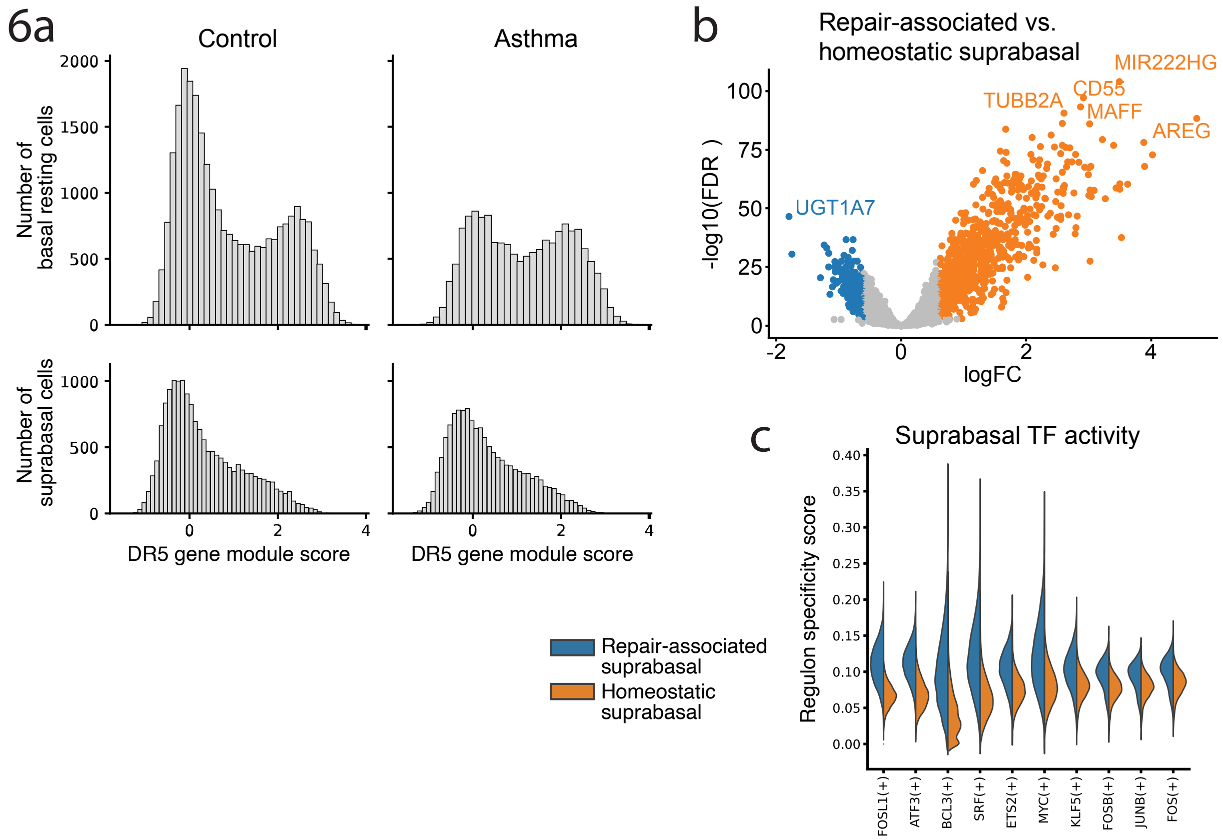


**Supplemental Figure 6. Repair-associated suprabasal cell proportions and gene expression in the asthma cell atlas. a)** Distribution of the expression of the DR5 latent factor in basal resting or suprabasal cells from healthy controls or patients with asthma. **b)** Log10(fold-change) versus -log10(F-DR-corrected p-value) of the differential gene expression between repair-associated and homeostatic suprabasal cells, corrected for disease status. **c)** Transcription factor activity score in repair-associated and homeostatic suprabasal cells.

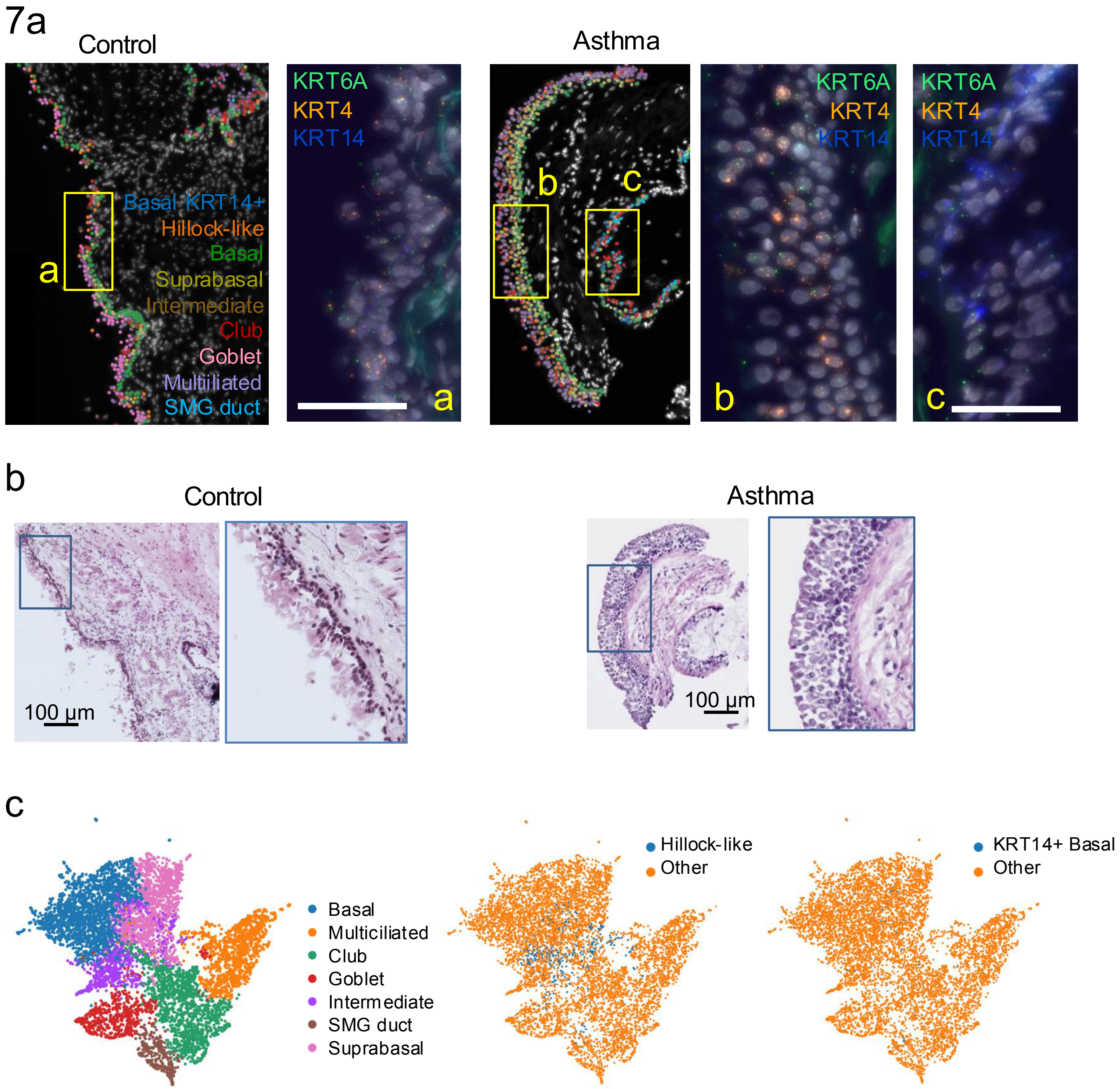


**Supplemental figure 7. SCRINSHOT spatial transcriptomic characterization of hillock-like and *KRT14+* epithelial populations in bronchial biopsies from patients with asthma and controls. a)** Cell type maps from main figure 4A with high-magnification views of gene expression from healthy and asthmatic biopsies. Nuclei (DAPI-stained, grey), *KRT6A* (green dots), *KRT4* (orange dots), and *KRT14* (blue dots) are presented on microscopic images, each dot representing a detected transcript. Scale bar: 50 $\mu$m. **b)** Histological images from serial sections corresponding to regions shown in a with high magnification areas of airway epithelium, demonstrating pseudostratified epithelium without visible squamous cell morphology. Scale bar: 100 $\mu$m. **c)** UMAP of spatially detected epithelial cell types, including cluster-annotated general labels, and manually annotated hillock-like and *KRT14+* basal cells. Positivity annotation criteria for hillock-like cells was defined by the expression of any of the following markers: *KRT4, SPRR3, KRT6A, KRT13* (≥ 2 transcripts), and negativity for all of the following: *SCGB1A1, SCGB3A1, LTF, CAPS, KRT15, KRT5, MUC5AC, LCN2* (< 5 transcripts). *KRT14+* basal cells were selected by positivity for *KRT14* (≥ 2 transcripts), and negativity for all of the following: *SCGB1A1, SCGB3A1, LTF, CAPS, KRT15, KRT5, MUC5AC, LCN2* (< 8 transcripts).

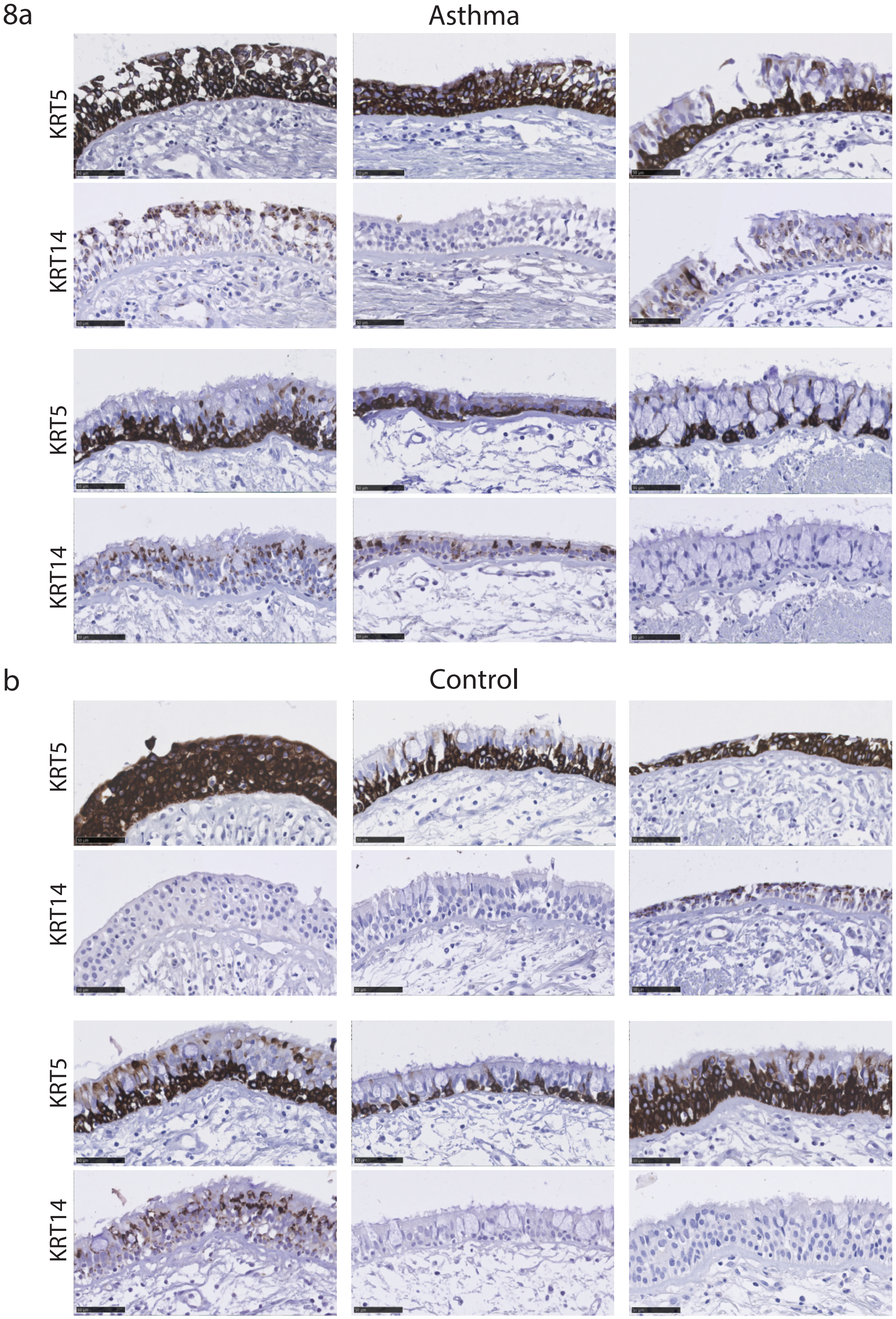


**Supplemental figure 8. Immunohistochemical staining for *KRT5* and *KRT14* in bronchial biopsies from patients with asthma and controls. a)** *KRT5* (upper panels) and *KRT14* (lower panels) stainings of the airway epithelium using 4 uM section of formalin-fixed paraffin embedded (FFPE) bronchial biopsies from patients with childhood-onset asthma of the ARMS cohort. Representative paired images spanning at least 200 uM of intact airway epithelium are shown from 6 independent donors. Scale bar indicates 50 uM. Panel b) shows the same for 6 independent healthy control donors of the ARMS cohort.

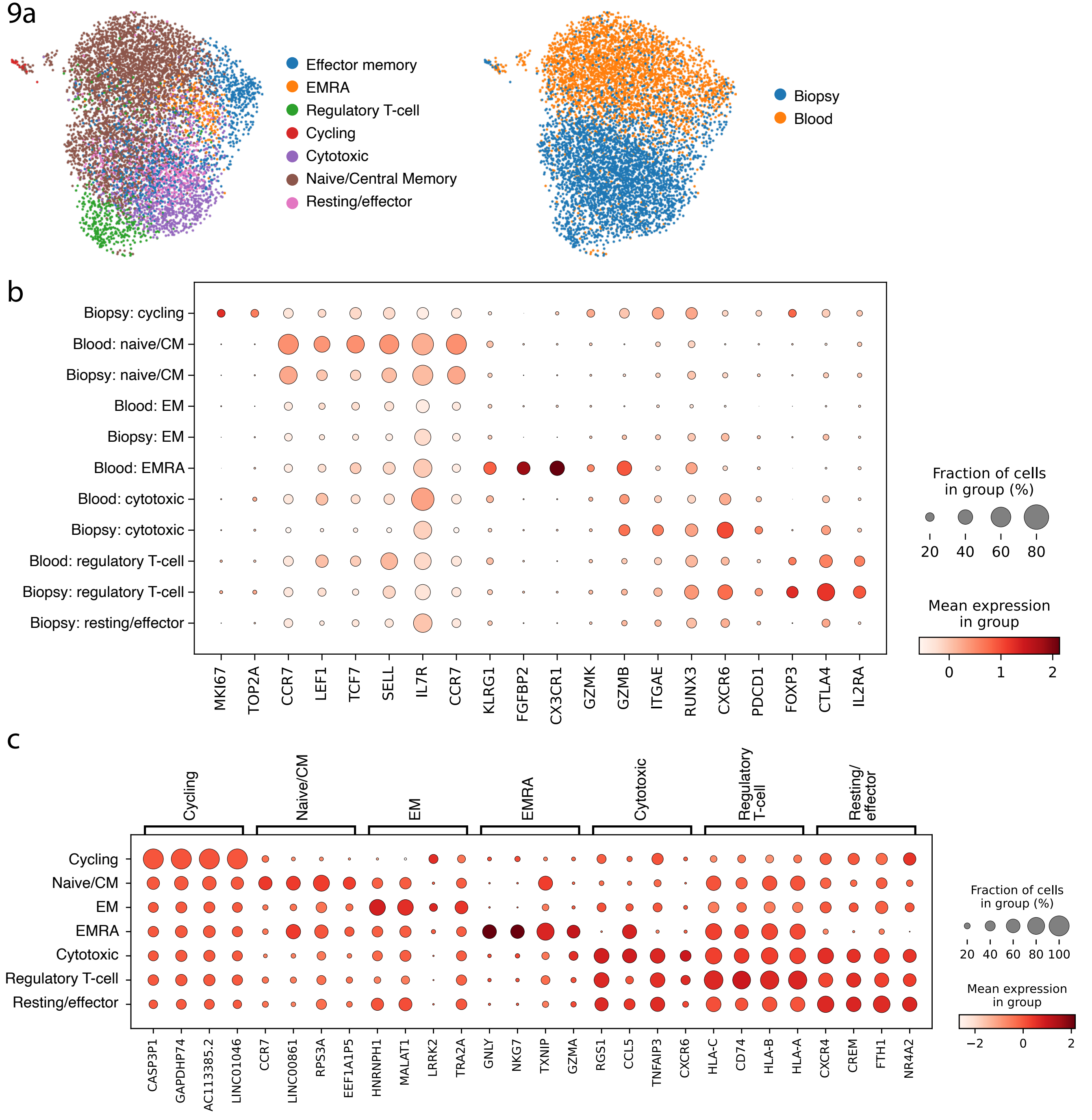


**Supplemental figure 9. Gene expression in CD4+ T-cells in blood and bronchial biopsies from patients with asthma and healthy controls. a)** UMAPs of FACS sorted CD4+ T-cells obtained from blood and bronchial biopsies from patients with asthma and healthy controls, showing cell type and tissue of origin. **b)** Expression of classic cell type marker genes, and **c)** the top 4 cell-type-specific genes per cell type as annotated in the CD4+ T-cell dataset.
